# Climate change, weather, and geography shape seed mass variation and decline across western North America

**DOI:** 10.1101/2024.10.23.619734

**Authors:** Leah J. Lenzo, Matthew L. Forister, Peggy Olwell, Elizabeth A. Leger

## Abstract

Seeds are an essential reproductive strategy for most plants, and seed mass is predicted to respond to environmental factors, but we do not know if climate change-related size changes observed in other organisms will also be reflected in seed mass. We investigated the temporal, geographic, and environmental patterns that influence seed mass across 2,800 species of native plants from over 13,000 populations across the western US using information from two decades of collections from a national program, Seeds of Success. Most species exhibited low variation in seed mass, though some varied by up to 220% over their occupied area. Measures of climate change influenced seed mass in over 40% of species, which could be evidence of adaptation to new optimal values or maladaptive responses. Species with the most variation in seed mass were those collected across sites with a wide range of minimum temperatures, from more northern latitudes, and across a wide range of dates, and seeds were generally larger in areas where it was wetter and warmer. Seed mass declined for over half (55%) of species, and declined overall by an average of 0.012 mg each year, though there were many species-specific differences, including 17% of species increasing appreciably in mass. Relationships between seed mass and environmental variables varied widely in strength, identity, and direction among species, including those that often co-occur. These results support the need to continue native seed collections for conservation and restoration, as plant reproductive strategies may be shifting with changing climates.

## Introduction

Seed production is an essential component of fitness for the majority of plants. Because adult plants are immobile, barring long distance dispersal, their reproductive strategies must adapt to local environments in order to persist (Clausen et al. 1940; Leimu and Fischer 2008; Baughman et al. 2019). Seed mass can strongly influence plant fitness during the seed-to-seedling transition, a crucial life history stage (Moles and Westoby 2004; Norden et al. 2009; Larson et al. 2015; Leger et al. 2019). Natural selection should favor an optimal seed mass, balancing seed mass with seed number, with local optima determined by interacting selection pressures including competition, dispersal, and herbivores (Venable 1992; Muller-Landau 2010; Seale and Nakayama 2020; Tuthill et al. 2023). Yet, seed mass varies greatly within and among individuals (Völler et al. 2012; Liu et al. 2013) possibly due to fitness benefits of variation in seed mass such as reducing competition among siblings or bet hedging in unpredictable landscapes (McGinley et al. 1987; Westoby et al. 1996). Variation in environmental sensitivity of seed mass could lead to the loss of some species in response to climate change if there is a mismatch between local optima and expressed traits, or increasing dominance of highly variable or adaptable species. In the era of climate change, there is evidence of shifting body size across taxa, with more species declining than increasing (Martins et al. 2023). Here we seek to understand how environmental factors may be influencing seed mass, which is critical for predicting species persistence (Cochrane et al. 2015; Tangney et al. 2020).

Seed traits vary across geographic, temporal, and phylogenetic scales (Westoby et al. 1992; Moles et al. 2007; Gorden et al. 2016). Larger seeds are typically more common in resource-poor environments where maternal provisioning is necessary for survival, and the opposite in resource-rich environments, where many small seeds may increase fitness (Smith and Fretwell 1974; Adler et al. 2013). At the largest spatial scale, environment affects seed mass across latitudinal gradients (Moles et al. 2007), with larger seeded species found closer to the equator. Hypotheses for this pattern include correlations with plant growth form (greater dominance of trees in the tropics), differences in seed dispersal strategies (e.g. wind vs. animal dispersal), and net primary productivity (greater towards the equator) (Murray et al. 2004). Latitudinal gradients in seed mass are present but weaker in North America (Liu et al. 2013; Gorden et al. 2016), possibly due to topographic variability (Malanson et al. 2007; Albrecht and McCarthy 2009). Further, these broad gradients may result from taxonomic turnover across latitudes but may not explain intraspecific, among-population variation in seed mass (Hulshof et al. 2013). It is also possible that within a community, species may produce different seed sizes in response to the same environmental conditions, which would be predicted if seed size plays a role in niche partitioning and community assembly (Turnbull et al. 1999; Leishman 2001; Muller-Landau 2010). Understanding the geographic, temporal, and taxonomic patterns of seed mass will provide valuable insight into evolutionary pressures experienced by native plants, including ongoing climate change (Walck et al. 2011, Frenne et al. 2013; Cochrane et al. 2015).

As climates change, many plant traits may change in response, resulting in cascading effects on seeds and reproductive strategies. For example, anthropogenic climate change can alter optimal phenological traits such as flowering time, with plants responding to warming temperatures by flowering earlier in the season (Bertin 2008; Burgess et al. 2007, Anderson et al. 2012). This can affect seed mass: plant genetic lines that were artificially selected for earlier flowering phenology produced fewer and larger seeds (Burgess et al. 2007) and globally, plant species have been shown to produce larger seeds under warming conditions (Zi et al. 2023). In addition to temperature, increasing CO2 levels are associated with changes in seed traits including number, mass, and germinability (Thürig et al. 2003, Hovenden et al. 2008). Thus, climate change and warming temperatures may result in selection for genotypes that produce fewer and larger seeds, and species that already have these characteristics, or are able to evolve them, may become more dominant. Alternately, resource stress can result in diminished seed mass (e.g. Suárez-Vidal et al. 2017), which could reduce plant establishment under climate change. Thus, understanding directional change in seed mass and the causes of seed mass variation has implications for our ability to mitigate the effects of climate change.

Biologists have characterized variation in seed characteristics for centuries, yet only recently do we have the fine-scaled resolution and longitudinal data to closely link seed traits with maternal environmental conditions (Kattge et al. 2011; Liu et al. 2019), which may help predict seed responses to climate change (Walck et al. 2011). In North America, we have an opportunity to examine the effects of geography and environment on seed mass through Seeds of Success (hereafter “SOS” see: blm.gov/sos Fig. 1), which is the national native seed collection effort in the United States led by the Bureau of Land Management. Through collaborative efforts and labor of thousands of field interns and seed processing technicians, SOS has made over 27,000 native seed collections across the United States since 2001 (Barga et al. 2020) (Fig. 2*A*). The mission of SOS is to collect wildland native seed for development, conservation, and restoration, and target species include restoration priorities as well as taxa not previously collected. Therefore, the dataset includes multiple accessions of the same species over multiple years from many locations, from a broad range of taxa, collected using a consistent protocol. The scale and scope of this effort has resulted in an invaluable source of seed for conservation and restoration across the US, and has generated an unparalleled wealth of information about the properties of seed in wild plant populations that can be used to inform ecological theory, global change biology, and restoration practices (Siefert et al. 2015; Moran et al. 2016).

**Figure 1.** Photos depicting seeds and collecting crews of the United States national native seed collection program, Seeds of Success (SOS). Clockwise, starting at bottom left: Seeds of Achnatherum thurberianum collected by BLM crew NV060, 2022; Avery Sigarroa, US Fish and Wildlife Service crew, NV0800 Nevada 2023; inflorescence of Bouteloua barbata, BLM crew NV052, Nevada 2022; soil color analysis, Nevada FWS crew NV0800, 2023; Allium acumunatum seed collected by Nevada BLM crew NV060, 2022. Yucca brevifolia var. jaegeriana seed collection by BLM crew NV052, 2020; voucher collection, Brian Cox, BLM Utah crew DUT020, 2024. All photos courtesy USFWS and Bureau of Land Management. Our estimates for cost per collection range from $5000-$10,000. Thus, this dataset stems from a $65M-$130M effort to fund seed collection crews, seed processing and cleaning technicians, seed storage, and the expertise and knowledge of botanists and ecologists.

**Figure 2.**
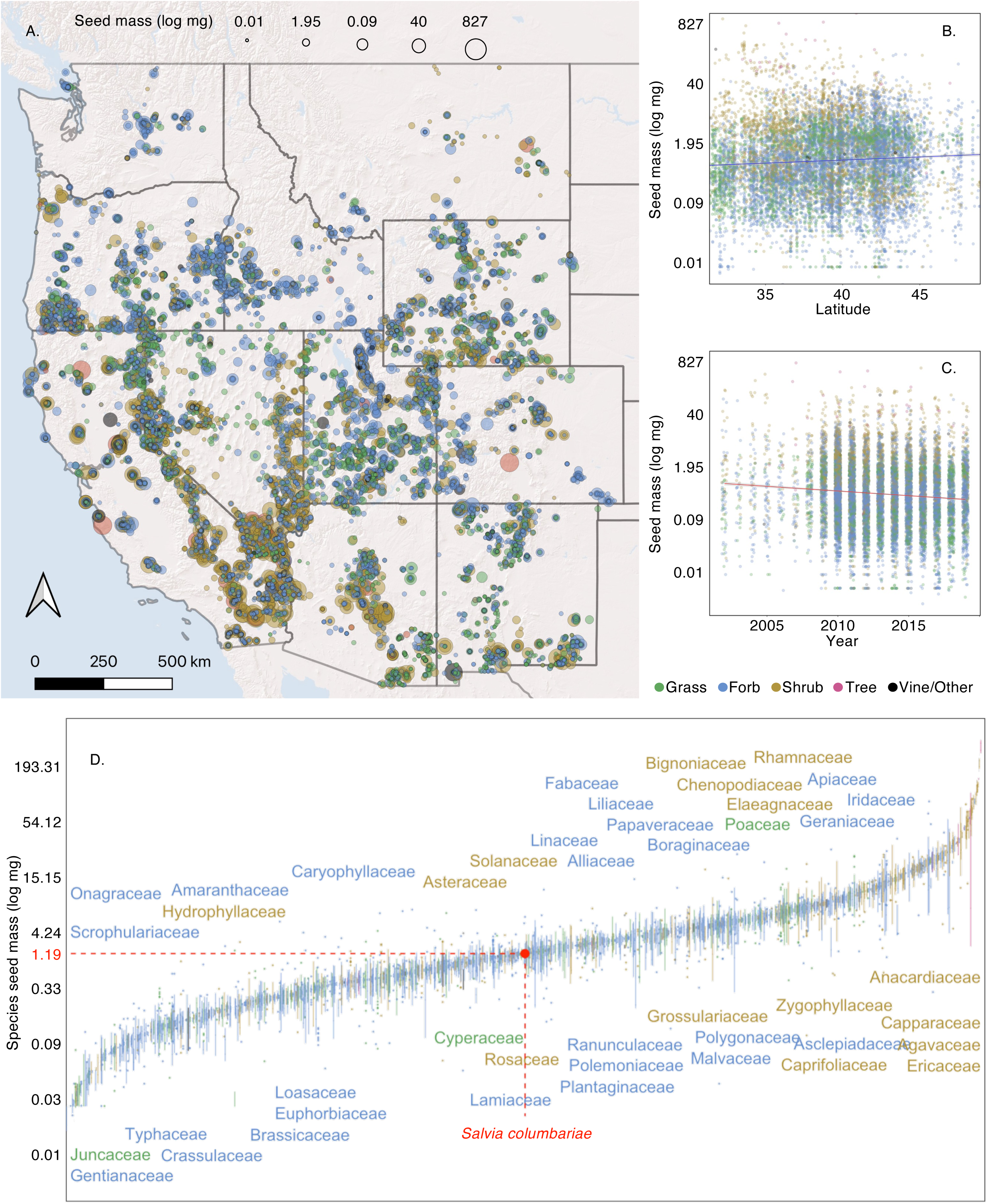
Seed mass of over 2,000 native plant species colored by growth form. A) Collection locations of 13,004 filled seed lots from Seeds of Success. Each point is a site sampled by a field crew after scouting and careful phenology tracking, representing a collection of at least 10,000 seeds from at least 50 plants. Seed mass was log-transformed for analysis, and sized by 20 quantile groups (5 examples shown at top). Log transformed seed mass of 13,004 populations plotted along B) latitude and C) year with 5 log breaks indicated on the y axis, lines represent simple linear regression of seed mass and latitude or year on the 3 collection dataset (see Q2, Q3 and Table S2, S3 for full generalized linear model and full spatial simultaneous autoregressive error model results, partial effects est = 0.0266, p < 0.0001; est = −0.010, p = 0.007 respectively). D) Log-transformed seed mass for 2093 species from 108 plant families from a dataset of seed collections across 11 western US states. For each species, the interquartile range of seed mass is shown by the lines and outliers are indicated as points and are colored by growth form. The y axis is on a log 10 scale in seed mass and species are ordered along the x axis by mean seed mass; the subset of families represented in the 10 sample dataset are listed near the first time the largest species appears in the distribution. *Salvia columbariae* is labeled as the median species, for reference. Seed mass was generated from the US Forest Service’s Bend Seed Extractory (BSE) processing of the Bureau of Land Management’s collections (BSE weighs a subset of 500 seeds per collection) and as part of the Seeds of Success program, which has been collecting native seeds since 2000.

To assess patterns of seed mass variation within and among species, we first took a broad approach, maximizing the inclusion of data to identify patterns of seed mass variation at the largest possible scale, asking 1) where do plants produce bigger seeds, and is there evidence of widespread changes in seed mass over time. Then, we investigated environmental influence on seed mass for more well-sampled species, both as a group and individually, asking 2) if there are life history traits or environmental, climate, or climate change variables that have a consistent influence on seed mass across and within species. Finally, we asked 3) if environment and geography predict not only seed mass but also seed mass variation across the range of collections, including a phylogenetic perspective.

## Methods

### Seed mass from Seeds of Success

SOS collecting teams follow a consistent seed collection protocol, collecting from plants of all sizes, aiming for a representative population sample. Teams identify plant populations that are large enough to collect at least 10,000 seeds on any given day of collection without sampling more than 20% of the available seed. For most species, mating system information (whether they are self-compatible or exclusively outcrossing) is unknown. Therefore, to attempt to capture at least one copy of 95% of alleles occurring within the population, teams collect from at least 50 individual plants, and often more, and teams are instructed to be sure to collect from small plants, not just the largest individuals (Brown et al. 1995). Teams also record the GPS coordinate of the center of the plant population. The majority of the SOS seed collections are made in the western continental US, and the vast majority of those are processed at the USDA Forest Service Seed Extractory in Bend, Oregon (BSE). Seed mass is assessed at BSE by weighing 5 subsamples of 100 seeds from each collection and BSE takes X-ray images of sample seeds to determine seed fill (presence or absence of embryo). We reduced the dataset by removing any seed lots with collection or packaging errors such as the presence of mold (poor drying time) or immature seed (poor collecting time). Collecting teams are required to do a ‘cut test’ to determine viability of seed on each day of collection, which involves slicing multiple seeds open to determine presence of a healthy embryo and endosperm, and thus collections should primarily be filled seeds. However, for certain species, this is difficult due to size of seed or lack of research on what constitutes a visibly healthy seed. Therefore, teams may occasionally and unintentionally make collections with low viability. When analyzing seed mass, we did not include 322 seed lots with 0% fill, instead focusing on asking how seed mass (including embryo and endosperm) vary rather than where and when populations produce viable seed. After merging BSE and national SOS datasets, we retained 13,004 unique accessions with seed mass, collection date, and location information (Fig. 2*A*).

For species’ name corrections, we referenced USDA Plants (https://plants.usda.gov/), and Tropicos (http://www.tropicos.org/) (accessed: Dec 7, 2020). After cross checking with a global phylogeny of angiosperm plants (Janssens et al. 2020), the taxonomy of 2,861 collections was accepted and 1,713 were corrected with updated names. We also gathered information on plant traits from the USDA Plants database, including growth form (we simplified to five growth forms: shrubs, grasses, trees, forbs, and vines, combining less represented categories, for example, shrubs and sub-shrubs.

Initially, we considered whether this dataset could be used to assess relationships between seed size and seed number, thus encompassing the expected trade-off between these aspects of natural history. However, an association between seed mass and clean weight was not evident (the average correlation across taxa was 0.15), and thus we do not include this analysis here (Supporting Information Data S1).

### Environmental data

We explored a variety of environmental covariates including, but not limited to, 30-year climate normals (1981-2010), annual and monthly temperatures, annual and monthly precipitation, silt content, sand content, folded aspect, soil at water capacity, day length, melt factor, saturation vapor pressure at mean monthly temperature, potential evapotranspiration, soil moisture of the previous month, actual evapotranspiration. Measures of climate change were estimated by extracting the mean daily minimum and maximum temperatures at each site in the spring (March-May) and fall (September-November) over 30 years prior to each collection date and finding the slope of each regression line against year, annual weather in the year of collection, and six month cumulative precipitation up to date of collection, all at 4k x 4k resolution from PRISM (PRISM Climate Group at Oregon State University). Additional climate and site variables were also obtained from the Climatic Water Deficit Toolbox and then run through an R script provided by Redmond (2019) to convert monthly normals into evapotranspiration estimates (McCune and Keon 2002; Lutz et al. 2010; Dilts et al. 2015; Redmond 2019). Soil data was obtained from SSURGO, GNATSGO as well as STATSGO2 using the soilDB package in R (Beaudette et al. 2022; Soil Survey Geographic Database (SSURGO)). To reduce the number of environmental covariates, we removed correlated variables (based on Pearson’s |r| 0.60) and selected year, day of year, and 14 environmental variables to move forward into analysis, including four measures of local climate change in temperature and precipitation (30 year changes in fall maximum and minimum temperatures, spring maximum temperatures, and annual precipitation), three annual weather variables (annual minimum temperature, cumulative precipitation 6 months prior to collection, and a composite of temperature and precipitation of the month prior to collection), slope, aspect, an estimate of heat load (Dilts et al. 2015), and four soil variables (clay content, an estimate of water content, soil pH, and organic content) (Beaudette et al. 2022) (Table S1).

### Statistical analyses

We first investigated spatial patterns of seed mass by using the full SOS dataset in generalized linear (GLM) and mixed (GLMM) models with log-transformed seed mass. Then, we subset the data to species collected from at least 3 populations to ask whether there were environmental influences on seed mass that could be estimated across a large species pool, with seed mass scaled within each species. Next, we reduced the data to species collected from at least 20 populations, and ran individual species models, with log-transformed seed mass, asking how the local environment influences seed mass for that subset of well-sampled species. Finally, we aggregated our data at the species level to ask if environment and geography predict seed mass variation for species that had been collected at least 10 times, including a phylogenetic perspective. Models were assessed with visual inspection of QQ plots and VIF scores.

#### Spatial distribution of seed mass

We mapped seed mass to visually assess spatial patterns separately for grasses, forbs, and shrubs, using the full dataset of 13,004 accessions. To predict seed mass across the western United States, we used inverse Distance Weighting (IDW) interpolation of the full dataset using the idw function in the gstat package in R (Pebesma and Graeler 2023). To test if there is a spatial gradient in seed mass among all species and populations in the western United States, we ran a GLM with log-transformed seed mass as the response variable using the full dataset with latitude, longitude and elevation as predictors; we also ran a second model that was identical but added growth form and duration as fixed effects. We next ran a GLMM with the same three spatial variables and response variable but also included plant family as a random factor to account for taxonomic variation. Not all species were collected from multiple populations, therefore we used family, rather than species, as a random intercept effect using the lmer function from the lme4 package in R (Bates et al. 2024). Similar to the GLM, we also ran a version of this model that included growth form and duration as fixed effects. After observing an effect of growth form in this model, we ran individual GLM and GLMM models for three highly collected growth forms to understand how seed-environment relationships varied among shrubs, grasses, and forbs.

#### Temporal patterns and environmental predictors of seed mass across species

To investigate whether there were consistent influences on seed mass across species, we analyzed relationships between seed mass and a series of predictors for all species that had a minimum of 3 collections in a spatial simultaneous autoregressive error model (SAR) using the errorsarlm function from the R package spatialreg (Bivand et al. 2024). This dataset contained 10,958 collections from 859 species from 73 plant families, and we scaled seed mass within each species. Prior to running the SAR, covariate selection was done using a random forest variable selection pipeline across all species, which implemented the Boruta function from the Boruta package in R (max runs = 100, p < 0.0001, Supporting Information Data S1) (Kursa and Rudnicki 2010), considering the suite of 14 environmental variables discussed above (Table S1), along with day of year, year, plant growth form and duration. In addition to the SAR, we ran a GLM with similar model structure to check for variance inflation factor for each independent variable, all of which were below 5. Each environmental variable was z transformed prior to analyses.

To ask whether seed mass has changed significantly over time across species, we ran individual GLMs for each species with seed mass scaled within species as the response variable and year as the predictor. We subset to species that had been collected from at least 3 unique years, resulting in 631 species, and saved the coefficient of year from all models. Next, we used the anscombe.test function from the moments package (Komsta and Novomestky 2022) to quantify kurtosis and test (with the Anscombe-Glynn test) for the possibility that the density of observations in the tails of our distribution is different from what would be expected given a normal distribution.

#### Temporal and environmental predictors of seed mass within well-sampled species

To understand how local environmental variables affect seed mass for each of the most well-sampled species, we ran individual SAR models for each species that was collected a minimum of 20 times (N = 111 species, N = 5196 total collections). We included the same set of 14 environmental variables (Table S1) and two temporal variables (day of year and year) as potential fixed effects: clay content, soil pH, soil organic matter, annual minimum temperature, previous six-month cumulative precipitation, change in fall maximum temperatures, change in spring maximum temperatures, change in precipitation, composite of previous month precipitation and temperature, available water capacity (AWC), heat load, slope, day of year, and year, as potential fixed effects. If a species had at least one confirmed important variable, these variables were included in a regression analysis. If a species did not have at least one confirmed important variable then the function TentativeRoughFix from the R package Boruta (Kursa and Rudnicki 2010) was used to confirm any tentative attributes, which were then moved into a regression. This function runs a simplified test comparing median feature z-scores to the most important shadow or simulated feature. For seven species (*Bothriochloa barbinodis, Castilleja linariifolia, Chaenactis douglasii, Eriogonum fasciculatum, Mimulus guttatus, Sarcobatus vermiculatus, Sphaeralcea munroana*), no variables of any kind were confirmed, and thus they were not analyzed but are included in our presentation of results. These species for which environmental variables could not be selected did not have any apparent differences in sample size or variation in seed mass relative to other taxa, but the lack of environmental predictors does raise the important issue of variation in sampling effort across all taxa.

The SOS dataset, while extensive, also has considerable variation among species in the number of populations sampled. Some species, particularly those of the greatest restoration need, have been collected from over 100 populations, whereas other species in the 20-collection dataset have only been collected from 20 populations. Therefore, to determine whether sampling effort affected our ability to estimate environment-seed mass relationships, we ran additional linear models on random subsamples of each species that has been collected from over 25 populations using the same environmental covariates as discussed in analyses above, with seed mass as the response variable. For each of those 77 species, we randomly sampled 20 populations 500 times with replacement and a minimum requirement of 15 unique populations, moved each random subset through a random forest model to determine important variables, and then ran a spatial error GLM on those populations. We set the 15 unique population threshold because models with fewer unique populations frequently failed to run or returned errors, likely due to insufficient variation in either seed mass or the spatial distance matrix. Coefficients were identified for each model for a total of up to 500 models (occasionally, no environmental variables were associated with seed traits of those 20 subsetted populations through the random forest). Model outputs were then saved, and compared to full species models to determine whether sampling effort may have affected individual species model results. We calculated quartiles for each covariate, tallied the number of subsampled models (out of 500) it appeared in and was significant at p < 0.05, summarized the number of times a subsampled model contained coefficients from the original model and how many times both the original model and subsampled coefficients were significant at p < 0.05.

#### Climate, environmental, and geographic patterns of seed mass variation (CV)

Finally, to understand the factors that affect the degree of variation in seed mass among species, we calculated the coefficient of variation (CV) of seed mass for each species that had been collected at least 10 times (Supporting Information Data S1). Not all SOS species have been collected with similar frequency, therefore we used an unbiased CV following standard protocols (Abdi et al. 2010; Barga et al. 2017) where CV is then weighted by sample size:

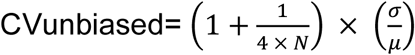

Values above 1 indicate greater variability whereas values less than 1 indicate low variation. We incorporated phylogenetic relatedness in these models using relationships described in Mishler et al. (2020). Seventy-nine species were not present in the phylogenetic tree even after correction and cross-checking taxonomy, thus, our final model for assessing variation in seed mass included 235 species from 6865 populations.

We aggregated environmental and climate variables across all collections for each species for a total of 33 environmental, spatial, temporal, and life history variables in addition to growth form and duration describing each of 235 species (Supporting Information Data S1, Table S1). Next, we ran all 33 variables through the random forest feature selection pipeline (described above and Supporting Information Data S1). Confirmed important attributes were then moved into a GLM with log-transformed unbiased CV as response to check VIF (all were below 5) and model diagnostics. Then, attributes were moved into a phylogenetic generalized least squares regression (PGLS) using Pagel’s correlation of trait evolution and the phylogeny from Mishler et al. (2020).

## Results

Seed mass ranged from 0.0046 mg (the average across-population mass of *Epipactis gigantea*) to 827.7 mg (for *Pinus sabiniana*) with a mean across species of 4.4 mg and median of 1.2 mg (Fig. 2D). Certain plant families, including Fabaceae and Ericaceae, consistently had larger seeds whereas others (e.g. Juncaceae and Onagraceae) exhibited smaller seeds (Fig. 3). Seed mass ranged from highly conserved across populations (e.g. CV of 0.0005 for *Wyethia magna*) to highly variable (CV of 2.2 for *Juncus occidentalis*), with a mean CV of 0.32 and median of 0.27 (Fig. 3).

**Figure 3.**
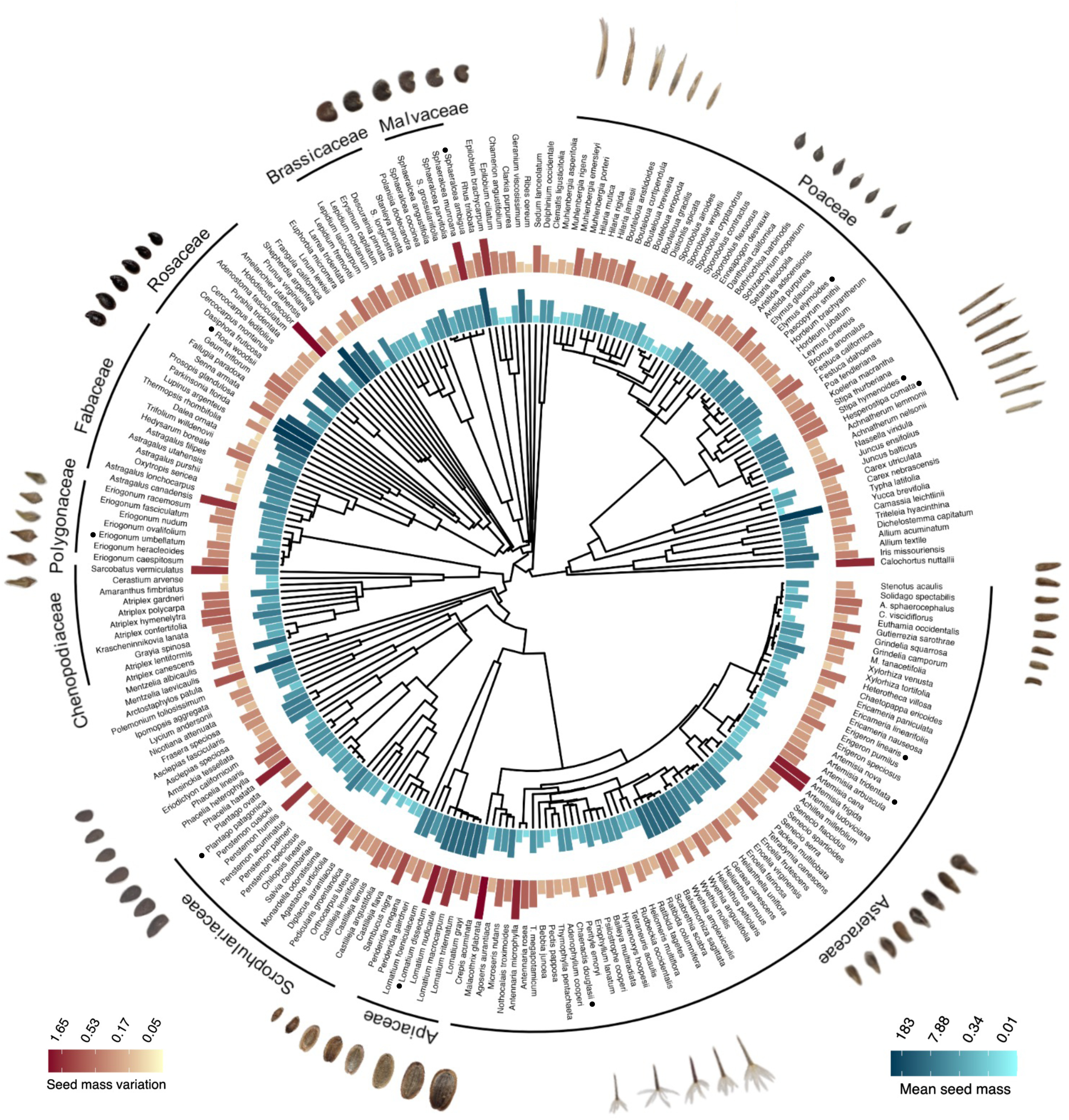
Phylogenetic tree of 235 plant species from 44 families collected from at least 10 populations using phylogenetic tree from Mishler et al. (2020). Inner bar chart represents log-transformed mean seed mass of each species, outer bar chart represents log-transformed coefficient of variation of seed mass for each species. Seed mass is from the US Forest Service’s Bend Seed Extractory processing the Bureau of Land Management’s Seeds of Success collections. Seed mass coefficient of variation is calculated as CVunbiased=(1+14×*N*)×CV. Images of native seeds are included; species names are indicated with a nearby bullet (photos: LL).

### Spatial distribution of seed mass

Most areas across the western US were dominated by smaller seeds with no obvious spatial segregation (Fig. 2*A*; Fig. S1). Latitude had a small, positive effect on seed mass in models that included latitude, longitude, and elevation as predictors (GLM: β = 0.027, p < 0.001) as well as models that included plant family as a random factor, which had a better model fit than geography alone (Supporting Information Data S1, GLMM: β = 0.031, p < 0.001; Fig. 2*B*, Table S2, Table S3). Seeds were, on average, 1.03 mg larger for every 1 degree north, in contrast to global patterns, though in models that considered seed mass by growth form, shrubs had a contrasting response to other plant growth forms, with smaller seeds in more northern latitudes (Table S2). Longitude had a small negative effect on seed mass (GLM: β =−0.005, p = 0.089, GLMM: β =−0.01, p < 0.001), with smaller seeds (0.989 mg smaller for every 1 degree) in more eastern areas.

### Temporal patterns and environmental predictors of seed mass across species

We examined relationships between seed mass, environment, and time for well collected species (10,958 collections from 859 species collected from at least 3 populations) and found that, overall, seed mass decreased by 0.012 mg each year (Fig. 2C, β = −0.010, SE = 0.004, p = 0.007). When considering species independently, 38% changed appreciably in seed mass, representing a distribution of change with greater density in the tails of the distribution (species getting larger or smaller) than would be expected from a normal distribution (kurtosis = 11.153, z = 10.596, p < 0.001). Of the 631 species that were collected from 3 unique populations and years, 55% were declining in seed mass over time, with 132 declining significantly and 107 increasing significantly (at p < 0.05) in seed mass.

Multiple environmental variables had consistent positive effects on seed mass across species (Fig. 4A). The strongest positive relationships were between seed mass and cumulative precipitation over the previous 6 months leading up to collection (β = 0.051, SE = 0.012, p < 0.0001), as well as annual minimum temperature (β = 0.044, SE = 0.011, p < 0.001), with generally greater seed mass in the relatively wetter and warmer collection sites (Fig. 4A, Table S4). Clay content and slope also had positive relationships with seed mass (β = 0.045, SE = 0.011, p < 0.001, and β = 0.045, SE = 0.011, p < 0.001 respectively), with generally larger seeds in areas with higher clay content and steeper slopes. Heat load also had a positive relationship with seed mass (β = 0.024, SE = 0.012, p = 0.038), with larger seeds in relatively warm collection sites. Neither growth form (shrub, grass, forb, etc.) nor duration (annual, perennial, etc.) were retained as important effects in the model selection process.

**Figure 4.**
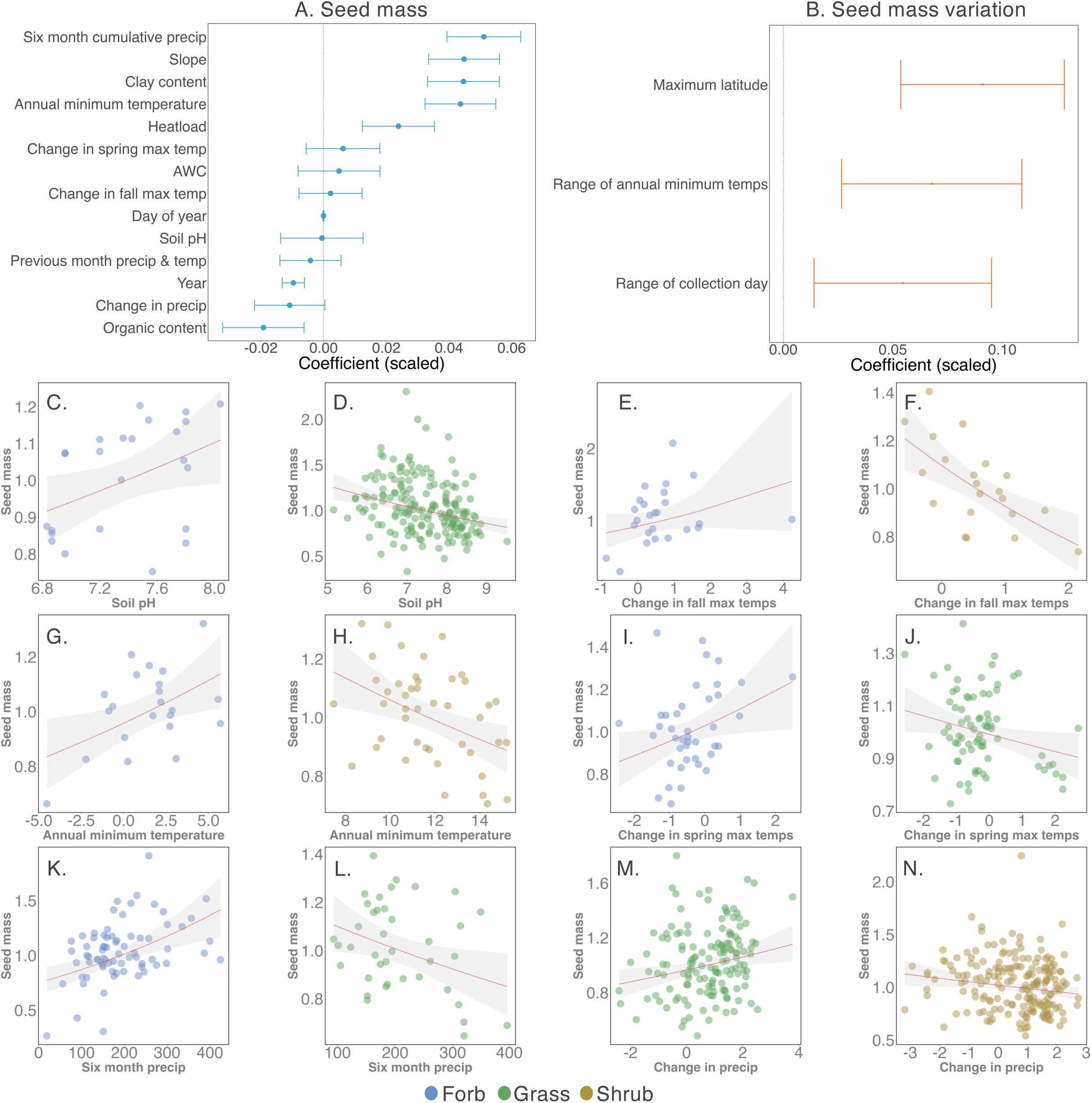
Relationships between seed mass and environment for native plant species from across the western US. Environmental influence on A) seed mass using a spatial simultaneous autoregressive error model (N = 859 species, 10,958 populations, 73 plant families in the 3 collection dataset) and B) the variability of seed mass estimated from a phylogenetic generalized least squares regression (N = of 235 species, 6865 total populations, in the 10 collection dataset). Covariates are centered and scaled, and error bars represent standard error produced by the PGLS. Relationships between soil pH and seed mass for C) *Balsamorhiza hookeri* and D) *Elymus elymoides*; relationships between change in fall maximum temperatures 30 years prior to each collection and seed mass for E) *Erigeron pumilus* and F) *Hymenoclea salsola*; relationships between annual minimum temperature and seed mass for G) *Asclepias speciosa* and H) *Plantago ovata*; relationships between change in spring maximum temperatures 30 years prior to each collection and seed mass for I) *Eriogonum heracleoides* and J) *Pseudoroegneria spicata*; relationships between cumulative precipitation six months leading up to collection and seed mass for K) *Heterotheca villosa* and L) *Achnatherum thurberianum*; relationships between change in annual precipitation 30 years prior to each collection and seed mass for M) *Hesperostipa comata* and N) *Artemisia tridentata*. For C-N, relationships were analyzed with a spatial simultaneous autoregressive error model (SAR) seed mass is log-transformed and predictor variables are centered and scaled; species specific results and full model structures are in Table S5. Lines represent the semi-partial regression plots between a single predictor variable and seed mass, points are colored by growth form.

### Temporal patterns and environmental predictors of seed mass across species

We found several common predictors of seed mass among well-sampled species (those with at least 20 collections each; 111 species, 5196 total collections), confirming previous results (Fig. 5, Table S5). Specifically, annual minimum temperature, clay content, heat load, cumulative precipitation six months prior to collection, soil pH, and day of year were statistically significant in at least 10 species models. However, the strength and direction of relationships varied widely among species (Fig. 4*C-N*; Fig. 5). For example, soil pH had a strong positive influence on seed mass of *Balsamorhiza hookeri*, increasing seed mass 1.095 mg as soil pH increased by 1 (Fig. 4*C*, Fig. 5), while *Elymus elymoides* had the opposite response, decreasing seed mass by 0.889 mg as soil pH increased by 1 (Fig 4*D*, Fig 5). We used a resampling approach to investigate the influence of variation in sample size (the number of populations) on species-specific results; results were robust to systematic reductions in the number of populations included in models (Table S6).

**Figure 5.**
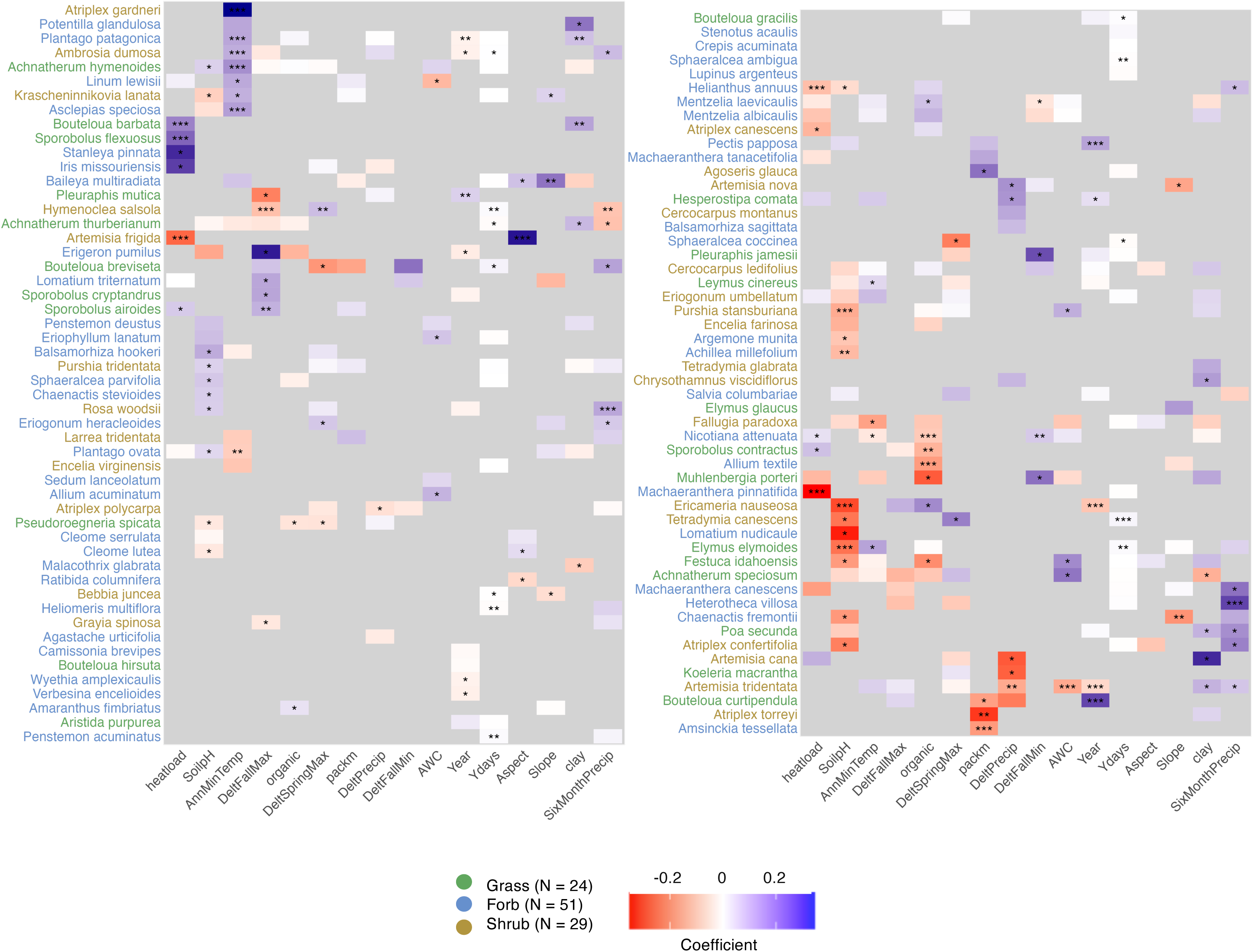
Environmental and climatic influence on seed mass estimated from 104 individual spatial simultaneous autoregressive error models for species collected from at least 20 different populations (5,010 total populations). Axes indicate individual species models as rows and possible covariates as columns; gray boxes indicate variables that did not enter final models. Red boxes indicate negative relationships, grading to blue for positive relationships. Significance is indicated with *** = P <0.0001, ** = P<0.001, and * = P<0.05. Species’ names are colored by growth form. Species and variables are organized by a similarity matrix based on model results. Figure is halved at *Penstemon acuminatus* continuing to *Stenotus acaulis* following the similarity matrix.

Multiple measures of climate change across a species collection range (including change in spring and fall maximum temperatures, change in precipitation, and change in fall minimum temperatures) were included as important predictors in 47 species models (out of a possible 111 species) and were significant 23 times in 22 models (Fig. 5). Of those, 11 had a statistically significant negative correlation between measures of climate change and seed mass and 12 climate change coefficients were positive, again illustrating heterogeneity of responses (Fig. 5, Table S5). *Atriplex polycarpa* and *Artemisia tridentata* were particularly responsive to climate change, including three measures of climate change in each species model, with change in precipitation significantly affecting seed mass (Table S5). When we ran individual species models testing whether seed mass has changed over time, we found that 132 species were getting smaller (p < 0.05), 107 species were getting bigger (p < 0.05), and 392 had not changed seed mass (p > 0.05).

We conducted a subsampling exercise to understand whether collection effort affected the identification of important variables. Within the subsampled models, the variables identified with the full sample were also among the most frequent variables in the subsampling for 48 of 78 species (62%) (Table S6). When a subsampled model contained a variable from the original full species model it fell within the original 95% confidence interval 65% of the time. Of those models that shared a variable, coefficients were significant in both the full species model and sample models 63% of the time. Overall, as one might expect, sampling effort had an effect on our ability to identify the most important predictors of seed mass, but many seed-environment relationships are robust to sampling effects.

#### Climate, environmental, and geographic patterns of seed mass variation (CV)

Our models thus far have assessed mean seed mass, but because natural selection acts on variation, we also quantified seed mass variation among populations of a single species to understand the potential for evolutionary change. Here, we aggregated our data at the species level to ask how environment and geography predict seed mass coefficient of variation while considering phylogenetic relationships among species. Species differed in the amount of variation (CV) in seed mass across collection locations and collection years (Fig. 3). Most species exhibited low variation in seed mass, but some, including *Artemisia frigida, Phacelia hastata, Chaenactis stevioides,* and *Sphaeralcea munroana*, exhibited CV values above 1 indicating higher variation. Considering the 235 species that had been collected at least 10 times and for which phylogenetic information was available, the strongest predictors of seed mass variation were the maximum latitude from which a species was collected (β = 0.091, SE = 0.037, p = <0.001), the range of annual minimum temperatures experienced across the collection area (β = 0.068, SE = 0.041, p = 0.101), and the range of collecting days (difference between first and last day of collection) (β = 0.055, SE = 0.041, p = 0.180; Fig. 4*B*, Table S7). Neither growth form nor duration were selected as important, and, when included in models with environmental covariates or alone in a simple model, neither predicted seed mass variation (results not shown).

## Discussion

Climate change continues to affect the morphology of individual species of plants and animals, but due to challenges of amassing quality datasets, effects across entire assemblages of species occupying large geographic areas are rarely investigated (Sheridan and Bickford 2011; Violle et al. 2012). Furthermore, there is increasing consensus about the importance of intraspecific variation for understanding ecological and evolutionary processes that affect species and community responses to climate change (Siefert et al. 2015). Yet, we often lack consistent population-level replication for specific fitness related traits, leading to uncertainties in identifying these intraspecific patterns (Moran et al. 2016). By assessing seed mass variation within and among species simultaneously, our results offer a unique perspective on plant reproductive biology and species coexistence, illuminating how environment and climate change affects propagule size in land plants.

We found spatial patterns and environmental predictors of seed mass across all species and common environmental predictors of seed mass within multiple species, including precipitation, heat load, and soil characteristics like pH. Seeds were generally larger in warmer and wetter areas, consistent with tree seed sizes observed in Boreal forests (Liu et al. 2013). However, we observed even stronger signals within species, though often in heterogeneous directions and magnitudes, confirming that intraspecific seed trait-environment relationships can be idiosyncratic, and differ from broad interspecific patterns (Gorden et al. 2016). Heterogeneity in response among species may be an important form of resilience, which can only be seen when analyzing traits at the scale of communities and entire floras. Conservation and restoration practices could benefit from considering intraspecific variation in addition to interspecific variation (Sgrò et al. 2011). While conservationists may prioritize protecting species that are more sensitive to climate and climate change, practitioners may benefit from planting species with seed traits that are less sensitive to climate change or selecting a mix of species that are sensitive to different factors, thereby incorporating natural resilience into restoration.

While seed mass was highly conserved for most species, others varied more across space and time, including species collected across broader minimum temperature gradients, higher latitudes, and those with longer collecting seasons. Given that warming minimum temperatures are an early consequence of anthropogenic climate change (Parry et al. 2007), with northern latitudes showing particular acute change (Bunn et al. 2007), these sensitivities suggest that climate change will be exerting selection pressure on seed mass in plants. Previous studies linking increased CO_2_ levels with larger seeds could lead to the expectation that seeds would be overall increasing in mass (Thürig et al. 2003, Hovenden et al. 2008). Instead, we observed overall declines in seed mass across the western US over the past 20 years, though some species increased in seed mass, consistent with heterogeneous changes observed in seed mass in alpine plants (Zhang et al. 2019). Given the importance of seed mass for plant regeneration (Moles and Westoby 2004) and the role of seeds as a food source for granivores especially in arid lands (Solbreck and Knape 2017; Dylewski et al. 2020), these small changes in seed mass could have cascading effects on ecosystems (Sheridan and Bickford 2011).

Further highlighting the impact of climate change on terrestrial plant seeds, we detected sensitivity to direct measures of temperature and precipitation changes for over 40% of well-collected species, though directionality was not consistent. For example, showy milkweed, *Asclepias speciosa*, an important host plant for the Monarch butterfly, makes larger seeds in warmer areas, while *Plantago ovata* shows the opposite pattern. Another very important species in the region, big sagebrush (*Artemisia tridentata),* was particularly responsive to climate change, with three of our four measures of climate change affecting seed mass. It is possible that seed mass sensitivity to climate change represents an adaptive response, or if changes are a result of resource limitation, it could indicate that these species are particularly vulnerable to climate change. Field studies of the relationship between seed mass and fitness in highly sensitive species identified here would be important to understand if these plants should be prioritized in conservation efforts.

We are particularly fascinated by the many cases where species had strong responses to the same variables, but in contrasting ways (Fig. 5). For example, soil pH had a strong positive relationship with seed mass in the widespread forb *Balsamorhiza hookeri*, while *Elymus elymoides*, a widely distributed perennial grass commonly used in ecological restoration had the opposite response. Other species did not change seed mass in response to any of the variables we considered, maintaining nearly constant seed mass across collection locations and years (e.g. *Chimaphila umbellata*). Some species varied in seed mass across space and time but not in response to any environmental variables we included here (e.g. *Chaenactis douglasii*). It may be that other ecological influences such as herbivory, competition, success of insect pollination, or polyploidy may have an effect on seed size in these species (Paige and Whitham 1987; Vaughton and Ramsey 1997; Chan et al. 2022). In the future, close monitoring of native plant communities may uncover these ecological effects on seed mass, and our results could inform comparative studies across species with contrasting levels of seed mass variation.

Overall, few species exhibited high (>1) coefficients of variation, supporting our understanding that seed mass is a highly conserved trait with some exceptions (Zhang et al. 2020), and it is possible that species that are more variable in seed mass may be poised to adapt to changing climates (Frei et al. 2014; Cochrane et al. 2015). However, species demonstrating changes in seed mass in response to climate change may be adjusting seed mass in maladaptive ways. For example, if smaller seeds confer greater dispersal capability, but a longer, warmer growing season results in larger seeds, this may reduce dispersal distances and lead to greater competition with siblings (Harper et al. 1970; Greene and Johnson 1993). In contrast, species that are demonstrating decreasing seed mass in response to warming temperatures may be experiencing resource limitation, and changes in seed mass may reduce progeny resources and stress tolerance (Westoby et al. 1996).

Understanding where and how species vary in seed mass can help elucidate the reproductive strategies of land plants, and understanding the consequences of climate change for seed characteristics is essential for informing applied science. Our work provides a foundation for multifaceted future investigations into how changes in seed mass can affect plant fitness and highlights the need to increase our population level monitoring and consider intraspecific variation in key traits. Our findings also support the need for continued collection of locally adapted native seeds from many sites for restoration through Seeds of Success (National Academies of Sciences, Engineering, and Medicine 2023), and the species-specific nature of seed mass responses to climate support targeting a wide range of collection sites to collect and preserve high quality seed for multiple species. Additionally, because the reproductive output of many native plant species in the western US is sensitive to climate change, the urgency to continue native seed collections for conservation is greater than ever.

## Supporting information

SupportingInformation

## Acknowledgements

This work was partially supported by U.S. Department of Interior Bureau of Land Management award L20AC00317 and L22AC00506 to E. A. L. We thank J. Allen, K. Badik, E. Hanan, A. Agneray, J. McClinton, T. Bartz, R. Frederick, L. Shriver, C. Silliman, S. Swim, A. Kim, B. Nagelson, C. Phelan, H. Noble, S. Bisbing, M. Lohman, and J. Brockman for input on questions, analysis, and data visualization. The Seeds of Success program and data would not be possible without the hard work of seed collection crews, seed processing and cleaning technicians, availability of steed storage, and the expertise and knowledge of botanists and ecologists. The authors thank all their federal, state, and nongovernmental agency partners whose efforts towards native plant community restoration and conservation in the U.S. created this rich dataset. MLF thanks the National Science Foundation, OIA-2019528, for support.

## Data Availability

The dataset will be available on Dryad.

