## SupportingInformation for "Climate change, weather, and geography shape seed mass variation and decline across western North America"

This file includes:

- Supplementary Text
- Figs. S1 and S2
- Tables S1 to S7
- References

**Supplementary Text**

The Seed Size, Seed Number Tradeoff (SSNT) is well established (Smith and Fretwell 1974; Venable 1992; Leishman 2001). Therefore, we assessed any correlation between average seed mass and total clean weight collected for each species collected from at least 3 populations. However, we did not find any strong correlations between seed mass and clean weight (mean correlation 0.15), especially as sample size increased. This may be because collecting teams aim to collect at least 20,000 seeds or as much as possible. Within our dataset, total seed collected may not be related to seed mass or influenced by biological effects but is more likely a function of collecting team efforts and time, and thus we do not include this analysis here. Additionally, we have data at the population level rather than the individual plant level which may not accurately capture the trade-off between seed mass and production

**Supplemental Methods**

**Statistical Analyses**

Boruta is a feature selection algorithm for random forest which aims to select the most important features using a criterion of robust predictive power. For computational efficiency, only variables with mean importance score above 10 were moved into the SAR after visually assessing the Boruta output plot and distribution of scores. We then ran a SAR with seed mass (scaled within each species) as a response variable in a model that included the following subset of variables identified through the Random Forest pipeline: clay content, soil pH, soil organic matter, annual minimum temperature, previous six month cumulative precipitation, change in fall maximum temperatures, change in spring maximum temperatures, change in precipitation, composite of previous month precipitation and temperature, available water capacity (AWC), heatload, slope, day of year, and year, as fixed effects, with latitude and longitude included in the neighbors list with spatial weights. Occasionally, multiple species were collected from the same location and spatial neighbors cannot have a distance of zero. Therefore, we slightly jittered GPS coordinates of completely overlapping collections by a random value between 0.0001 and 0.00001 degrees.

*Temporal and environmental predictors of seed mass within well-sampled species*

For each species, we quantified spatial autocorrelation in seed mass among populations using Moran's I and a permuted test of significance and by comparing Nagelkerke and McFadden pseduo-$R^2$ between spatial error models, linear regression models, and linear regression models with Moran's eigenvector maps as covariates. For the majority of species, seed mass was spatially autocorrelated and the spatial error model performed best. Therefore, all species were analyzed with the same SAR and variable selection method as Q2 (Bivand et al. 2024).

*Climate, environmental, and geographic patterns of seed mass variation (CV)*

To understand whether species-level characteristics predict variation in seed mass as estimated with CVunbiased, we subset the data to species collected from at least 10 populations (the 10-collection dataset). We used a cut-off of 10 for this analysis, rather than 20, because the focus is among-species variation and preliminary analyses suggested that 10 samples was adequate to estimate coefficient of variation and allowed us to include an additional 14 species and 129 collections.

We aggregated environmental and climate variables across all collections for each species, identifying the mean and range of each of the 14 environmental covariates (Table S1), except for the 4 measures of climate change for which we only calculated mean, for a total of 24 environmental covariates. We also calculated the mean year of collection for each entire species, as well as the mean day of year collections were made for each species. We then calculated the difference between the first and last year of collection as well as the first and last day of year to measure the range of each species' collection across years and collecting dates. This added 4 temporal covariates. Finally, for each species, we calculated the minimum and maximum latitude and longitudes of the collection sites. In total, we considered 31 environmental, spatial, and temporal variables in addition to growth form and duration, for a total of 33 variables, describing each of 119 species.

### Supplemental Results

#### *Spatial distribution of seed mass*

The inclusion of plant family as a random intercept effect significantly improved the model fit compared to the GLM ($\chi^2 = 6138.1$, $\Delta$ AIC = 6137, df = 1, $p < 0.001$, Table S3) and consistent with phylogenetic differences in seed mass among plant families (variance = 4.605) beyond spatial and elevational patterns.

We observed significant differences in seed-environment relationships in shrubs compared to other growth forms: the effect of latitude alone was positive in all models except shrubs alone, for which it was negative (Table S2). The effect of latitude was also positive in both annual models (GLM: $\beta = 0.076$, $p <0.001$, GLMM: $\beta = 0.062$, $p < 0.001$) and the perennial mixed model (GLMM: $\beta = 0.019$, $p < 0.001$). There were no significant spatial patterns of seed mass in biennials. Elevation, although statistically significant in the GLMM, had a near zero effect size in both models (GLM: $\beta = 0.00003$, $p = 0.2294$, GLMM: $\beta = 0.00009$, $p < 0.001$).

**Table S1.** Environmental and temporal variables used to describe collection site conditions, after removing correlated variables.

| Abbreviation | Definition |
|---|---|
| DeltFallMin | Change in fall minimum temperature over 30 years[1] |
| DeltFallMax | Change in fall maximum temperature over 30 years[1] |
| DeltSpringMax | Change in spring maximum temperature over 30 years[1] |
| DeltPrecip | Change in annual precipitation over 30 years[2] |
| AnnMinTemp | Minimum temperature from year of collection (ºC)[3] |
| SixMonthPrecip | Cumulative water from 6 months prior through collection date |
| Slope | Slope in degrees[4] |
| Aspect | Aspect in degrees from north[4] |
| packm | Snowpack of the previous month calculated as previous month precipitation multiplied by a melt factor determined by monthly mean temperature [5] |
| heatload | Heat Load Index calculated from slope, folded aspect, and latitude [6] |
| AWC | Available water content; expressed as a fraction (water contents at field capacity compared to permanent wilting point).[7] |
| clay | Mineral particles less than 0.002mm in diameter as a percentage extracted from |

[1] Calculated by obtaining either the absolute minimum or maximum daily temperature from September 1 to November 30 (fall) or March 1 to May 31 (spring) each year from 30 years prior to collection to year of collection and measuring the total change (regression slope) over 30 years (PRISM Climate Group)
[2] Calculated by obtaining annual total precipitation each year from 30 years prior to collection to year of collection and measuring the total change (regression slope) over 30 years (PRISM Climate Group)
[3] PRISM Climate Group
[4] Klingebiel et al. 2015
[5] Dilts et al., 2015; Lutz et al., 2010
[6] Dilts et al., 2015, McCune and Keon 2002
[7] Beaudette et al., 2021, Soil Survey Staff

| Abbreviation | Definition |
| --- | --- |
| | SSURGO [7] |
| SoilpH | Relative acidity or alkalinity of the soil extracted from SSURGO[7] |
| organic | Amount of decomposed plant and animal residue extracted from SSURGO[7] |
| Year | Year collection was made |
| Ydays | Day of year that collection was made where January 1 = 1 and December 31 = 365 |

85

**Table S2.** Results of generalized linear model and generalized linear mixed model testing the effects of latitude, longitude, and elevation on seed mass using the full dataset of 13004 populations of 2092 species from 108 plant families, and GLM and GLMMs with the inclusion of growth form or duration as interaction effects (due to the lack of adequate combinations we could not run growth form and duration together). The response variable, seed mass, is log transformed. Further models were run by subsetting the full dataset by the most collected growth forms: grasses (N = 2764 populations, 241 species, 4 plant families), forbs (N = 6857 populations, 1420 species, 69 plant families), and shrubs (N = 3212 populations, 363 species, 47 plant families); and duration: annuals (N = 1837 populations, 380 species, 36 plant families), biennials (N = 129 populations, 6 species, 6 plant families), and perennials (N = 10211 populations, 1510 species, 103 plant families).

| Model & parameters | *estimate* | *SE* | *t* | *p* | *lower 95% CI* | *upper 95% CI* |
|---|---|---|---|---|---|---|
| GLM Full dataset | | | | | | |
| *Fixed effects* | | | | | | |
| (intercept) | -1.578 | 0.357 | -4.418 | <0.001 | | |
| latitude | 0.027 | 0.004 | 6.253 | <0.001 | 0.018 | 0.035 |
| longitude | -0.005 | 0.003 | -1.703 | 0.089 | -0.012 | 0.001 |
| elevation | <0.001 | <0.001 | 1.202 | 0.229 | < 0.001 | <0.001 |
| GLMM Full dataset | | | | | | |
| *Fixed effects* | | | | | | |
| (intercept) | -2.099 | 0.361 | -5.811 | <0.001 | | |
| latitude | 0.031 | 0.003 | 9.230 | <0.001 | 0.025 | 0.038 |

| Model & parameters | *estimate* | *SE* | *t* | *p* | *lower 95% CI* | *upper 95% CI* |
|---|---|---|---|---|---|---|
| longitude | -0.010 | 0.002 | -4.128 | <0.001 | -0.015 | -0.005 |
| elevation | <0.001 | <0.001 | 4.553 | <0.001 | <0.001 | <0.001 |
| *Random intercept effects* | | | | | | |
| plant family | 4.605 | 2.146 | | | | |
| GLM Full dataset, duration | | | | | | |
| *Fixed effects* | $\chi^2$ | *p* | | | | |
| latitude | 7.87 | 0.005 | | | | |
| longitude | 4.07 | 0.044 | | | | |
| elevation | 6.89 | 0.009 | | | | |
| duration | 637 | <0.001 | | | | |
| latitude * duration | 36.34 | <0.001 | | | | |
| longitude * duration | 29.28 | <0.001 | | | | |
| elevation * duration | 27.19 | <0.001 | | | | |
| GLM Full dataset, duration | | | | | | |
| *Fixed effects* | | | | | | |
| (intercept) (annual) | 1.072 | 1.056 | 1.016 | 0.31 | -0.997 | 3.141 |
| latitude | 0.076 | 0.014 | 5.603 | <0.001 | 0.049 | 0.102 |
| longitude | 0.04 | 0.01 | 3.797 | <0.001 | 0.019 | 0.06 |

| Model & parameters | *estimate* | *SE* | *t* | *p* | *lower 95% CI* | *upper 95% CI* |
|---|---|---|---|---|---|---|
| elevation | <0.001 | <0.001 | 0.801 | 0.423 | <0.001 | <0.001 |
| biennial | -8.423 | 6.27 | -1.344 | 0.179 | -20.711 | 3.865 |
| perennial | -2.103 | 1.125 | -1.868 | 0.062 | -4.308 | 0.103 |
| variable | -2.091 | 1.658 | -1.261 | 0.207 | -5.342 | 1.159 |
| latitude * biennial | -0.085 | 0.079 | -1.081 | 0.28 | -0.239 | 0.069 |
| latitude * perennial | -0.075 | 0.014 | -5.267 | <0.001 | -0.103 | -0.047 |
| latitude * variable | -0.02 | 0.021 | -0.983 | 0.326 | -0.061 | 0.02 |
| longitude * biennial | -0.098 | 0.054 | -1.81 | 0.07 | -0.205 | 0.008 |
| longitude * perennial | -0.053 | 0.011 | -4.807 | <0.001 | -0.075 | -0.031 |
| longitude * variable | -0.021 | 0.016 | -1.338 | 0.181 | -0.051 | 0.01 |
| elevation * biennial | <0.001 | 0.001 | -0.282 | 0.778 | -0.001 | 0.001 |
| elevation * perennial | <0.001 | <0.001 | -2.153 | 0.031 | <0.001 | <0.001 |
| elevation * variable | <0.001 | <0.001 | 2.525 | 0.012 | <0.001 | 0.001 |
| GLM Full dataset, duration | | | | | | |
| *Fixed effects* | | | | | | |
| (intercept) (annual) | -1.182 | 9327.694 | -1.335 | 0.182 | -2.917 | 0.552 |
| latitude | 0.045 | 12103.092 | 4.113 | <0.001 | 0.023 | 0.066 |
| longitude | 0.006 | 12100.906 | 0.719 | 0.472 | -0.011 | 0.023 |
| elevation | <0.001 | 12101.442 | 2.021 | 0.043 | <0.001 | <0.001 |
| biennial | -4.788 | 12089.991 | -0.976 | 0.329 | -14.4 | 4.823 |
| perennial | -0.826 | 12109.89 | -0.901 | 0.367 | -2.62 | 0.969 |

| Model & parameters | *estimate* | *SE* | *t* | *p* | *lower 95% CI* | *upper 95% CI* |
|---|---|---|---|---|---|---|
| variable | 0.546 | 12102.801 | 0.412 | 0.681 | -2.054 | 3.147 |
| latitude * biennial | -0.057 | 12089.856 | -0.933 | 0.351 | -0.178 | 0.063 |
| latitude * perennial | -0.026 | 12105.029 | -2.259 | 0.024 | -0.048 | -0.003 |
| latitude * variable | 0.002 | 12098.931 | 0.142 | 0.887 | -0.03 | 0.034 |
| longitude * biennial | -0.061 | 12089.856 | -1.441 | 0.15 | -0.145 | 0.022 |
| longitude * perennial | -0.022 | 12102.45 | -2.425 | 0.015 | -0.039 | -0.004 |
| longitude * variable | 0.006 | 12099.658 | 0.459 | 0.646 | -0.019 | 0.03 |
| elevation * biennial | <0.001 | 12089.728 | -0.68 | 0.497 | -0.001 | 0.001 |
| elevation * perennial | <0.001 | 12104.742 | -1.498 | 0.134 | <0.001 | <0.001 |
| elevation * variable | <0.001 | 12103.586 | 1.18 | 0.238 | <0.001 | <0.001 |
| *Random intercept effects* | | | | | | |
| plant family | 4.413 | 2.101 | | | | |
| GLM Full dataset, growth form | | | | | | |
| *Fixed effects* | $\chi^2$ | *p* | | | | |
| latitude | 90.01 | <0.001 | | | | |
| longitude | 0.34 | 0.558 | | | | |
| elevation | 13.81 | <0.001 | | | | |
| growth form | 786 | <0.001 | | | | |
| latitude * growth form | 295 | <0.001 | | | | |

| Model & parameters | *estimate* | *SE* | *t* | *p* | *lower 95% CI* | *upper 95% CI* |
|---|---|---|---|---|---|---|
| longitude * growth form | 8.37 | 0.079 | | | | |
| elevation * growth form | 5.98 | 0.201 | | | | |
| GLM Full dataset, growth form | | | | | | |
| *Fixed effects* | | | | | | |
| intercept (forbs) | -2.064 | 0.479 | -4.312 | <0.001 | -3.002 | -1.126 |
| latitude | 0.073 | 0.006 | 12.875 | <0.001 | 0.062 | 0.084 |
| longitude | 0.009 | 0.004 | 2.194 | 0.028 | 0.001 | 0.018 |
| elevation | <0.001 | <0.001 | 2.498 | 0.012 | <0.001 | <0.001 |
| graminoid | -2.653 | 0.825 | -3.215 | 0.001 | -4.271 | -1.036 |
| shrub | 5.493 | 0.895 | 6.137 | <0.001 | 3.739 | 7.247 |
| tree | 13.857 | 6.235 | 2.222 | 0.026 | 1.637 | 26.077 |
| vine | 19.67 | 11.204 | 1.756 | 0.079 | -2.288 | 41.629 |
| latitude * graminoid | 0.018 | 0.010 | 1.774 | 0.076 | -0.002 | 0.039 |
| latitude * shrub | -0.162 | 0.010 | -15.493 | <0.001 | -0.182 | -0.141 |
| latitude * tree | -0.215 | 0.069 | -3.103 | 0.002 | -0.351 | -0.079 |
| latitude * vine | -0.229 | 0.103 | -2.234 | 0.025 | -0.43 | -0.028 |
| longitude * graminoid | -0.018 | 0.008 | -2.396 | 0.017 | -0.033 | -0.003 |
| longitude * shrub | -0.015 | 0.008 | -1.894 | 0.058 | -0.03 | 0.001 |

| Model & parameters | *estimate* | *SE* | *t* | *p* | *lower 95% CI* | *upper 95% CI* |
|---|---|---|---|---|---|---|
| longitude * tree | 0.024 | 0.056 | 0.44 | 0.66 | -0.084 | 0.133 |
| longitude * vine | 0.066 | 0.082 | 0.795 | 0.426 | -0.096 | 0.227 |
| elevation * graminoid | <0.001 | <0.001 | 1.271 | 0.204 | <0.001 | <0.001 |
| elevation * shrub | <0.001 | <0.001 | -0.313 | 0.755 | <0.001 | <0.001 |
| elevation * tree | <0.001 | <0.001 | 0.327 | 0.744 | -0.001 | 0.001 |
| elevation * vine | -0.001 | 0.001 | -1.934 | 0.053 | -0.003 | <0.001 |
| GLMM Full dataset, growth form | | | | | | |
| *Fixed effects* | | | | | | |
| (intercept) (forbs) | -0.798 | 0.442 | -1.806 | 0.071 | -1.662 | 0.066 |
| latitude | 0.043 | 0.005 | 9.251 | <0.001 | 0.034 | 0.052 |
| longitude | 0.007 | 0.003 | 1.942 | 0.052 | <0.001 | 0.013 |
| elevation | <0.001 | <0.001 | 4.707 | <0.001 | <0.001 | <0.001 |
| graminoid | -8.312 | 1.22 | -6.814 | <0.001 | -10.694 | -5.93 |
| shrub | 3.36 | 0.756 | 4.443 | <0.001 | 1.88 | 4.843 |
| tree | -0.761 | 5.796 | -0.131 | 0.896 | -12.088 | 10.614 |
| vine | 12.17 | 9.874 | 1.233 | 0.218 | -7.111 | 31.591 |
| latitude * graminoid | 0.055 | 0.008 | 6.631 | <0.001 | 0.039 | 0.071 |
| latitude * shrub | -0.105 | 0.009 | -12.212 | <0.001 | -0.121 | -0.088 |
| latitude * tree | -0.077 | 0.059 | -1.288 | 0.198 | -0.193 | 0.04 |
| latitude * vine | -0.169 | 0.086 | -1.975 | 0.048 | -0.338 | -0.002 |

| Model & parameters | *estimate* | *SE* | *t* | *p* | *lower 95% CI* | *upper 95% CI* |
|---|---|---|---|---|---|---|
| longitude * graminoid | -0.041 | 0.006 | -6.648 | <0.001 | -0.052 | -0.029 |
| longitude * shrub | -0.01 | 0.006 | -1.582 | 0.114 | -0.022 | 0.002 |
| longitude * tree | -0.042 | 0.048 | -0.882 | 0.378 | -0.136 | 0.052 |
| longitude * vine | 0.031 | 0.069 | 0.446 | 0.655 | -0.105 | 0.168 |
| elevation * graminoid | <0.001 | <0.001 | 0.125 | 0.901 | <0.001 | <0.001 |
| elevation * shrub | <0.001 | <0.001 | -0.816 | 0.414 | <0.001 | <0.001 |
| elevation * tree | <0.001 | <0.001 | -0.131 | 0.896 | -0.001 | 0.001 |
| elevation * vine | -0.001 | 0.001 | -1.714 | 0.087 | -0.002 | <0.001 |
| *Random intercept effects* | | | | | | |
| plant family | 3.881 | 1.970 | | | | |
| GLM Grasses | | | | | | |
| *Fixed effects* | | | | | | |
| (intercept) | -4.717 | 0.590 | -7.993 | <0 .001 | -5.874 | -3.560 |
| latitude | 0.091 | 0.008 | 11.921 | <0.001 | 0.076 | 0.106 |
| longitude | -0.009 | 0.006 | -1.616 | 0.106 | -0.020 | 0.002 |
| elevation | <0.001 | <0.001 | 3.314 | <0.001 | <0.001 | <0.001 |
| GLMM Grasses | | | | | | |
| *Fixed effects* | | | | | | |
| (intercept) | -9.11 | 1.059 | -8.603 | <0.001 | -11.306 | 6.912 |
| latitude | 0.098 | 0.006 | 15.328 | <0.001 | 0.085 | 0.110 |

| Model & parameters | *estimate* | *SE* | *t* | *p* | *lower 95% CI* | *upper 95% CI* |
|---|---|---|---|---|---|---|
| longitude | -0.034 | 0.005 | -7.173 | <0.001 | -0.043 | -0.025 |
| elevation | <0.001 | <0.001 | 3.157 | 0.002 | <0.001 | <0.001 |
| *Random intercept effects* | | | | | | |
| plant family | 3.358 | 1.832 | | | | |
| GLM Forbs | | | | | | |
| *Fixed effects* | | | | | | |
| (intercept) | -2.064 | 0.476 | -4.335 | <0.001 | -2.997 | 1.131 |
| latitude | 0.073 | 0.006 | 12.945 | <0.001 | 0.062 | 0.084 |
| longitude | 0.009 | 0.004 | 2.206 | 0.0274 | 0.001 | 0.018 |
| elevation | <0.001 | <0.001 | 2.512 | 0.0120 | <0.001 | <0.001 |
| GLMM Forbs | | | | | | |
| *Fixed effects* | | | | | | |
| (intercept) | -2.076 | 0.436 | -4.761 | <0.001 | -2.930 | -1.222 |
| latitude | 0.047 | 0.004 | 10.618 | <0.001 | 0.039 | 0.056 |
| longitude | 0.002 | 0.003 | 0.576 | 0.564 | -0.005 | 0.008 |
| elevation | <0.001 | <0.001 | 6.816 | <0.001 | <0.001 | <0.001 |
| *Random intercept effects* | | | | | | |
| plant family | 2.971 | 1.724 | | | | |

| Model & parameters | *estimate* | *SE* | *t* | *p* | *lower 95% CI* | *upper 95% CI* |
|---|---|---|---|---|---|---|
| GLM Shrubs | | | | | | |
| *Fixed effects* | | | | | | |
| (intercept) | 3.429 | 0.832 | 4.119 | <0.001 | 1.798 | 5.061 |
| latitude | -0.089 | 0.010 | -9.212 | <0.001 | -0.108 | -0.070 |
| longitude | -0.005 | 0.007 | -0.752 | 0.452 | -0.019 | 0.009 |
| elevation | <0.001 | <0.001 | 1.110 | 0.267 | <0.001 | <0.001 |
| GLMM Shrubs | | | | | | |
| *Fixed effects* | | | | | | |
| (intercept) | 3.021 | 0.748 | 4.039 | <0.001 | 1.557 | 4.486 |
| latitude | -0.049 | 0.007 | -6.585 | <0.001 | -0.064 | -0.035 |
| longitude | <0.001 | 0.006 | 0.023 | 0.982 | -0.011 | 0.011 |
| elevation | <0.001 | <0.001 | 1.265 | 0.206 | <0.001 | <0.001 |
| *Random intercept effects* | | | | | | |
| plant family | 4.278 | 2.068 | | | | |
| GLM Annuals | | | | | | |
| *Fixed effects* | | | | | | |
| (intercept) | 1.072 | 0.968 | 1.108 | 0.268 | -0.824 | 2.969 |
| latitude | 0.076 | 0.012 | 6.113 | <0.001 | 0.051 | 0.100 |
| longitude | 0.040 | 0.010 | 4.142 | <0.001 | 0.021 | 0.059 |
| elevation | <0.001 | <0.001 | 0.873 | 0.383 | <0.001 | <0.001 |

| Model & parameters | *estimate* | *SE* | *t* | *p* | *lower 95% CI* | *upper 95% CI* |
|---|---|---|---|---|---|---|
| GLMM Annuals | | | | | | |
| *Fixed effects* | | | | | | |
| (intercept) | -1.286 | 1212.8 | -1.608 | 0.108 | -2.851 | 0.279 |
| latitude | 0.062 | 1593.769 | 6.612 | <0.001 | 0.043 | 0.080 |
| longitude | 0.018 | 1592.582 | 2.429 | 0.015 | 0.003 | 0.032 |
| elevation | <0.001 | 1586.481 | 3.662 | <0.001 | <0.001 | <0.001 |
| *Random intercept effects* | | | | | | |
| plant family | 1.910 | 1.382 | | | | |
| GLM Biennials | | | | | | |
| *Fixed effects* | | | | | | |
| (intercept) | -7.351 | 7.111 | -1.034 | 0.309 | -21.289 | 6.586 |
| latitude | -0.009 | 0.089 | -0.103 | 0.918 | -0.184 | 0.165 |
| longitude | -0.059 | 0.061 | -0.954 | 0.347 | -0.179 | 0.062 |
| elevation | <0.001 | 0.001 | -0.138 | 0.891 | -0.001 | 0.001 |
| GLMM Biennials | | | | | | |
| *Fixed effects* | | | | | | |
| (intercept) | -7.351 | 32 | -1.034 | 0.309 | -20.850 | 6.148 |
| latitude | -0.009 | 32 | -0.103 | 0.918 | -0.178 | 0.160 |
| longitude | -0.059 | 32 | -0.954 | 0.347 | -0.175 | 0.058 |

| Model & parameters | *estimate* | *SE* | *t* | *p* | *lower 95% CI* | *upper 95% CI* |
|---|---|---|---|---|---|---|
| elevation | <0.001 | 32 | -0.138 | 0.891 | -0.001 | 0.001 |
| *Random intercept effects* | | | | | | |
| plant family | <0.001 | <0.001 | | | | |
| GLM Perennials | | | | | | |
| *Fixed effects* | | | | | | |
| (intercept) | -1.030 | 0.401 | -2.571 | 0.010 | -1.816 | -0.245 |
| latitude | <0.001 | 0.005 | 0.096 | 0.924 | -0.009 | 0.010 |
| longitude | -0.013 | 0.004 | -3.761 | <0.001 | -0.020 | -0.006 |
| elevation | <0.001 | <0.001 | -4.139 | <0.001 | <0.001 | <0.001 |
| GLMM Perennials | | | | | | |
| *Fixed effects* | | | | | | |
| (intercept) | -1.877 | 933.387 | -4.727 | <0.001 | -2.655 | -1.099 |
| latitude | 0.019 | 9573.906 | 4.841 | <0.001 | 0.011 | 0.026 |
| longitude | -0.016 | 9564.48 | -5.503 | <0.001 | -0.021 | -0.010 |
| elevation | <0.001 | 9572.876 | 0.633 | 0.527 | <0.001 | <0.001 |
| *Random intercept effects* | | | | | | |
| plant family | 4.550 | 2.133 | | | | |

97

**Table S3.** Comparison of generalized linear model and generalized linear mixed models (Table S2) testing the effects of latitude, longitude, and elevation on seed mass. Both GLM and GLMMs have the same predictor variables and response variable (log transformed seed mass), the GLMM has the addition of plant family as a random intercept term. Models are compared in pairs, first using the full dataset (N = 13004 populations, 2092 species,108 plant families) GLM to GLMM, then comparing grass (N = 2764 populations, 241 species, 4 plant families) GLM to GLMM, forb (N = 6857 populations, 1420 species, 69 plant families) GLM to GLMM, and finally shrub (N = 3212 populations, 363 species, 47 plant families) GLM to GLMM.

| Model | Par | AIC | BIC | logLik | Deviance | $\chi^2$ | *df* | *p* |
|---|---|---|---|---|---|---|---|---|
| GLM Full | 5 | 46954 | 46991 | -23472 | 46944 | | | |
| GLMM Full | 6 | 40817 | 40862 | -20403 | 40805 | 6138.1 | 1 | <0.001 |
| GLM Grass | 5 | 9131.7 | 9161.1 | -4560.8 | 9121.7 | | | |
| GLMM Grass | 6 | 8198.8 | 8234.1 | -4093.4 | 8186.8 | 934.89 | 1 | <0.001 |
| GLM Forbs | 5 | 24102 | 24136 | -12046 | 24092 | | | |
| GLMM Forbs | 6 | 20706 | 20747 | -10347 | 20694 | 3397.5 | 1 | <0.001 |
| GLM Shrubs | 5 | 11835 | 11865 | -5912.4 | 11825 | | | |
| GLMM Shrubs | 6 | 10093 | 10129 | -5040.4 | 10081 | 1744 | 1 | <0.001 |

**Table S4.** Results of spatial simultaneous autoregressive error model (SAR) testing the effects of climate and environment on seed mass from the 3 collection dataset (N = 10,958 populations, 859 species, 73 plant families). Covariates were selected using random forest and are centered and scaled.

| *Covariate* | *Estimate* | *SE* | *z* | *p* | *lower 95% CI* | *Upper 95% CI* |
|---|---|---|---|---|---|---|
| Pack | -0.004 | 0.010 | -0.429 | 0.668 | -0.023 | 0.015 |
| Δ Fall max temp | 0.002 | 0.010 | 0.219 | 0.826 | -0.018 | 0.022 |
| Δ Spring max temp | 0.006 | 0.012 | 0.527 | 0.598 | -0.018 | 0.022 |
| Slope | 0.045 | 0.011 | 3.955 | <0.001 | 0.023 | 0.067 |
| AWC | 0.005 | 0.013 | 0.377 | 0.706 | -0.021 | 0.031 |
| Δ Precip | -0.011 | 0.011 | -0.964 | 0.335 | -0.033 | 0.011 |
| Soil pH | -0.001 | 0.013 | -0.042 | 0.966 | -0.026 | 0.025 |
| Heatload | 0.024 | 0.012 | 2.072 | 0.038 | 0.001 | 0.046 |
| Day of year | <0.001 | <0.001 | -0.098 | 0.922 | <0.001 | <0.001 |
| Year | -0.010 | 0.004 | -2.716 | 0.007 | -0.017 | -0.003 |
| Organic matter | -0.019 | 0.013 | -1.481 | 0.139 | -0.045 | 0.006 |
| Clay content | 0.045 | 0.011 | 3.894 | <0.001 | 0.022 | 0.067 |
| Six month precip | 0.051 | 0.012 | 4.339 | <0.001 | 0.028 | 0.074 |
| Annual min temp | 0.044 | 0.011 | 3.869 | <0.001 | 0.022 | 0.066 |

**Table S5.** Results of 104 individual spatial simultaneous autoregressive error model examining the effects of environment, climate, and time on seed mass from the 20 collection dataset (5,003 total populations). Covariates were centered and scaled and selected using random forest. Seed mass was log transformed.

| Species | N | Formula | $R^2$ | Covariate | Est. | SE | *z* | *p* | *lower 95% CI* | *upper 95% CI* |
|---|---|---|---|---|---|---|---|---|---|---|
| *Achillea millefolium* | 78 | Seed mass ~ SoilpH | 0.158 | (Intercept) | -2.098 | 0.022 | -96.336 | < **0.001** | -2.141 | -2.056 |
| | | | | SoilpH | -0.072 | 0.020 | -3.509 | < **0.001** | -0.112 | -0.032 |
| *Achnatherum hymenoides* | 218 | Seed mass ~ Ydays + DeltFallMax + DeltSpringMax + AWC + SoilpH + clay + organic + AnnMinTemp | 0.307 | (Intercept) | 1.334 | 0.029 | 46.129 | < **0.001** | 1.277 | 1.391 |

| Species | N | Formula | $R^2$ | Covariate | Est. | SE | *z* | *p* | *lower 95% CI* | *upper 95% CI* |
|---|---|---|---|---|---|---|---|---|---|---|
| | | | | AnnMinTemp | 0.149 | 0.032 | 4.696 | < **0.001** | 0.087 | 0.211 |
| | | | | AWC | 0.055 | 0.030 | 1.856 | 0.063 | -0.003 | 0.113 |
| | | | | clay | -0.030 | 0.030 | -1.004 | 0.315 | -0.088 | 0.028 |
| | | | | DeltFallMax | -0.010 | 0.023 | -0.429 | 0.668 | -0.056 | 0.036 |
| | | | | DeltSpringMax | -0.007 | 0.023 | -0.290 | 0.772 | -0.052 | 0.038 |
| | | | | organic | 0.003 | 0.027 | 0.116 | 0.908 | -0.050 | 0.056 |
| | | | | SoilpH | 0.060 | 0.028 | 2.183 | **0.029** | 0.006 | 0.115 |
| | | | | Ydays | 0.000 | 0.001 | 0.450 | 0.653 | -0.002 | 0.002 |
| *Achnatherum speciosum* | 30 | Seed mass ~ Ydays + DeltFallMax + DeltSpringMax + AWC + SoilpH + clay + | 0.349 | (Intercept) | 1.058 | 0.022 | 47.127 | < **0.001** | 1.014 | 1.102 |

| Species | N | Formula | $R^2$ | Covariate | Est. | SE | *z* | *p* | *lower 95% CI* | *upper 95% CI* |
|---|---|---|---|---|---|---|---|---|---|---|
| | | organic + AnnMinTemp | | | | | | | | |
| | | | | AnnMinTemp | -0.017 | 0.034 | -0.502 | 0.616 | -0.084 | 0.050 |
| | | | | AWC | 0.107 | 0.046 | 2.337 | **0.019** | 0.017 | 0.197 |
| | | | | clay | -0.083 | 0.040 | -2.053 | **0.040** | -0.161 | -0.004 |
| | | | | DeltFallMax | -0.076 | 0.043 | -1.762 | 0.078 | -0.161 | 0.009 |
| | | | | DeltSpringMax | 0.050 | 0.040 | 1.252 | 0.211 | -0.028 | 0.128 |
| | | | | organic | -0.058 | 0.037 | -1.572 | 0.116 | -0.131 | 0.014 |
| | | | | SoilpH | -0.040 | 0.036 | -1.100 | 0.271 | -0.111 | 0.031 |
| | | | | Ydays | -0.001 | 0.002 | -0.727 | 0.467 | -0.005 | 0.002 |
| *Achnatherum thurberianum* | 39 | Seed mass ~ Ydays + SixMonthPrecip | 0.632 | (Intercept) | 1.267 | 0.030 | 42.180 | < **0.001** | 1.208 | 1.326 |

| Species | N | Formula | $R^2$ | Covariate | Est. | SE | *z* | *p* | *lower 95% CI* | *upper 95% CI* |
|---|---|---|---|---|---|---|---|---|---|---|
| | | + DeltFallMax + SoilpH + clay + organic + AnnMinTemp | | | | | | | | |
| | | | | AnnMinTemp | -0.039 | 0.038 | -1.033 | 0.302 | -0.114 | 0.035 |
| | | | | clay | 0.092 | 0.033 | 2.761 | **0.006** | 0.027 | 0.157 |
| | | | | DeltFallMax | -0.048 | 0.032 | -1.489 | 0.137 | -0.111 | 0.015 |
| | | | | organic | -0.025 | 0.036 | -0.696 | 0.486 | -0.095 | 0.045 |
| | | | | SixMonthPrecip | -0.110 | 0.034 | -3.226 | **0.001** | -0.177 | -0.043 |
| | | | | SoilpH | -0.016 | 0.033 | -0.495 | 0.621 | -0.082 | 0.049 |
| | | | | Ydays | -0.007 | 0.002 | -2.746 | **0.006** | -0.012 | -0.002 |
| *Agastache urticifolia* | 38 | Seed mass ~ DeltPrecip | 0.058 | (Intercept) | -0.491 | 0.023 | -21.650 | < **0.001** | -0.535 | -0.446 |
| | | | | DeltPrecip | -0.036 | 0.023 | -1.551 | 0.121 | -0.081 | 0.009 |

| Species | N | Formula | $R^2$ | Covariate | Est. | SE | *z* | *p* | *lower 95% CI* | *upper 95% CI* |
|---|---|---|---|---|---|---|---|---|---|---|
| *Agoseris glauca* | 20 | Seed mass ~ packm + Ydays | 0.320 | (Intercept) | 0.818 | 0.047 | 17.551 | < **0.001** | 0.726 | 0.909 |
| | | | | packm | 0.111 | 0.053 | 2.105 | **0.035** | 0.008 | 0.214 |
| | | | | Ydays | -0.004 | 0.003 | -1.482 | 0.138 | -0.009 | 0.001 |
| *Allium acuminatum* | 24 | Seed mass ~ AWC | 0.413 | (Intercept) | 0.701 | 0.049 | 14.299 | < **0.001** | 0.605 | 0.797 |
| | | | | AWC | 0.098 | 0.032 | 3.067 | **0.002** | 0.036 | 0.161 |
| *Allium textile* | 28 | Seed mass ~ Slope + organic | 0.491 | (Intercept) | 1.097 | 0.024 | 45.122 | < **0.001** | 1.049 | 1.144 |
| | | | | organic | -0.102 | 0.024 | -4.258 | < **0.001** | -0.149 | -0.055 |
| | | | | Slope | -0.031 | 0.024 | -1.281 | 0.200 | -0.079 | 0.016 |
| *Amaranthus fimbriatus* | 23 | Seed mass ~ Slope + organic | 0.112 | (Intercept) | -1.619 | 0.014 | -111.818 | < **0.001** | -1.647 | -1.591 |
| | | | | organic | 0.038 | 0.018 | 2.119 | **0.034** | 0.003 | 0.072 |

| Species | N | Formula | $R^2$ | Covariate | Est. | SE | *z* | *p* | *lower 95% CI* | *upper 95% CI* |
|---|---|---|---|---|---|---|---|---|---|---|
| | | | | Slope | -0.004 | 0.019 | -0.217 | 0.829 | -0.041 | 0.033 |
| *Ambrosia dumosa* | 75 | Seed mass ~ Year + Ydays + SixMonthPrecip + DeltFallMax + DeltPrecip + AnnMinTemp | 0.460 | (Intercept) | 1.633 | 0.024 | 68.896 | < **0.001** | 1.586 | 1.679 |
| | | | | AnnMinTemp | 0.107 | 0.026 | 4.153 | < **0.001** | 0.057 | 0.158 |
| | | | | DeltFallMax | -0.042 | 0.028 | -1.478 | 0.140 | -0.097 | 0.014 |
| | | | | DeltPrecip | 0.045 | 0.033 | 1.366 | 0.172 | -0.020 | 0.109 |
| | | | | SixMonthPrecip | 0.088 | 0.029 | 3.035 | **0.002** | 0.031 | 0.145 |
| | | | | Ydays | 0.001 | 0.001 | 2.055 | **0.040** | 0.000 | 0.002 |
| | | | | Year | -0.025 | 0.009 | -2.821 | **0.005** | -0.043 | -0.008 |

| Species | N | Formula | $R^2$ | Covariate | Est. | SE | *z* | *p* | *lower 95% CI* | *upper 95% CI* |
|---|---|---|---|---|---|---|---|---|---|---|
| *Amsinckia tessellata* | 20 | Seed mass ~ packm | 0.509 | (Intercept) | 1.194 | 0.028 | 43.322 | < **0.001** | 1.140 | 1.248 |
| | | | | packm | -0.104 | 0.024 | -4.298 | < **0.001** | -0.152 | -0.057 |
| *Argemone munita* | 26 | Seed mass ~ SoilpH | 0.334 | (Intercept) | 1.187 | 0.019 | 61.365 | < **0.001** | 1.149 | 1.225 |
| | | | | SoilpH | -0.060 | 0.019 | -3.203 | **0.001** | -0.097 | -0.023 |
| *Aristida purpurea* | 45 | Seed mass ~ Year + Ydays | 0.055 | (Intercept) | 0.717 | 0.059 | 12.121 | < **0.001** | 0.601 | 0.833 |
| | | | | Ydays | -0.001 | 0.001 | -1.157 | 0.247 | -0.004 | 0.001 |
| | | | | Year | 0.028 | 0.023 | 1.221 | 0.222 | -0.017 | 0.072 |
| *Artemisia cana* | 21 | Seed mass ~ heatload + DeltSpringMax | 0.566 | (Intercept) | -1.206 | 0.050 | -24.016 | < **0.001** | -1.304 | -1.107 |

| Species | N | Formula | $R^2$ | Covariate | Est. | SE | *z* | *p* | *lower 95% CI* | *upper 95% CI* |
|---|---|---|---|---|---|---|---|---|---|---|
| | | + DeltPrecip + clay | | | | | | | | |
| | | | | clay | 0.172 | 0.068 | 2.510 | **0.012** | 0.038 | 0.306 |
| | | | | DeltPrecip | -0.162 | 0.075 | -2.150 | **0.032** | -0.310 | -0.014 |
| | | | | DeltSpringMax | -0.038 | 0.060 | -0.627 | 0.530 | -0.155 | 0.080 |
| | | | | heatload | 0.060 | 0.084 | 0.706 | 0.480 | -0.106 | 0.225 |
| *Artemisia frigida* | 21 | Seed mass ~ Aspect + heatload | 0.604 | (Intercept) | -2.294 | 0.054 | -42.410 | < **0.001** | -2.400 | -2.188 |
| | | | | Aspect | 0.325 | 0.070 | 4.656 | < **0.001** | 0.188 | 0.461 |
| | | | | heatload | -0.261 | 0.059 | -4.436 | < **0.001** | -0.377 | -0.146 |
| *Artemisia nova* | 20 | Seed mass ~ Slope + | 0.441 | (Intercept) | -0.755 | 0.040 | -18.693 | < **0.001** | -0.834 | -0.676 |

| Species | N | Formula | $R^2$ | Covariate | Est. | SE | *z* | *p* | *lower 95% CI* | *upper 95% CI* |
|---|---|---|---|---|---|---|---|---|---|---|
| | | DeltFallMin + DeltPrecip | | | | | | | | |
| | | | | DeltFallMin | 0.015 | 0.042 | 0.359 | 0.719 | -0.068 | 0.098 |
| | | | | DeltPrecip | 0.084 | 0.042 | 1.987 | **0.047** | 0.001 | 0.167 |
| | | | | Slope | -0.094 | 0.041 | -2.273 | **0.023** | -0.174 | -0.013 |
| *Artemisia tridentata* | 198 | Seed mass ~ Year + SixMonthPrecip + DeltFallMax + DeltSpringMax + DeltPrecip + AWC + clay + AnnMinTemp | 0.321 | (Intercept) | -1.462 | 0.019 | -75.577 | < **0.001** | -1.500 | -1.424 |

| Species | N | Formula | $R^2$ | Covariate | Est. | SE | *z* | *p* | *lower 95% CI* | *upper 95% CI* |
|---|---|---|---|---|---|---|---|---|---|---|
| | | | | AnnMinTemp | 0.035 | 0.020 | 1.750 | 0.080 | -0.004 | 0.075 |
| | | | | AWC | -0.089 | 0.020 | -4.359 | < **0.001** | -0.129 | -0.049 |
| | | | | clay | 0.058 | 0.022 | 2.578 | **0.010** | 0.014 | 0.102 |
| | | | | DeltFallMax | 0.018 | 0.020 | 0.881 | 0.379 | -0.022 | 0.058 |
| | | | | DeltPrecip | -0.089 | 0.027 | -3.329 | **0.001** | -0.141 | -0.036 |
| | | | | DeltSpringMax | -0.011 | 0.023 | -0.490 | 0.624 | -0.055 | 0.033 |
| | | | | SixMonthPrecip | 0.043 | 0.021 | 2.061 | **0.039** | 0.002 | 0.084 |
| | | | | Year | -0.041 | 0.008 | -4.890 | < **0.001** | -0.057 | -0.024 |
| *Asclepias speciosa* | 22 | Seed mass ~ SoilpH + AnnMinTemp | 0.587 | (Intercept) | 1.871 | 0.028 | 66.953 | < **0.001** | 1.816 | 1.926 |
| | | | | AnnMinTemp | 0.126 | 0.029 | 4.302 | < **0.001** | 0.068 | 0.183 |
| | | | | SoilpH | -0.055 | 0.029 | -1.891 | 0.059 | -0.113 | 0.002 |

| Species | N | Formula | $R^2$ | Covariate | Est. | SE | *z* | *p* | *lower 95% CI* | *upper 95% CI* |
|---|---|---|---|---|---|---|---|---|---|---|
| *Atriplex canescens* | 126 | Seed mass ~ heatload + organic | 0.044 | (Intercept) | 2.184 | 0.038 | 57.377 | < **0.001** | 2.110 | 2.259 |
| | | | | heatload | -0.077 | 0.037 | -2.067 | **0.039** | -0.151 | -0.004 |
| | | | | organic | 0.024 | 0.037 | 0.654 | 0.513 | -0.048 | 0.096 |
| *Atriplex confertifolia* | 71 | Seed mass ~ Aspect + Ydays + SixMonthPrecip + SoilpH | 0.334 | (Intercept) | 1.620 | 0.055 | 29.441 | < **0.001** | 1.512 | 1.728 |
| | | | | Aspect | -0.059 | 0.044 | -1.347 | 0.178 | -0.146 | 0.027 |
| | | | | SixMonthPrecip | 0.097 | 0.049 | 1.963 | **0.050** | 0.000 | 0.194 |
| | | | | SoilpH | -0.129 | 0.049 | -2.666 | **0.008** | -0.224 | -0.034 |
| | | | | Ydays | -0.001 | 0.001 | -1.513 | 0.130 | -0.003 | 0.000 |

| Species | N | Formula | $R^2$ | Covariate | Est. | SE | *z* | *p* | *lower 95% CI* | *upper 95% CI* |
|---|---|---|---|---|---|---|---|---|---|---|
| *Atriplex gardneri* | 30 | Seed mass ~ AnnMinTemp | 0.545 | (Intercept) | 1.315 | 0.047 | 27.990 | < **0.001** | 1.223 | 1.407 |
| | | | | AnnMinTemp | 0.346 | 0.052 | 6.681 | < **0.001** | 0.245 | 0.448 |
| *Atriplex polycarpa* | 43 | Seed mass ~ SixMonthPrecip + DeltFallMin + DeltSpringMax + DeltPrecip | 0.396 | (Intercept) | -0.932 | 0.031 | -30.390 | < **0.001** | -0.992 | -0.872 |
| | | | | DeltFallMin | -0.041 | 0.025 | -1.632 | 0.103 | -0.091 | 0.008 |
| | | | | DeltPrecip | -0.056 | 0.026 | -2.169 | **0.030** | -0.107 | -0.005 |
| | | | | DeltSpringMax | -0.042 | 0.026 | -1.645 | 0.100 | -0.092 | 0.008 |
| | | | | SixMonthPrecip | -0.009 | 0.025 | -0.336 | 0.737 | -0.058 | 0.041 |

| Species | N | Formula | $R^2$ | Covariate | Est. | SE | *z* | *p* | *lower 95% CI* | *upper 95% CI* |
|---|---|---|---|---|---|---|---|---|---|---|
| *Atriplex torreyi* | 25 | Seed mass ~ packm + clay | 0.350 | (Intercept) | -0.729 | 0.054 | -13.619 | < **0.001** | -0.834 | -0.624 |
| | | | | clay | 0.032 | 0.053 | 0.604 | 0.546 | -0.072 | 0.136 |
| | | | | packm | -0.184 | 0.051 | -3.639 | < **0.001** | -0.283 | -0.085 |
| *Baileya multiradiata* | 24 | Seed mass ~ Slope + Aspect + packm + Ydays + clay + AnnMinTemp | 0.714 | (Intercept) | -0.567 | 0.035 | -15.995 | < **0.001** | -0.637 | -0.498 |
| | | | | AnnMinTemp | 0.076 | 0.056 | 1.369 | 0.171 | -0.033 | 0.185 |
| | | | | Aspect | 0.081 | 0.041 | 1.988 | **0.047** | 0.001 | 0.160 |
| | | | | clay | -0.080 | 0.049 | -1.630 | 0.103 | -0.177 | 0.016 |
| | | | | packm | -0.031 | 0.055 | -0.563 | 0.574 | -0.139 | 0.077 |
| | | | | Slope | 0.187 | 0.049 | 3.775 | < **0.001** | 0.090 | 0.284 |

| Species | N | Formula | $R^2$ | Covariate | Est. | SE | *z* | *p* | *lower 95% CI* | *upper 95% CI* |
|---|---|---|---|---|---|---|---|---|---|---|
| | | | | Ydays | -0.001 | 0.001 | -1.117 | 0.264 | -0.003 | 0.001 |
| *Balsamorhiza hookeri* | 24 | Seed mass ~ DeltSpringMax + SoilpH + AnnMinTemp | 0.460 | (Intercept) | 1.717 | 0.045 | 37.770 | < **0.001** | 1.628 | 1.806 |
| | | | | AnnMinTemp | -0.029 | 0.037 | -0.776 | 0.438 | -0.102 | 0.044 |
| | | | | DeltSpringMax | 0.035 | 0.037 | 0.934 | 0.350 | -0.038 | 0.108 |
| | | | | SoilpH | 0.109 | 0.037 | 2.913 | **0.004** | 0.036 | 0.183 |
| *Balsamorhiza sagittata* | 64 | Seed mass ~ DeltPrecip | 0.108 | (Intercept) | 2.229 | 0.036 | 61.500 | < **0.001** | 2.158 | 2.300 |
| | | | | DeltPrecip | 0.051 | 0.031 | 1.668 | 0.095 | -0.009 | 0.112 |
| *Bebbia juncea* | 21 | Seed mass ~ Slope + Ydays | 0.272 | (Intercept) | -0.450 | 0.041 | -11.107 | < **0.001** | -0.530 | -0.371 |
| | | | | Slope | -0.060 | 0.030 | -1.994 | **0.046** | -0.120 | -0.001 |

| Species | N | Formula | $R^2$ | Covariate | Est. | SE | *z* | *p* | *lower 95% CI* | *upper 95% CI* |
|---|---|---|---|---|---|---|---|---|---|---|
| | | | | Ydays | -0.002 | 0.001 | -2.538 | **0.011** | -0.003 | 0.000 |
| *Bouteloua barbata* | 25 | Seed mass ~ heatload + clay | 0.308 | (Intercept) | -2.149 | 0.032 | -66.575 | < **0.001** | -2.213 | -2.086 |
| | | | | clay | 0.121 | 0.036 | 3.387 | **0.001** | 0.051 | 0.190 |
| | | | | heatload | 0.165 | 0.039 | 4.206 | < **0.001** | 0.088 | 0.243 |
| *Bouteloua breviseta* | 24 | Seed mass ~ packm + Ydays + SixMonthPrecip + DeltFallMin + DeltFallMax + DeltSpringMax | 0.658 | (Intercept) | -0.916 | 0.023 | -39.024 | < **0.001** | -0.962 | -0.870 |
| | | | | DeltFallMax | 0.078 | 0.123 | 0.633 | 0.527 | -0.163 | 0.319 |

| Species | N | Formula | $R^2$ | Covariate | Est. | SE | *z* | *p* | *lower 95% CI* | *upper 95% CI* |
|---|---|---|---|---|---|---|---|---|---|---|
| | | | | DeltFallMin | 0.191 | 0.140 | 1.361 | 0.173 | -0.084 | 0.466 |
| | | | | DeltSpringMax | -0.187 | 0.059 | -3.183 | **0.001** | -0.302 | -0.072 |
| | | | | packm | -0.162 | 0.233 | -0.693 | 0.488 | -0.619 | 0.296 |
| | | | | SixMonthPrecip | 0.122 | 0.051 | 2.381 | **0.017** | 0.022 | 0.222 |
| | | | | Ydays | 0.016 | 0.006 | 2.814 | **0.005** | 0.005 | 0.028 |
| *Bouteloua curtipendula* | 82 | Seed mass ~ Year + packm + DeltFallMax + DeltPrecip | 0.623 | (Intercept) | 0.475 | 0.050 | 9.575 | < **0.001** | 0.378 | 0.572 |
| | | | | DeltFallMax | 0.030 | 0.074 | 0.409 | 0.683 | -0.116 | 0.176 |
| | | | | DeltPrecip | -0.133 | 0.074 | -1.784 | 0.074 | -0.278 | 0.013 |
| | | | | packm | -0.098 | 0.044 | -2.258 | **0.024** | -0.184 | -0.013 |
| | | | | Year | 0.147 | 0.016 | 9.018 | < **0.001** | 0.115 | 0.179 |

| Species | N | Formula | $R^2$ | Covariate | Est. | SE | *z* | *p* | *lower 95% CI* | *upper 95% CI* |
|---|---|---|---|---|---|---|---|---|---|---|
| *Bouteloua gracilis* | 131 | Seed mass ~ Year + Ydays + DeltSpringMax | 0.125 | (Intercept) | -0.752 | 0.030 | -25.217 | < **0.001** | -0.811 | -0.694 |
| | | | | DeltSpringMax | 0.003 | 0.032 | 0.090 | 0.928 | -0.060 | 0.065 |
| | | | | Ydays | 0.002 | 0.001 | 2.334 | **0.020** | 0.000 | 0.004 |
| | | | | Year | 0.010 | 0.009 | 1.156 | 0.248 | -0.007 | 0.028 |
| *Bouteloua hirsuta* | 31 | Seed mass ~ Year | 0.018 | (Intercept) | -0.481 | 0.042 | -11.376 | < **0.001** | -0.564 | -0.398 |
| | | | | Year | -0.008 | 0.012 | -0.658 | 0.511 | -0.031 | 0.016 |
| *Camissonia brevipes* | 28 | Seed mass ~ Year | 0.005 | (Intercept) | -0.893 | 0.086 | -10.421 | < **0.001** | -1.061 | -0.725 |
| | | | | Year | -0.011 | 0.030 | -0.370 | 0.712 | -0.069 | 0.047 |
| *Cercocarpus ledifolius* | 63 | Seed mass ~ Year + Aspect | 0.220 | (Intercept) | 2.357 | 0.020 | 115.793 | < **0.001** | 2.317 | 2.397 |

| Species | N | Formula | $R^2$ | Covariate | Est. | SE | *z* | *p* | *lower 95% CI* | *upper 95% CI* |
|---|---|---|---|---|---|---|---|---|---|---|
| | | + DeltFallMin + SoilpH + clay + organic + AnnMinTemp | | | | | | | | |
| | | | | AnnMinTemp | 0.004 | 0.023 | 0.176 | 0.860 | -0.040 | 0.048 |
| | | | | Aspect | -0.025 | 0.023 | -1.098 | 0.272 | -0.069 | 0.020 |
| | | | | clay | 0.042 | 0.025 | 1.662 | 0.097 | -0.008 | 0.091 |
| | | | | DeltFallMin | 0.042 | 0.025 | 1.704 | 0.088 | -0.006 | 0.091 |
| | | | | organic | 0.011 | 0.025 | 0.435 | 0.664 | -0.039 | 0.061 |
| | | | | SoilpH | -0.040 | 0.024 | -1.623 | 0.105 | -0.087 | 0.008 |
| | | | | Year | 0.002 | 0.009 | 0.229 | 0.819 | -0.015 | 0.019 |
| *Cercocarpus montanus* | 26 | Seed mass ~ DeltPrecip | 0.339 | (Intercept) | 2.520 | 0.051 | 49.206 | < **0.001** | 2.420 | 2.621 |
| | | | | DeltPrecip | 0.066 | 0.041 | 1.622 | 0.105 | -0.014 | 0.146 |

| Species | N | Formula | $R^2$ | Covariate | Est. | SE | *z* | *p* | *lower 95% CI* | *upper 95% CI* |
|---|---|---|---|---|---|---|---|---|---|---|
| *Chaenactis fremontii* | 43 | Seed mass ~ Slope + SixMonthPrecip + SoilpH | 0.333 | (Intercept) | -0.602 | 0.027 | -21.948 | < **0.001** | -0.656 | -0.549 |
| | | | | SixMonthPrecip | 0.064 | 0.033 | 1.916 | 0.055 | -0.001 | 0.129 |
| | | | | Slope | -0.111 | 0.033 | -3.352 | **0.001** | -0.176 | -0.046 |
| | | | | SoilpH | -0.106 | 0.035 | -3.014 | **0.003** | -0.175 | -0.037 |
| *Chaenactis stevioides* | 29 | Seed mass ~ SoilpH | 0.206 | (Intercept) | -0.969 | 0.020 | -48.471 | < **0.001** | -1.008 | -0.929 |
| | | | | SoilpH | 0.065 | 0.021 | 3.044 | **0.002** | 0.023 | 0.107 |
| *Chrysothamnus viscidiflorus* | 67 | Seed mass ~ DeltPrecip + clay | 0.109 | (Intercept) | -0.747 | 0.032 | -23.339 | < **0.001** | -0.810 | -0.684 |
| | | | | clay | 0.075 | 0.030 | 2.494 | **0.013** | 0.016 | 0.135 |

| Species | N | Formula | $R^2$ | Covariate | Est. | SE | *z* | *p* | *lower 95% CI* | *upper 95% CI* |
|---|---|---|---|---|---|---|---|---|---|---|
| | | | | DeltPrecip | 0.055 | 0.031 | 1.797 | 0.072 | -0.005 | 0.115 |
| *Cleome lutea* | 46 | Seed mass ~ Aspect + SoilpH | 0.368 | (Intercept) | 1.480 | 0.023 | 63.829 | < **0.001** | 1.434 | 1.525 |
| | | | | Aspect | 0.046 | 0.016 | 2.919 | **0.004** | 0.015 | 0.076 |
| | | | | SoilpH | -0.041 | 0.020 | -2.118 | **0.034** | -0.080 | -0.003 |
| *Cleome serrulata* | 40 | Seed mass ~ Aspect + SoilpH | 0.261 | (Intercept) | 1.878 | 0.028 | 68.253 | < **0.001** | 1.824 | 1.932 |
| | | | | Aspect | 0.033 | 0.018 | 1.839 | 0.066 | -0.002 | 0.067 |
| | | | | SoilpH | -0.014 | 0.017 | -0.835 | 0.404 | -0.047 | 0.019 |
| *Crepis acuminata* | 43 | Seed mass ~ Ydays | 0.058 | (Intercept) | 1.028 | 0.045 | 22.747 | < **0.001** | 0.940 | 1.117 |
| | | | | Ydays | 0.000 | 0.002 | -0.127 | 0.899 | -0.003 | 0.003 |

| Species | N | Formula | $R^2$ | Covariate | Est. | SE | *z* | *p* | *lower 95% CI* | *upper 95% CI* |
|---|---|---|---|---|---|---|---|---|---|---|
| *Elymus elymoides* | 184 | Seed mass ~ Slope + Ydays + SixMonthPrecip + SoilpH + organic + AnnMinTemp | 0.396 | (Intercept) | 0.959 | 0.021 | 46.568 | < **0.001** | 0.919 | 1.000 |
| | | | | AnnMinTemp | 0.070 | 0.024 | 2.905 | **0.004** | 0.023 | 0.117 |
| | | | | organic | -0.003 | 0.024 | -0.119 | 0.905 | -0.050 | 0.044 |
| | | | | SixMonthPrecip | 0.034 | 0.024 | 1.412 | 0.158 | -0.013 | 0.080 |
| | | | | Slope | -0.002 | 0.022 | -0.082 | 0.935 | -0.045 | 0.041 |
| | | | | SoilpH | -0.141 | 0.026 | -5.420 | < **0.001** | -0.192 | -0.090 |
| | | | | Ydays | 0.004 | 0.001 | 3.608 | < **0.001** | 0.002 | 0.006 |

| Species | N | Formula | $R^2$ | Covariate | Est. | SE | *z* | *p* | *lower 95% CI* | *upper 95% CI* |
|---|---|---|---|---|---|---|---|---|---|---|
| *Elymus glaucus* | 21 | Seed mass ~ Slope | 0.040 | (Intercept) | 1.341 | 0.097 | 13.876 | < **0.001** | 1.152 | 1.531 |
| | | | | Slope | 0.078 | 0.101 | 0.765 | 0.444 | -0.121 | 0.276 |
| *Encelia farinosa* | 40 | Seed mass ~ SoilpH + organic | 0.096 | (Intercept) | 0.213 | 0.029 | 7.417 | < **0.001** | 0.157 | 0.270 |
| | | | | organic | -0.045 | 0.045 | -0.996 | 0.319 | -0.133 | 0.043 |
| | | | | SoilpH | -0.081 | 0.046 | -1.750 | 0.080 | -0.171 | 0.010 |
| *Encelia virginensis* | 25 | Seed mass ~ Ydays + AnnMinTemp | 0.323 | (Intercept) | 0.768 | 0.062 | 12.408 | < **0.001** | 0.646 | 0.889 |
| | | | | AnnMinTemp | -0.103 | 0.053 | -1.937 | 0.053 | -0.208 | 0.001 |
| | | | | Ydays | 0.000 | 0.002 | 0.171 | 0.864 | -0.003 | 0.003 |

| Species | N | Formula | $R^2$ | Covariate | Est. | SE | *z* | *p* | *lower 95% CI* | *upper 95% CI* |
|---|---|---|---|---|---|---|---|---|---|---|
| *Ericameria nauseosa* | 88 | Seed mass ~ Year + DeltFallMax + SoilpH + organic | 0.413 | (Intercept) | -0.374 | 0.051 | -7.333 | < **0.001** | -0.473 | -0.274 |
| | | | | DeltFallMax | 0.059 | 0.044 | 1.341 | 0.180 | -0.027 | 0.145 |
| | | | | organic | 0.090 | 0.040 | 2.267 | **0.023** | 0.012 | 0.168 |
| | | | | SoilpH | -0.173 | 0.044 | -3.945 | < **0.001** | -0.259 | -0.087 |
| | | | | Year | -0.060 | 0.015 | -4.052 | < **0.001** | -0.090 | -0.031 |
| *Erigeron pumilus* | 25 | Seed mass ~ Year + DeltFallMax + SoilpH + organic | 0.429 | (Intercept) | -2.473 | 0.095 | -26.132 | < **0.001** | -2.658 | -2.287 |

| Species | N | Formula | $R^2$ | Covariate | Est. | SE | *z* | *p* | *lower* *95% CI* | *upper* *95% CI* |
|---|---|---|---|---|---|---|---|---|---|---|
| | | | | DeltFallMax | 0.316 | 0.112 | 2.819 | **0.005** | 0.096 | 0.536 |
| | | | | organic | -0.134 | 0.120 | -1.114 | 0.265 | -0.369 | 0.102 |
| | | | | SoilpH | -0.157 | 0.091 | -1.723 | 0.085 | -0.336 | 0.022 |
| | | | | Year | -0.035 | 0.017 | -2.079 | **0.038** | -0.067 | -0.002 |
| *Eriogonum heracleoides* | 41 | Seed mass ~ Slope + SixMonthPrecip + DeltSpringMax | 0.414 | (Intercept) | 0.891 | 0.030 | 29.355 | < **0.001** | 0.832 | 0.951 |
| | | | | DeltSpringMax | 0.073 | 0.030 | 2.412 | **0.016** | 0.014 | 0.132 |
| | | | | SixMonthPrecip | 0.070 | 0.032 | 2.204 | **0.028** | 0.008 | 0.133 |
| | | | | Slope | 0.050 | 0.032 | 1.585 | 0.113 | -0.012 | 0.112 |
| *Eriogonum umbellatum* | 70 | Seed mass ~ heatload + | 0.465 | (Intercept) | 0.898 | 0.043 | 21.011 | < **0.001** | 0.814 | 0.982 |

| Species | N | Formula | $R^2$ | Covariate | Est. | SE | *z* | *p* | *lower 95% CI* | *upper 95% CI* |
|---|---|---|---|---|---|---|---|---|---|---|
| | | Ydays + DeltSpringMax + SoilpH + clay + AnnMinTemp | | | | | | | | |
| | | | | AnnMinTemp | 0.051 | 0.035 | 1.454 | 0.146 | -0.018 | 0.120 |
| | | | | clay | 0.030 | 0.026 | 1.147 | 0.251 | -0.021 | 0.081 |
| | | | | DeltSpringMax | 0.009 | 0.035 | 0.245 | 0.806 | -0.061 | 0.078 |
| | | | | heatload | 0.015 | 0.026 | 0.560 | 0.576 | -0.037 | 0.067 |
| | | | | SoilpH | -0.048 | 0.030 | -1.583 | 0.114 | -0.107 | 0.011 |
| | | | | Ydays | 0.002 | 0.001 | 1.384 | 0.166 | -0.001 | 0.004 |
| *Eriophyllum lanatum* | 39 | Seed mass ~ Ydays + AWC + SoilpH | 0.503 | (Intercept) | -0.752 | 0.055 | -13.729 | < **0.001** | -0.860 | -0.645 |
| | | | | AWC | 0.076 | 0.033 | 2.340 | **0.019** | 0.012 | 0.140 |

| Species | N | Formula | $R^2$ | Covariate | Est. | SE | *z* | *p* | *lower 95% CI* | *upper 95% CI* |
|---|---|---|---|---|---|---|---|---|---|---|
| | | | | SoilpH | 0.085 | 0.049 | 1.713 | 0.087 | -0.012 | 0.181 |
| | | | | Ydays | -0.003 | 0.002 | -1.486 | 0.137 | -0.008 | 0.001 |
| *Fallugia paradoxa* | 46 | Seed mass ~ Aspect + Ydays + AWC + SoilpH + clay + organic + AnnMinTemp | 0.575 | (Intercept) | -0.231 | 0.051 | -4.569 | **< 0.001** | -0.330 | -0.132 |
| | | | | AnnMinTemp | -0.096 | 0.046 | -2.117 | **0.034** | -0.186 | -0.007 |
| | | | | Aspect | 0.016 | 0.026 | 0.614 | 0.539 | -0.035 | 0.068 |
| | | | | AWC | -0.059 | 0.043 | -1.382 | 0.167 | -0.142 | 0.025 |
| | | | | clay | -0.048 | 0.035 | -1.363 | 0.173 | -0.118 | 0.021 |
| | | | | organic | -0.059 | 0.030 | -1.945 | 0.052 | -0.119 | 0.000 |
| | | | | SoilpH | -0.038 | 0.031 | -1.225 | 0.221 | -0.098 | 0.023 |

| Species | N | Formula | $R^2$ | Covariate | Est. | SE | *z* | *p* | *lower 95% CI* | *upper 95% CI* |
|---|---|---|---|---|---|---|---|---|---|---|
| | | | | Ydays | -0.001 | 0.001 | -1.648 | 0.099 | -0.003 | 0.000 |
| *Festuca idahoensis* | 31 | Seed mass ~ Aspect + Ydays + AWC + SoilpH + clay + organic + AnnMinTemp | 0.473 | (Intercept) | 0.186 | 0.022 | 8.488 | < **0.001** | 0.143 | 0.229 |
| | | | | AnnMinTemp | -0.014 | 0.036 | -0.374 | 0.708 | -0.085 | 0.058 |
| | | | | Aspect | 0.019 | 0.030 | 0.655 | 0.513 | -0.039 | 0.077 |
| | | | | AWC | 0.090 | 0.041 | 2.219 | **0.026** | 0.011 | 0.169 |
| | | | | clay | 0.050 | 0.032 | 1.559 | 0.119 | -0.013 | 0.114 |
| | | | | organic | -0.116 | 0.045 | -2.584 | **0.010** | -0.205 | -0.028 |
| | | | | SoilpH | -0.103 | 0.039 | -2.610 | **0.009** | -0.180 | -0.026 |
| | | | | Ydays | 0.000 | 0.002 | 0.161 | 0.872 | -0.004 | 0.004 |

| Species | N | Formula | $R^2$ | Covariate | Est. | SE | *z* | *p* | *lower 95% CI* | *upper 95% CI* |
|---|---|---|---|---|---|---|---|---|---|---|
| *Grayia spinosa* | 49 | Seed mass ~ SixMonthPrecip + DeltFallMax | 0.194 | (Intercept) | -0.075 | 0.018 | -4.118 | < **0.001** | -0.111 | -0.039 |
| | | | | DeltFallMax | -0.043 | 0.022 | -1.960 | **0.050** | -0.085 | 0.000 |
| | | | | SixMonthPrecip | 0.037 | 0.021 | 1.782 | 0.075 | -0.004 | 0.078 |
| *Helianthus annuus* | 37 | Seed mass ~ heatload + SixMonthPrecip + SoilpH + organic | 0.514 | (Intercept) | 1.822 | 0.016 | 115.695 | < **0.001** | 1.791 | 1.852 |
| | | | | heatload | -0.068 | 0.017 | -3.976 | < **0.001** | -0.101 | -0.034 |
| | | | | organic | 0.031 | 0.017 | 1.765 | 0.078 | -0.003 | 0.065 |
| | | | | SixMonthPrecip | 0.050 | 0.018 | 2.793 | **0.005** | 0.015 | 0.085 |
| | | | | SoilpH | -0.037 | 0.019 | -1.983 | **0.047** | -0.073 | 0.000 |

| Species | N | Formula | $R^2$ | Covariate | Est. | SE | *z* | *p* | *lower 95% CI* | *upper 95% CI* |
|---|---|---|---|---|---|---|---|---|---|---|
| *Heliomeris multiflora* | 53 | Seed mass ~ Ydays + SixMonthPrecip | 0.234 | (Intercept) | -0.622 | 0.046 | -13.411 | < **0.001** | -0.712 | -0.531 |
| | | | | SixMonthPrecip | 0.058 | 0.062 | 0.941 | 0.347 | -0.063 | 0.179 |
| | | | | Ydays | -0.005 | 0.001 | -3.673 | < **0.001** | -0.008 | -0.002 |
| *Hesperostipa comata* | 144 | Seed mass ~ Year + heatload + DeltPrecip + AnnMinTemp | 0.240 | (Intercept) | 1.826 | 0.028 | 65.908 | < **0.001** | 1.772 | 1.881 |
| | | | | AnnMinTemp | 0.040 | 0.026 | 1.530 | 0.126 | -0.011 | 0.090 |
| | | | | DeltPrecip | 0.081 | 0.028 | 2.896 | **0.004** | 0.026 | 0.137 |
| | | | | heatload | 0.032 | 0.021 | 1.520 | 0.128 | -0.009 | 0.073 |
| | | | | Year | 0.024 | 0.009 | 2.664 | **0.008** | 0.006 | 0.042 |

| Species | N | Formula | $R^2$ | Covariate | Est. | SE | *z* | *p* | *lower 95% CI* | *upper 95% CI* |
|---|---|---|---|---|---|---|---|---|---|---|
| *Heterotheca villosa* | 73 | Seed mass ~ Ydays + SixMonthPrecip + DeltFallMax + DeltSpringMax | 0.311 | (Intercept) | -0.745 | 0.037 | -20.390 | < **0.001** | -0.817 | -0.673 |
| | | | | DeltFallMax | -0.065 | 0.039 | -1.661 | 0.097 | -0.142 | 0.012 |
| | | | | DeltSpringMax | -0.056 | 0.037 | -1.497 | 0.134 | -0.129 | 0.017 |
| | | | | SixMonthPrecip | 0.149 | 0.036 | 4.100 | < **0.001** | 0.078 | 0.220 |
| | | | | Ydays | 0.001 | 0.001 | 0.905 | 0.365 | -0.001 | 0.004 |
| *Hymenoclea salsola* | 21 | Seed mass ~ Ydays + SixMonthPrecip + DeltFallMax | 0.561 | (Intercept) | 1.726 | 0.043 | 39.960 | < **0.001** | 1.641 | 1.810 |

| Species | N | Formula | $R^2$ | Covariate | Est. | SE | *z* | *p* | *lower 95% CI* | *upper 95% CI* |
|---|---|---|---|---|---|---|---|---|---|---|
| | | + DeltSpringMax | | | | | | | | |
| | | | | DeltFallMax | -0.114 | 0.022 | -5.096 | < **0.001** | -0.158 | -0.070 |
| | | | | DeltSpringMax | 0.086 | 0.025 | 3.479 | **0.001** | 0.037 | 0.134 |
| | | | | SixMonthPrecip | -0.106 | 0.029 | -3.655 | < **0.001** | -0.163 | -0.049 |
| | | | | Ydays | 0.006 | 0.002 | 3.712 | < **0.001** | 0.003 | 0.009 |
| *Iris missouriensis* | 22 | Seed mass ~ heatload + DeltSpringMax + DeltPrecip | 0.465 | (Intercept) | 2.708 | 0.088 | 30.600 | < **0.001** | 2.535 | 2.881 |
| | | | | DeltPrecip | -0.035 | 0.079 | -0.438 | 0.661 | -0.190 | 0.121 |
| | | | | DeltSpringMax | 0.010 | 0.082 | 0.117 | 0.907 | -0.152 | 0.171 |
| | | | | heatload | 0.263 | 0.084 | 3.132 | **0.002** | 0.099 | 0.428 |

| Species | N | Formula | $R^2$ | Covariate | Est. | SE | *z* | *p* | *lower 95% CI* | *upper 95% CI* |
|---|---|---|---|---|---|---|---|---|---|---|
| *Koeleria macrantha* | 34 | Seed mass ~ DeltSpringMax + DeltPrecip | 0.257 | (Intercept) | -1.217 | 0.041 | -29.625 | < **0.001** | -1.298 | -1.137 |
| | | | | DeltPrecip | -0.156 | 0.054 | -2.870 | **0.004** | -0.263 | -0.050 |
| | | | | DeltSpringMax | 0.020 | 0.054 | 0.363 | 0.717 | -0.087 | 0.127 |
| *Krascheninnikovia lanata* | 80 | Seed mass ~ Slope + packm + Ydays + SoilpH + AnnMinTemp | 0.422 | (Intercept) | 0.739 | 0.029 | 25.259 | < **0.001** | 0.681 | 0.796 |
| | | | | AnnMinTemp | 0.103 | 0.037 | 2.821 | **0.005** | 0.031 | 0.175 |
| | | | | packm | 0.007 | 0.031 | 0.210 | 0.833 | -0.054 | 0.067 |
| | | | | Slope | 0.058 | 0.026 | 2.196 | **0.028** | 0.006 | 0.109 |
| | | | | SoilpH | -0.085 | 0.028 | -3.052 | **0.002** | -0.139 | -0.030 |

| Species | N | Formula | $R^2$ | Covariate | Est. | SE | *z* | *p* | *lower 95% CI* | *upper 95% CI* |
|---|---|---|---|---|---|---|---|---|---|---|
| | | | | Ydays | 0.000 | 0.001 | -0.645 | 0.519 | -0.002 | 0.001 |
| *Larrea tridentata* | 74 | Seed mass ~ packm + SixMonthPrecip + AnnMinTemp | 0.098 | (Intercept) | 1.389 | 0.056 | 24.709 | < **0.001** | 1.279 | 1.499 |
| | | | | AnnMinTemp | -0.091 | 0.062 | -1.469 | 0.142 | -0.212 | 0.030 |
| | | | | packm | 0.084 | 0.064 | 1.310 | 0.190 | -0.042 | 0.209 |
| | | | | SixMonthPrecip | 0.059 | 0.061 | 0.962 | 0.336 | -0.061 | 0.178 |
| *Leymus cinereus* | 155 | Seed mass ~ Year + DeltSpringMax + SoilpH + AnnMinTemp | 0.174 | (Intercept) | 1.207 | 0.015 | 79.781 | < **0.001** | 1.177 | 1.236 |
| | | | | AnnMinTemp | 0.030 | 0.015 | 1.989 | **0.047** | 0.000 | 0.059 |

| Species | N | Formula | $R^2$ | Covariate | Est. | SE | *z* | *p* | *lower 95% CI* | *upper 95% CI* |
|---|---|---|---|---|---|---|---|---|---|---|
| | | | | DeltSpringMax | -0.024 | 0.015 | -1.591 | 0.112 | -0.054 | 0.006 |
| | | | | SoilpH | -0.023 | 0.014 | -1.663 | 0.096 | -0.051 | 0.004 |
| | | | | Year | -0.005 | 0.004 | -1.384 | 0.166 | -0.013 | 0.002 |
| *Linum lewisii* | 31 | Seed mass ~ packm + heatload + AWC + AnnMinTemp | 0.329 | (Intercept) | 0.585 | 0.054 | 10.819 | < **0.001** | 0.479 | 0.691 |
| | | | | AnnMinTemp | 0.112 | 0.057 | 1.968 | **0.049** | 0.000 | 0.223 |
| | | | | AWC | -0.127 | 0.058 | -2.180 | **0.029** | -0.242 | -0.013 |
| | | | | heatload | 0.020 | 0.055 | 0.373 | 0.709 | -0.087 | 0.128 |
| | | | | packm | 0.026 | 0.059 | 0.448 | 0.654 | -0.089 | 0.142 |
| *Lomatium nudicaule* | 23 | Seed mass ~ SoilpH | 0.351 | (Intercept) | 2.573 | 0.092 | 27.949 | < **0.001** | 2.392 | 2.753 |

| Species | N | Formula | $R^2$ | Covariate | Est. | SE | *z* | *p* | *lower 95% CI* | *upper 95% CI* |
|---|---|---|---|---|---|---|---|---|---|---|
| | | | | SoilpH | -0.195 | 0.085 | -2.310 | **0.021** | -0.361 | -0.030 |
| *Lomatium triternatum* | 29 | Seed mass ~ Slope + heatload + DeltFallMin + DeltFallMax | 0.521 | (Intercept) | 2.117 | 0.033 | 64.791 | < **0.001** | 2.053 | 2.181 |
| | | | | DeltFallMax | 0.116 | 0.045 | 2.562 | **0.010** | 0.027 | 0.205 |
| | | | | DeltFallMin | 0.072 | 0.039 | 1.842 | 0.065 | -0.005 | 0.148 |
| | | | | heatload | 0.000 | 0.076 | 0.002 | 0.999 | -0.149 | 0.150 |
| | | | | Slope | -0.129 | 0.081 | -1.601 | 0.109 | -0.287 | 0.029 |
| *Lupinus argenteus* | 21 | Seed mass ~ Ydays | 0.104 | (Intercept) | 3.149 | 0.052 | 60.176 | < **0.001** | 3.047 | 3.252 |
| | | | | Ydays | -0.003 | 0.002 | -1.574 | 0.115 | -0.007 | 0.001 |

| Species | N | Formula | $R^2$ | Covariate | Est. | SE | *z* | *p* | *lower 95% CI* | *upper 95% CI* |
|---|---|---|---|---|---|---|---|---|---|---|
| *Machaeranthera canescens* | 63 | Seed mass ~ Slope + heatload + Ydays + SixMonthPrecip + DeltFallMax | 0.400 | (Intercept) | -0.861 | 0.055 | -15.752 | < **0.001** | -0.968 | -0.754 |
| | | | | DeltFallMax | -0.052 | 0.043 | -1.206 | 0.228 | -0.136 | 0.032 |
| | | | | heatload | -0.101 | 0.058 | -1.727 | 0.084 | -0.215 | 0.014 |
| | | | | SixMonthPrecip | 0.110 | 0.048 | 2.275 | **0.023** | 0.015 | 0.205 |
| | | | | Slope | 0.004 | 0.052 | 0.080 | 0.936 | -0.098 | 0.106 |
| | | | | Ydays | -0.002 | 0.001 | -1.289 | 0.198 | -0.005 | 0.001 |
| *Machaeranthera pinnatifida* | 29 | Seed mass ~ heatload + Ydays | 0.426 | (Intercept) | -1.198 | 0.039 | -31.065 | < **0.001** | -1.274 | -1.122 |

| Species | N | Formula | $R^2$ | Covariate | Est. | SE | *z* | *p* | *lower 95% CI* | *upper 95% CI* |
|---|---|---|---|---|---|---|---|---|---|---|
| | | | | heatload | -0.201 | 0.044 | -4.572 | < **0.001** | -0.287 | -0.115 |
| | | | | Ydays | -0.001 | 0.001 | -1.463 | 0.144 | -0.003 | 0.000 |
| *Machaeranthera tanacetifolia* | 40 | Seed mass ~ packm + heatload | 0.092 | (Intercept) | -0.197 | 0.033 | -6.001 | < **0.001** | -0.262 | -0.133 |
| | | | | heatload | -0.029 | 0.034 | -0.849 | 0.396 | -0.096 | 0.038 |
| | | | | packm | 0.067 | 0.035 | 1.901 | 0.057 | -0.002 | 0.135 |
| *Malacothrix glabrata* | 32 | Seed mass ~ clay | 0.172 | (Intercept) | -1.606 | 0.033 | -48.727 | < **0.001** | -1.670 | -1.541 |
| | | | | clay | -0.091 | 0.034 | -2.663 | **0.008** | -0.158 | -0.024 |
| *Mentzelia albicaulis* | 38 | Seed mass ~ heatload + DeltFallMin + AWC + clay + | 0.580 | (Intercept) | -1.317 | 0.039 | -33.794 | < **0.001** | -1.393 | -1.240 |

| Species | N | Formula | $R^2$ | Covariate | Est. | SE | *z* | *p* | *lower 95% CI* | *upper 95% CI* |
|---|---|---|---|---|---|---|---|---|---|---|
| | | organic + AnnMinTemp | | | | | | | | |
| | | | | AnnMinTemp | 0.011 | 0.043 | 0.254 | 0.800 | -0.073 | 0.094 |
| | | | | AWC | -0.001 | 0.029 | -0.033 | 0.974 | -0.057 | 0.055 |
| | | | | clay | 0.031 | 0.036 | 0.882 | 0.378 | -0.038 | 0.101 |
| | | | | DeltFallMin | -0.036 | 0.031 | -1.156 | 0.248 | -0.096 | 0.025 |
| | | | | heatload | -0.060 | 0.036 | -1.679 | 0.093 | -0.131 | 0.010 |
| | | | | organic | 0.057 | 0.038 | 1.504 | 0.133 | -0.017 | 0.131 |
| *Mentzelia laevicaulis* | 30 | Seed mass ~ heatload + DeltFallMin + AWC + clay + organic + AnnMinTemp | 0.331 | (Intercept) | 0.121 | 0.011 | 11.430 | < **0.001** | 0.100 | 0.142 |

| Species | N | Formula | $R^2$ | Covariate | Est. | SE | *z* | *p* | *lower 95% CI* | *upper 95% CI* |
|---|---|---|---|---|---|---|---|---|---|---|
| | | | | AnnMinTemp | 0.016 | 0.014 | 1.160 | 0.246 | -0.011 | 0.042 |
| | | | | AWC | 0.004 | 0.015 | 0.245 | 0.806 | -0.027 | 0.034 |
| | | | | clay | -0.031 | 0.022 | -1.413 | 0.158 | -0.075 | 0.012 |
| | | | | DeltFallMin | -0.026 | 0.013 | -1.988 | **0.047** | -0.052 | 0.000 |
| | | | | heatload | -0.023 | 0.014 | -1.649 | 0.099 | -0.050 | 0.004 |
| | | | | organic | 0.047 | 0.018 | 2.569 | **0.010** | 0.011 | 0.082 |
| *Muhlenbergia porteri* | 35 | Seed mass ~ heatload + DeltFallMin + AWC + clay + organic + AnnMinTemp | 0.368 | (Intercept) | -1.817 | 0.031 | -57.674 | < **0.001** | -1.878 | -1.755 |
| | | | | AnnMinTemp | -0.053 | 0.037 | -1.456 | 0.145 | -0.125 | 0.018 |
| | | | | AWC | -0.037 | 0.040 | -0.934 | 0.350 | -0.116 | 0.041 |

| Species | N | Formula | $R^2$ | Covariate | Est. | SE | *z* | *p* | *lower 95% CI* | *upper 95% CI* |
|---|---|---|---|---|---|---|---|---|---|---|
| | | | | clay | 0.068 | 0.059 | 1.153 | 0.249 | -0.047 | 0.182 |
| | | | | DeltFallMin | 0.111 | 0.038 | 2.879 | **0.004** | 0.035 | 0.186 |
| | | | | heatload | -0.074 | 0.041 | -1.816 | 0.069 | -0.153 | 0.006 |
| | | | | organic | -0.160 | 0.054 | -2.969 | **0.003** | -0.266 | -0.055 |
| *Nicotiana attenuata* | 30 | Seed mass ~ heatload + DeltFallMin + AWC + clay + organic + AnnMinTemp | 0.516 | (Intercept) | -2.021 | 0.009 | -235.808 | < **0.001** | -2.037 | -2.004 |
| | | | | AnnMinTemp | -0.025 | 0.010 | -2.459 | **0.014** | -0.044 | -0.005 |
| | | | | AWC | 0.018 | 0.016 | 1.119 | 0.263 | -0.013 | 0.049 |
| | | | | clay | -0.010 | 0.014 | -0.706 | 0.480 | -0.038 | 0.018 |
| | | | | DeltFallMin | 0.040 | 0.011 | 3.633 | < **0.001** | 0.019 | 0.062 |

| Species | N | Formula | $R^2$ | Covariate | Est. | SE | *z* | *p* | *lower 95% CI* | *upper 95% CI* |
|---|---|---|---|---|---|---|---|---|---|---|
| | | | | heatload | 0.023 | 0.012 | 1.998 | **0.046** | 0.000 | 0.046 |
| | | | | organic | -0.056 | 0.013 | -4.349 | < **0.001** | -0.081 | -0.031 |
| *Pectis papposa* | 31 | Seed mass ~ Year + packm + SoilpH | 0.580 | (Intercept) | -1.856 | 0.028 | -65.808 | < **0.001** | -1.911 | -1.801 |
| | | | | packm | 0.049 | 0.025 | 1.957 | 0.050 | 0.000 | 0.098 |
| | | | | SoilpH | 0.027 | 0.025 | 1.065 | 0.287 | -0.023 | 0.077 |
| | | | | Year | 0.070 | 0.012 | 5.743 | < **0.001** | 0.046 | 0.094 |
| *Penstemon acuminatus* | 20 | Seed mass ~ Ydays + SixMonthPrecip | 0.481 | (Intercept) | -0.076 | 0.024 | -3.181 | **0.001** | -0.124 | -0.029 |
| | | | | SixMonthPrecip | 0.015 | 0.026 | 0.568 | 0.570 | -0.036 | 0.065 |
| | | | | Ydays | 0.007 | 0.002 | 3.451 | **0.001** | 0.003 | 0.011 |

| Species | N | Formula | $R^2$ | Covariate | Est. | SE | *z* | *p* | *lower 95% CI* | *upper 95% CI* |
|---|---|---|---|---|---|---|---|---|---|---|
| *Penstemon deustus* | 30 | Seed mass ~ AWC + SoilpH + clay | 0.461 | (Intercept) | -2.013 | 0.044 | -45.277 | < **0.001** | -2.101 | -1.926 |
| | | | | AWC | 0.046 | 0.047 | 0.992 | 0.321 | -0.045 | 0.138 |
| | | | | clay | 0.040 | 0.048 | 0.823 | 0.410 | -0.055 | 0.135 |
| | | | | SoilpH | 0.078 | 0.041 | 1.915 | 0.055 | -0.002 | 0.159 |
| *Plantago ovata* | 44 | Seed mass ~ Slope + heatload + SoilpH + clay + AnnMinTemp | 0.476 | (Intercept) | 0.169 | 0.020 | 8.492 | < **0.001** | 0.130 | 0.208 |
| | | | | AnnMinTemp | -0.088 | 0.023 | -3.785 | < **0.001** | -0.134 | -0.042 |
| | | | | clay | -0.028 | 0.025 | -1.124 | 0.261 | -0.078 | 0.021 |
| | | | | heatload | -0.010 | 0.031 | -0.314 | 0.753 | -0.071 | 0.052 |

| Species | N | Formula | $R^2$ | Covariate | Est. | SE | *z* | *p* | *lower 95% CI* | *upper 95% CI* |
|---|---|---|---|---|---|---|---|---|---|---|
| | | | | Slope | 0.035 | 0.029 | 1.220 | 0.223 | -0.022 | 0.092 |
| | | | | SoilpH | 0.052 | 0.022 | 2.393 | **0.017** | 0.009 | 0.094 |
| *Plantago patagonica* | 60 | Seed mass ~ Year + Ydays + DeltPrecip + clay + organic + AnnMinTemp | 0.701 | (Intercept) | -0.047 | 0.018 | -2.590 | **0.010** | -0.082 | -0.011 |
| | | | | AnnMinTemp | 0.117 | 0.023 | 5.080 | < **0.001** | 0.072 | 0.161 |
| | | | | clay | 0.081 | 0.022 | 3.702 | < **0.001** | 0.038 | 0.124 |
| | | | | DeltPrecip | -0.001 | 0.023 | -0.047 | 0.962 | -0.046 | 0.044 |
| | | | | organic | 0.012 | 0.019 | 0.641 | 0.522 | -0.025 | 0.049 |
| | | | | Ydays | 0.000 | 0.001 | -0.440 | 0.660 | -0.002 | 0.002 |
| | | | | Year | -0.024 | 0.007 | -3.348 | **0.001** | -0.038 | -0.010 |

| Species | N | Formula | $R^2$ | Covariate | Est. | SE | *z* | *p* | *lower 95% CI* | *upper 95% CI* |
|---|---|---|---|---|---|---|---|---|---|---|
| *Pleuraphis jamesii* | 37 | Seed mass ~ Year + Ydays + DeltFallMin + DeltSpringMax | 0.579 | (Intercept) | 0.434 | 0.044 | 9.887 | < **0.001** | 0.348 | 0.521 |
| | | | | DeltFallMin | 0.140 | 0.043 | 3.250 | **0.001** | 0.056 | 0.225 |
| | | | | DeltSpringMax | -0.047 | 0.052 | -0.890 | 0.373 | -0.149 | 0.056 |
| | | | | Ydays | -0.001 | 0.001 | -1.030 | 0.303 | -0.003 | 0.001 |
| | | | | Year | 0.014 | 0.020 | 0.729 | 0.466 | -0.024 | 0.053 |
| *Pleuraphis mutica* | 20 | Seed mass ~ Year + DeltFallMax + DeltPrecip | 0.722 | (Intercept) | 0.204 | 0.039 | 5.265 | < **0.001** | 0.128 | 0.281 |
| | | | | DeltFallMax | -0.222 | 0.091 | -2.433 | **0.015** | -0.400 | -0.043 |
| | | | | DeltPrecip | 0.016 | 0.107 | 0.148 | 0.882 | -0.193 | 0.225 |

| Species | N | Formula | $R^2$ | Covariate | Est. | SE | *z* | *p* | *lower 95% CI* | *upper 95% CI* |
|---|---|---|---|---|---|---|---|---|---|---|
| | | | | Year | 0.061 | 0.018 | 3.474 | **0.001** | 0.027 | 0.096 |
| *Poa secunda* | 126 | Seed mass ~ Year + SixMonthPrecip + SoilpH + clay | 0.182 | (Intercept) | -0.718 | 0.028 | -25.929 | < **0.001** | -0.772 | -0.663 |
| | | | | clay | 0.058 | 0.028 | 2.092 | **0.036** | 0.004 | 0.113 |
| | | | | SixMonthPrecip | 0.086 | 0.031 | 2.787 | **0.005** | 0.026 | 0.147 |
| | | | | SoilpH | -0.051 | 0.031 | -1.675 | 0.094 | -0.111 | 0.009 |
| | | | | Year | 0.004 | 0.009 | 0.415 | 0.678 | -0.014 | 0.022 |
| *Potentilla glandulosa* | 24 | Seed mass ~ clay + AnnMinTemp | 0.413 | (Intercept) | -1.496 | 0.064 | -23.195 | < **0.001** | -1.622 | -1.369 |
| | | | | AnnMinTemp | 0.116 | 0.066 | 1.760 | 0.078 | -0.013 | 0.246 |
| | | | | clay | 0.188 | 0.064 | 2.913 | **0.004** | 0.061 | 0.314 |

| Species | N | Formula | $R^2$ | Covariate | Est. | SE | *z* | *p* | *lower 95% CI* | *upper 95% CI* |
|---|---|---|---|---|---|---|---|---|---|---|
| *Pseudoroegneria spicata* | 76 | Seed mass ~ DeltSpringMax + DeltPrecip + SoilpH + organic | 0.275 | (Intercept) | 1.166 | 0.019 | 59.800 | < **0.001** | 1.127 | 1.204 |
| | | | | DeltPrecip | 0.016 | 0.021 | 0.763 | 0.446 | -0.025 | 0.056 |
| | | | | DeltSpringMax | -0.051 | 0.020 | -2.500 | **0.012** | -0.091 | -0.011 |
| | | | | organic | -0.049 | 0.017 | -2.910 | **0.004** | -0.083 | -0.016 |
| | | | | SoilpH | -0.041 | 0.019 | -2.144 | **0.032** | -0.079 | -0.004 |
| *Purshia stansburiana* | 25 | Seed mass ~ DeltSpringMax + AWC + SoilpH + clay + organic | 0.716 | (Intercept) | 1.944 | 0.014 | 140.428 | < **0.001** | 1.917 | 1.971 |

| Species | N | Formula | $R^2$ | Covariate | Est. | SE | *z* | *p* | *lower 95% CI* | *upper 95% CI* |
|---|---|---|---|---|---|---|---|---|---|---|
| | | | | AWC | 0.061 | 0.021 | 2.905 | **0.004** | 0.020 | 0.103 |
| | | | | clay | 0.028 | 0.020 | 1.405 | 0.160 | -0.011 | 0.067 |
| | | | | DeltSpringMax | 0.005 | 0.020 | 0.222 | 0.824 | -0.035 | 0.044 |
| | | | | organic | -0.005 | 0.025 | -0.204 | 0.838 | -0.054 | 0.044 |
| | | | | SoilpH | -0.087 | 0.020 | -4.450 | < **0.001** | -0.125 | -0.049 |
| *Purshia tridentata* | 70 | Seed mass ~ packm + Ydays + SixMonthPrecip + DeltSpringMax + SoilpH + clay | 0.207 | (Intercept) | 3.142 | 0.018 | 172.335 | < **0.001** | 3.107 | 3.178 |
| | | | | clay | -0.009 | 0.024 | -0.356 | 0.722 | -0.056 | 0.039 |
| | | | | DeltSpringMax | 0.011 | 0.026 | 0.419 | 0.676 | -0.040 | 0.062 |

| Species | N | Formula | $R^2$ | Covariate | Est. | SE | *z* | *p* | *lower 95% CI* | *upper 95% CI* |
|---|---|---|---|---|---|---|---|---|---|---|
| | | | | packm | 0.027 | 0.025 | 1.056 | 0.291 | -0.023 | 0.076 |
| | | | | SixMonthPrecip | 0.026 | 0.021 | 1.220 | 0.222 | -0.016 | 0.068 |
| | | | | SoilpH | 0.060 | 0.022 | 2.750 | **0.006** | 0.017 | 0.102 |
| | | | | Ydays | 0.001 | 0.001 | 1.091 | 0.275 | -0.001 | 0.004 |
| *Ratibida columnifera* | 27 | Seed mass ~ Aspect | 0.195 | (Intercept) | -0.437 | 0.029 | -15.102 | < **0.001** | -0.494 | -0.380 |
| | | | | Aspect | -0.070 | 0.028 | -2.500 | **0.012** | -0.125 | -0.015 |
| *Rosa woodsii* | 44 | Seed mass ~ Year + SixMonthPrecip + DeltSpringMax + SoilpH | 0.521 | (Intercept) | 2.177 | 0.027 | 79.331 | < **0.001** | 2.123 | 2.231 |
| | | | | DeltSpringMax | 0.027 | 0.034 | 0.814 | 0.416 | -0.038 | 0.093 |

| Species | N | Formula | $R^2$ | Covariate | Est. | SE | *z* | *p* | *lower 95% CI* | *upper 95% CI* |
|---|---|---|---|---|---|---|---|---|---|---|
| | | | | SixMonthPrecip | 0.116 | 0.028 | 4.147 | **< 0.001** | 0.061 | 0.171 |
| | | | | SoilpH | 0.063 | 0.028 | 2.287 | **0.022** | 0.009 | 0.117 |
| | | | | Year | -0.021 | 0.011 | -1.890 | 0.059 | -0.043 | 0.001 |
| *Salvia columbariae* | 31 | Seed mass ~ Year + SixMonthPrecip + DeltSpringMax + SoilpH | 0.315 | (Intercept) | -0.018 | 0.030 | -0.609 | 0.543 | -0.077 | 0.041 |
| | | | | DeltSpringMax | 0.050 | 0.028 | 1.781 | 0.075 | -0.005 | 0.106 |
| | | | | SixMonthPrecip | -0.047 | 0.029 | -1.597 | 0.110 | -0.104 | 0.011 |
| | | | | SoilpH | 0.012 | 0.026 | 0.476 | 0.634 | -0.039 | 0.064 |
| | | | | Year | 0.005 | 0.007 | 0.646 | 0.518 | -0.010 | 0.019 |

| Species | N | Formula | $R^2$ | Covariate | Est. | SE | *z* | *p* | *lower 95% CI* | *upper 95% CI* |
|---|---|---|---|---|---|---|---|---|---|---|
| *Sedum lanceolatum* | 26 | Seed mass ~ AWC | 0.103 | (Intercept) | -2.720 | 0.054 | -50.565 | < **0.001** | -2.826 | -2.615 |
| | | | | AWC | 0.051 | 0.061 | 0.837 | 0.402 | -0.069 | 0.171 |
| *Sphaeralcea ambigua* | 55 | Seed mass ~ Ydays | 0.204 | (Intercept) | 0.311 | 0.022 | 14.407 | < **0.001** | 0.269 | 0.353 |
| | | | | Ydays | -0.002 | 0.001 | -3.303 | **0.001** | -0.003 | -0.001 |
| *Sphaeralcea coccinea* | 31 | Seed mass ~ Ydays + DeltSpringMax | 0.405 | (Intercept) | 0.907 | 0.061 | 14.928 | < **0.001** | 0.788 | 1.027 |
| | | | | DeltSpringMax | -0.131 | 0.051 | -2.568 | **0.010** | -0.231 | -0.031 |
| | | | | Ydays | -0.004 | 0.002 | -2.031 | **0.042** | -0.007 | 0.000 |
| *Sphaeralcea parvifolia* | 54 | Seed mass ~ Ydays + SoilpH + organic | 0.312 | (Intercept) | 0.118 | 0.029 | 4.004 | < **0.001** | 0.060 | 0.176 |

| Species | N | Formula | $R^2$ | Covariate | Est. | SE | *z* | *p* | *lower 95% CI* | *upper 95% CI* |
|---|---|---|---|---|---|---|---|---|---|---|
| | | | | organic | -0.027 | 0.026 | -1.052 | 0.293 | -0.078 | 0.024 |
| | | | | SoilpH | 0.073 | 0.030 | 2.427 | **0.015** | 0.014 | 0.131 |
| | | | | Ydays | -0.001 | 0.001 | -0.549 | 0.583 | -0.003 | 0.001 |
| *Sporobolus airoides* | 82 | Seed mass ~ packm + heatload + DeltFallMax | 0.290 | (Intercept) | -1.349 | 0.030 | -45.710 | < **0.001** | -1.407 | -1.291 |
| | | | | DeltFallMax | 0.102 | 0.028 | 3.634 | < **0.001** | 0.047 | 0.157 |
| | | | | heatload | 0.069 | 0.026 | 2.604 | **0.009** | 0.017 | 0.120 |
| | | | | packm | 0.038 | 0.027 | 1.441 | 0.150 | -0.014 | 0.090 |
| *Sporobolus contractus* | 36 | Seed mass ~ heatload + DeltFallMax + organic | 0.346 | (Intercept) | -1.946 | 0.020 | -96.338 | < **0.001** | -1.986 | -1.907 |

| Species | N | Formula | $R^2$ | Covariate | Est. | SE | *z* | *p* | *lower 95% CI* | *upper 95% CI* |
|---|---|---|---|---|---|---|---|---|---|---|
| | | | | DeltFallMax | -0.021 | 0.022 | -0.942 | 0.346 | -0.065 | 0.023 |
| | | | | heatload | 0.047 | 0.022 | 2.151 | **0.032** | 0.004 | 0.090 |
| | | | | organic | -0.079 | 0.024 | -3.327 | **0.001** | -0.125 | -0.032 |
| *Sporobolus cryptandrus* | 78 | Seed mass ~ Year + DeltFallMax | 0.154 | (Intercept) | -2.266 | 0.038 | -60.403 | < **0.001** | -2.339 | -2.192 |
| | | | | DeltFallMax | 0.118 | 0.039 | 2.996 | **0.003** | 0.041 | 0.195 |
| | | | | Year | -0.022 | 0.013 | -1.726 | 0.084 | -0.046 | 0.003 |
| *Sporobolus flexuosus* | 32 | Seed mass ~ heatload | 0.352 | (Intercept) | -2.127 | 0.050 | -42.683 | < **0.001** | -2.225 | -2.029 |
| | | | | heatload | 0.208 | 0.049 | 4.285 | < **0.001** | 0.113 | 0.303 |
| *Stanleya pinnata* | 26 | Seed mass ~ heatload | 0.264 | (Intercept) | 0.099 | 0.093 | 1.063 | 0.288 | -0.083 | 0.280 |
| | | | | heatload | 0.296 | 0.096 | 3.094 | **0.002** | 0.108 | 0.483 |

| Species | N | Formula | $R^2$ | Covariate | Est. | SE | *z* | *p* | *lower 95% CI* | *upper 95% CI* |
|---|---|---|---|---|---|---|---|---|---|---|
| *Stenotus acaulis* | 23 | Seed mass ~ Ydays | 0.088 | (Intercept) | 0.030 | 0.049 | 0.608 | 0.543 | -0.067 | 0.127 |
| | | | | Ydays | 0.006 | 0.004 | 1.506 | 0.132 | -0.002 | 0.014 |
| *Tetradymia canescens* | 24 | Seed mass ~ Ydays + DeltSpringMax + SoilpH | 0.749 | (Intercept) | 1.414 | 0.034 | 41.509 | < **0.001** | 1.347 | 1.481 |
| | | | | DeltSpringMax | 0.099 | 0.046 | 2.166 | **0.030** | 0.009 | 0.189 |
| | | | | SoilpH | -0.138 | 0.047 | -2.916 | **0.004** | -0.231 | -0.045 |
| | | | | Ydays | 0.007 | 0.002 | 4.808 | < **0.001** | 0.004 | 0.010 |
| *Tetradymia glabrata* | 21 | Seed mass ~ clay | 0.274 | (Intercept) | 0.903 | 0.053 | 17.159 | < **0.001** | 0.800 | 1.006 |
| | | | | clay | 0.065 | 0.039 | 1.672 | 0.094 | -0.011 | 0.140 |

| Species | N | Formula | $R^2$ | Covariate | Est. | SE | *z* | *p* | *lower 95% CI* | *upper 95% CI* |
|---|---|---|---|---|---|---|---|---|---|---|
| *Verbesina encelioides* | 23 | Seed mass ~ Year | 0.272 | (Intercept) | 0.643 | 0.035 | 18.316 | < **0.001** | 0.574 | 0.711 |
| | | | | Year | -0.028 | 0.009 | -3.065 | **0.002** | -0.045 | -0.010 |
| *Wyethia amplexicaulis* | 22 | Seed mass ~ Year | 0.281 | (Intercept) | 2.703 | 0.021 | 126.230 | < **0.001** | 2.661 | 2.745 |
| | | | | Year | -0.024 | 0.008 | -2.990 | **0.003** | -0.039 | -0.008 |

**Table S6.** Results of subsampling. For each species in the 20 collection dataset that had been collected from at least 25 populations, we randomly sampled 20 populations 500 times with replacement and a minimum requirement of 15 unique populations. These 20 populations were then moved through the random forest pipeline to determine covariates. These covariates were then used a spatial error GLM and results were saved. Coefficient values for each covariates from each model were generalized and the percentiles are reported below in addition to the original model coefficient values when using the full species' dataset. We also report the counts of each time a covariate was selected in the 500 iterations and how many times that covariate was statistically significant at the p < 0.05 level.

| Species | Variable | Original model coefficient | Original model p | Median | Q 0% | Q 25% | Q 50% | Q 75% | Q 100% | Model Counts | P < 0.05 counts |
|---|---|---|---|---|---|---|---|---|---|---|---|
| *Achillea millefolium* | AnnMinTemp | | | 0.041 | -0.196 | -0.045 | 0.041 | 0.074 | 0.149 | 66 | 29 |
| | Aspect | | | 0.057 | -0.148 | 0.035 | 0.057 | 0.083 | 0.133 | 42 | 24 |
| | AWC | | | -0.028 | -0.135 | -0.073 | -0.028 | 0.036 | 0.131 | 22 | 10 |
| | clay | | | -0.051 | -0.104 | -0.077 | -0.051 | 0.039 | 0.172 | 27 | 16 |
| | DeltFallMax | | | -0.048 | -0.142 | -0.072 | -0.048 | 0.001 | 0.074 | 21 | 11 |
| | DeltFallMin | | | -0.078 | -0.15 | -0.101 | -0.078 | -0.044 | 0.044 | 50 | 29 |
| | DeltPrecip | | | 0.052 | -0.052 | 0.026 | 0.052 | 0.083 | 0.137 | 41 | 15 |

| Species | Variable | Original model coefficient | Original model p | Median | Q 0% | Q 25% | Q 50% | Q 75% | Q 100% | Model Counts | P < 0.05 counts |
|---|---|---|---|---|---|---|---|---|---|---|---|
| | DeltSpringMax | | | 0.071 | 0.009 | 0.053 | 0.071 | 0.101 | 0.203 | 53 | 34 |
| | heatload | | | 0.042 | -0.197 | -0.011 | 0.042 | 0.07 | 0.116 | 35 | 19 |
| | organic | | | 0.027 | -0.094 | 0.003 | 0.027 | 0.075 | 0.328 | 65 | 14 |
| | packm | | | -0.056 | -0.216 | -0.118 | -0.056 | -0.004 | 0.013 | 12 | 3 |
| | SixMonthPrecip | | | 0.059 | -0.085 | 0.023 | 0.059 | 0.082 | 0.167 | 73 | 29 |
| | Slope | | | 0.076 | -0.049 | 0.047 | 0.076 | 0.101 | 0.23 | 71 | 49 |
| | SoilpH | -0.072 | <0.001 | -0.083 | -0.189 | -0.108 | -0.083 | -0.055 | 0.064 | 184 | 121 |
| | Ydays | | | 0.001 | -0.004 | 0 | 0.001 | 0.002 | 0.007 | 36 | 16 |
| | Year | | | -0.022 | -0.054 | -0.03 | -0.022 | -0.007 | 0.02 | 60 | 37 |
| *Achnatherum hymenoides* | AnnMinTemp | 0.149 | <0.001 | 0.163 | -0.155 | 0.109 | 0.163 | 0.228 | 0.418 | 202 | 133 |
| | Aspect | | | -0.012 | -0.235 | -0.119 | -0.012 | 0.152 | 0.258 | 20 | 14 |
| | AWC | 0.055 | 0.063 | 0.118 | -0.185 | 0.086 | 0.118 | 0.164 | 0.346 | 36 | 18 |
| | clay | -0.03 | 0.315 | -0.048 | -0.25 | -0.185 | -0.048 | 0.144 | 0.341 | 17 | 10 |
| | DeltFallMax | -0.01 | 0.668 | -0.141 | -0.357 | -0.202 | -0.141 | -0.076 | 0.121 | 77 | 46 |

| Species | Variable | Original model coefficient | Original model p | Median | Q 0% | Q 25% | Q 50% | Q 75% | Q 100% | Model Counts | P < 0.05 counts |
|---|---|---|---|---|---|---|---|---|---|---|---|
| | DeltFallMin | | | -0.056 | -0.366 | -0.154 | -0.056 | 0.104 | 0.212 | 24 | 9 |
| | DeltPrecip | | | 0.068 | -0.174 | 0.009 | 0.068 | 0.138 | 0.277 | 36 | 13 |
| | DeltSpringMax | -0.007 | 0.772 | 0.048 | -0.338 | -0.079 | 0.048 | 0.115 | 0.248 | 47 | 17 |
| | heatload | | | 0.074 | -0.143 | 0.011 | 0.074 | 0.131 | 0.317 | 45 | 8 |
| | organic | 0.003 | 0.908 | -0.086 | -0.318 | -0.17 | -0.086 | -0.062 | 0.163 | 25 | 13 |
| | packm | | | -0.044 | -0.146 | -0.105 | -0.044 | -0.001 | 0.122 | 22 | 5 |
| | SixMonthPrecip | | | -0.11 | -0.351 | -0.152 | -0.11 | -0.064 | 0.175 | 35 | 12 |
| | Slope | | | -0.025 | -0.23 | -0.096 | -0.025 | -0.007 | 0.228 | 17 | 2 |
| | SoilpH | 0.06 | 0.029 | 0.131 | -0.293 | 0.077 | 0.131 | 0.173 | 0.265 | 74 | 43 |
| | Ydays | 0 | 0.653 | -0.005 | -0.012 | -0.007 | -0.005 | -0.002 | 0.001 | 94 | 44 |
| | Year | | | -0.041 | -0.086 | -0.055 | -0.041 | -0.009 | 0.041 | 24 | 18 |
| *Achnatherum speciosum* | AnnMinTemp | -0.017 | 0.616 | -0.045 | -0.184 | -0.067 | -0.045 | 0.005 | 0.088 | 33 | 11 |
| | Aspect | | | 0.099 | -0.02 | 0.077 | 0.099 | 0.117 | 0.211 | 206 | 177 |
| | AWC | 0.107 | 0.019 | 0.052 | -0.055 | 0.026 | 0.052 | 0.078 | 0.142 | 56 | 32 |

| Species | Variable | Original model coefficient | Original model p | Median | Q 0% | Q 25% | Q 50% | Q 75% | Q 100% | Model Counts | P < 0.05 counts |
|---|---|---|---|---|---|---|---|---|---|---|---|
| | clay | -0.083 | 0.040 | -0.045 | -0.108 | -0.071 | -0.045 | -0.026 | 0.069 | 65 | 26 |
| | DeltFallMax | -0.076 | 0.078 | -0.036 | -0.072 | -0.062 | -0.036 | 0.082 | 0.172 | 13 | 5 |
| | DeltFallMin | | | -0.059 | -0.167 | -0.078 | -0.059 | -0.033 | 0.059 | 42 | 18 |
| | DeltPrecip | | | -0.083 | -0.132 | -0.086 | -0.083 | -0.07 | -0.066 | 5 | 5 |
| | DeltSpringMax | 0.05 | 0.211 | 0.069 | -0.061 | 0.044 | 0.069 | 0.084 | 0.14 | 65 | 34 |
| | heatload | | | 0.021 | -0.092 | 0.006 | 0.021 | 0.053 | 0.2 | 32 | 7 |
| | organic | -0.058 | 0.116 | -0.046 | -0.111 | -0.062 | -0.046 | -0.026 | 0.023 | 19 | 7 |
| | packm | | | 0.01 | -0.029 | -0.014 | 0.01 | 0.063 | 0.077 | 6 | 1 |
| | SixMonthPrecip | | | -0.052 | -0.297 | -0.078 | -0.052 | -0.023 | 0.089 | 106 | 49 |
| | Slope | | | 0.013 | -0.08 | -0.029 | 0.013 | 0.042 | 0.208 | 39 | 8 |
| | SoilpH | -0.04 | 0.271 | -0.061 | -0.146 | -0.083 | -0.061 | -0.031 | 0.061 | 61 | 27 |
| | Ydays | -0.001 | 0.467 | -0.003 | -0.01 | -0.004 | -0.003 | -0.002 | 0.006 | 194 | 86 |
| | Year | | | -0.01 | -0.018 | -0.014 | -0.01 | 0.002 | 0.024 | 15 | 4 |
| *Achnatherum thurberianum* | AnnMinTemp | -0.039 | 0.302 | 0.018 | -0.184 | -0.036 | 0.018 | 0.05 | 0.153 | 95 | 20 |

| Species | Variable | Original model coefficient | Original model p | Median | Q 0% | Q 25% | Q 50% | Q 75% | Q 100% | Model Counts | P < 0.05 counts |
|---|---|---|---|---|---|---|---|---|---|---|---|
| | Aspect | | | 0.086 | -0.052 | 0.058 | 0.086 | 0.118 | 0.22 | 43 | 22 |
| | AWC | | | -0.013 | -0.304 | -0.045 | -0.013 | 0.055 | 0.159 | 39 | 11 |
| | clay | 0.092 | 0.006 | 0.116 | -0.119 | 0.086 | 0.116 | 0.141 | 0.245 | 175 | 134 |
| | DeltFallMax | -0.048 | 0.137 | -0.072 | -0.318 | -0.108 | -0.072 | -0.03 | 0.164 | 214 | 91 |
| | DeltFallMin | | | -0.065 | -0.201 | -0.12 | -0.065 | 0.001 | 0.064 | 18 | 11 |
| | DeltPrecip | | | 0.024 | -0.095 | -0.013 | 0.024 | 0.057 | 0.138 | 16 | 5 |
| | DeltSpringMax | | | 0.022 | -0.08 | -0.015 | 0.022 | 0.073 | 0.178 | 21 | 3 |
| | heatload | | | 0.088 | -0.11 | 0.03 | 0.088 | 0.144 | 0.289 | 63 | 32 |
| | organic | -0.025 | 0.486 | -0.021 | -0.234 | -0.07 | -0.021 | 0.042 | 0.283 | 151 | 56 |
| | packm | | | -0.081 | -0.193 | -0.124 | -0.081 | -0.05 | 0.03 | 21 | 12 |
| | SixMonthPrecip | -0.11 | 0.001 | -0.128 | -0.381 | -0.169 | -0.128 | -0.09 | 0.137 | 287 | 221 |
| | Slope | | | -0.049 | -0.234 | -0.083 | -0.049 | 0.017 | 0.115 | 18 | 6 |
| | SoilpH | -0.016 | 0.621 | 0.061 | -0.193 | 0.021 | 0.061 | 0.099 | 0.177 | 157 | 59 |
| | Ydays | -0.007 | 0.006 | -0.007 | -0.018 | -0.009 | -0.007 | -0.004 | 0.005 | 191 | 126 |
| | Year | | | 0.003 | -0.032 | -0.002 | 0.003 | 0.014 | 0.054 | 26 | 3 |

| Species | Variable | Original model coefficient | Original model p | Median | Q 0% | Q 25% | Q 50% | Q 75% | Q 100% | Model Counts | P < 0.05 counts |
|---|---|---|---|---|---|---|---|---|---|---|---|
| *Agastache urticifolia* | AnnMinTemp | | | 0.046 | -0.065 | 0.017 | 0.046 | 0.061 | 0.115 | 63 | 25 |
| | Aspect | | | 0.054 | -0.062 | 0.033 | 0.054 | 0.071 | 0.118 | 68 | 41 |
| | AWC | | | 0.045 | -0.046 | 0.029 | 0.045 | 0.068 | 0.13 | 48 | 18 |
| | clay | | | -0.029 | -0.075 | -0.047 | -0.029 | -0.016 | 0.107 | 26 | 8 |
| | DeltFallMax | | | -0.012 | -0.104 | -0.027 | -0.012 | -0.001 | 0.059 | 21 | 2 |
| | DeltFallMin | | | 0.014 | -0.085 | -0.029 | 0.014 | 0.056 | 0.075 | 28 | 11 |
| | DeltPrecip | -0.036 | 0.121 | -0.055 | -0.14 | -0.077 | -0.055 | -0.029 | 0.128 | 184 | 92 |
| | DeltSpringMax | | | -0.064 | -0.115 | -0.082 | -0.064 | 0.02 | 0.106 | 45 | 29 |
| | heatload | | | -0.05 | -0.167 | -0.069 | -0.05 | -0.038 | 0.029 | 76 | 41 |
| | organic | | | 0.037 | -0.127 | -0.001 | 0.037 | 0.045 | 0.159 | 36 | 12 |
| | packm | | | 0.056 | -0.019 | 0.032 | 0.056 | 0.076 | 0.13 | 29 | 17 |
| | SixMonthPrecip | | | 0.052 | -0.028 | 0.041 | 0.052 | 0.069 | 0.138 | 20 | 14 |
| | Slope | | | 0.043 | -0.041 | 0.031 | 0.043 | 0.06 | 0.102 | 61 | 30 |
| | SoilpH | | | -0.049 | -0.142 | -0.068 | -0.049 | -0.034 | 0.03 | 97 | 48 |

| Species | Variable | Original model coefficient | Original model p | Median | Q 0% | Q 25% | Q 50% | Q 75% | Q 100% | Model Counts | P < 0.05 counts |
|---|---|---|---|---|---|---|---|---|---|---|---|
| | Ydays | | | 0.002 | -0.006 | 0 | 0.002 | 0.003 | 0.004 | 17 | 12 |
| | Year | | | -0.015 | -0.051 | -0.02 | -0.015 | -0.011 | 0.017 | 71 | 48 |
| *Allium textile* | AnnMinTemp | | | -0.016 | -0.151 | -0.036 | -0.016 | -0.002 | 0.118 | 77 | 14 |
| | Aspect | | | 0.056 | -0.057 | 0.04 | 0.056 | 0.073 | 0.117 | 77 | 53 |
| | AWC | | | -0.025 | -0.067 | -0.046 | -0.025 | -0.011 | 0.037 | 12 | 3 |
| | clay | | | -0.032 | -0.079 | -0.046 | -0.032 | -0.016 | 0.083 | 24 | 5 |
| | DeltFallMax | | | -0.022 | -0.052 | -0.03 | -0.022 | -0.001 | 0.001 | 5 | 0 |
| | DeltFallMin | | | -0.032 | -0.045 | -0.035 | -0.032 | 0.025 | 0.102 | 7 | 1 |
| | DeltPrecip | | | -0.054 | -0.148 | -0.072 | -0.054 | -0.042 | 0.086 | 58 | 40 |
| | DeltSpringMax | | | -0.03 | -0.099 | -0.072 | -0.03 | 0.004 | 0.062 | 23 | 9 |
| | heatload | | | -0.028 | -0.15 | -0.075 | -0.028 | 0.02 | 0.201 | 49 | 22 |
| | organic | -0.102 | <0.001 | -0.112 | -0.345 | -0.143 | -0.112 | -0.088 | -0.039 | 390 | 371 |
| | packm | | | -0.026 | -0.175 | -0.085 | -0.026 | -0.006 | 0.052 | 21 | 9 |
| | SixMonthPrecip | | | 0.051 | -0.021 | 0.025 | 0.051 | 0.084 | 0.277 | 84 | 35 |
| | Slope | -0.031 | 0.200 | -0.049 | -0.236 | -0.075 | -0.049 | -0.026 | 0.068 | 256 | 110 |

| Species | Variable | Original model coefficient | Original model p | Median | Q 0% | Q 25% | Q 50% | Q 75% | Q 100% | Model Counts | P < 0.05 counts |
|---|---|---|---|---|---|---|---|---|---|---|---|
| | SoilpH | | | -0.058 | -0.248 | -0.093 | -0.058 | -0.016 | 0.041 | 123 | 47 |
| | Ydays | | | -0.001 | -0.012 | -0.002 | -0.001 | 0 | 0.005 | 8 | 2 |
| | Year | | | 0.022 | -0.003 | 0.015 | 0.022 | 0.036 | 0.059 | 15 | 10 |
| *Ambrosia dumosa* | AnnMinTemp | 0.107 | <0.001 | 0.104 | -0.166 | 0.063 | 0.104 | 0.143 | 0.26 | 262 | 164 |
| | Aspect | | | -0.078 | -0.217 | -0.101 | -0.078 | -0.052 | 0.119 | 59 | 31 |
| | AWC | | | -0.07 | -0.247 | -0.116 | -0.07 | -0.047 | 0.036 | 24 | 10 |
| | clay | | | 0.035 | -0.171 | -0.041 | 0.035 | 0.07 | 0.274 | 62 | 15 |
| | DeltFallMax | -0.042 | 0.140 | 0.038 | -0.219 | -0.042 | 0.038 | 0.074 | 0.25 | 49 | 13 |
| | DeltFallMin | | | -0.086 | -0.171 | -0.116 | -0.086 | -0.027 | 0.031 | 26 | 15 |
| | DeltPrecip | 0.045 | 0.172 | 0.067 | -0.145 | 0.028 | 0.067 | 0.119 | 0.305 | 117 | 44 |
| | DeltSpringMax | | | -0.078 | -0.209 | -0.114 | -0.078 | -0.029 | 0.051 | 54 | 23 |
| | heatload | | | 0.027 | -0.152 | -0.012 | 0.027 | 0.097 | 0.382 | 89 | 24 |
| | organic | | | 0.026 | -0.47 | -0.067 | 0.026 | 0.058 | 0.108 | 27 | 8 |
| | packm | | | 0.004 | -1.732 | -0.035 | 0.004 | 0.063 | 1.047 | 63 | 9 |
| | SixMonthPrecip | 0.088 | 0.002 | 0.117 | -0.077 | 0.078 | 0.117 | 0.177 | 0.356 | 142 | 96 |

| Species | Variable | Original model coefficient | Original model p | Median | Q 0% | Q 25% | Q 50% | Q 75% | Q 100% | Model Counts | P < 0.05 counts |
|---|---|---|---|---|---|---|---|---|---|---|---|
| | Slope | | | -0.064 | -0.412 | -0.117 | -0.064 | -0.025 | 0.205 | 71 | 29 |
| | SoilpH | | | 0.057 | -0.151 | -0.021 | 0.057 | 0.107 | 0.346 | 32 | 14 |
| | Ydays | 0.001 | 0.040 | 0.001 | -0.004 | 0 | 0.001 | 0.002 | 0.006 | 55 | 26 |
| | Year | -0.025 | 0.005 | -0.03 | -0.094 | -0.041 | -0.03 | -0.018 | 0.046 | 298 | 190 |
| *Argemone munita* | AnnMinTemp | | | -0.037 | -0.124 | -0.049 | -0.037 | -0.027 | 0.052 | 43 | 27 |
| | Aspect | | | -0.01 | -0.08 | -0.027 | -0.01 | 0.013 | 0.159 | 137 | 41 |
| | AWC | | | 0.027 | -0.061 | 0.002 | 0.027 | 0.033 | 0.048 | 8 | 5 |
| | clay | | | 0.015 | -0.024 | 0.009 | 0.015 | 0.027 | 0.059 | 40 | 11 |
| | DeltFallMax | | | 0.022 | -0.052 | 0.002 | 0.022 | 0.039 | 0.085 | 87 | 23 |
| | DeltFallMin | | | 0.023 | -0.061 | 0.008 | 0.023 | 0.031 | 0.065 | 69 | 26 |
| | DeltPrecip | | | -0.001 | -0.046 | -0.015 | -0.001 | 0.011 | 0.028 | 30 | 0 |
| | DeltSpringMax | | | 0.014 | -0.021 | -0.003 | 0.014 | 0.029 | 0.068 | 25 | 4 |
| | heatload | | | -0.031 | -0.112 | -0.044 | -0.031 | -0.019 | 0.066 | 105 | 39 |
| | organic | | | 0.031 | -0.132 | 0.013 | 0.031 | 0.071 | 0.168 | 60 | 31 |
| | packm | | | -0.021 | -0.152 | -0.033 | -0.021 | -0.009 | 0.036 | 200 | 56 |

| Species | Variable | Original model coefficient | Original model p | Median | Q 0% | Q 25% | Q 50% | Q 75% | Q 100% | Model Counts | P < 0.05 counts |
|---|---|---|---|---|---|---|---|---|---|---|---|
| | SixMonthPrecip | | | 0.033 | -0.048 | 0.009 | 0.033 | 0.051 | 0.146 | 214 | 113 |
| | Slope | | | 0.011 | -0.037 | -0.002 | 0.011 | 0.025 | 0.09 | 90 | 17 |
| | SoilpH | -0.06 | 0.001 | -0.048 | -0.142 | -0.061 | -0.048 | -0.034 | 0.036 | 387 | 275 |
| | Ydays | | | 0.002 | -0.001 | 0.001 | 0.002 | 0.002 | 0.004 | 23 | 16 |
| | Year | | | -0.016 | -0.016 | -0.016 | -0.016 | -0.016 | -0.016 | 1 | 1 |
| *Aristida purpurea* | AnnMinTemp | | | -0.207 | -0.313 | -0.238 | -0.207 | -0.136 | 0.285 | 40 | 31 |
| | Aspect | | | -0.148 | -0.335 | -0.208 | -0.148 | -0.101 | 0.128 | 30 | 21 |
| | AWC | | | 0.131 | -0.178 | 0.026 | 0.131 | 0.174 | 0.395 | 33 | 16 |
| | clay | | | -0.071 | -0.238 | -0.109 | -0.071 | 0.057 | 0.301 | 20 | 10 |
| | DeltFallMax | | | 0.046 | -0.385 | -0.049 | 0.046 | 0.12 | 0.223 | 35 | 8 |
| | DeltFallMin | | | 0.173 | -0.332 | 0.124 | 0.173 | 0.234 | 0.435 | 96 | 75 |
| | DeltPrecip | | | 0.048 | -0.256 | -0.102 | 0.048 | 0.198 | 0.528 | 22 | 10 |
| | DeltSpringMax | | | -0.152 | -0.259 | -0.19 | -0.152 | -0.111 | 0.124 | 50 | 37 |
| | heatload | | | -0.016 | -0.328 | -0.107 | -0.016 | 0.038 | 0.318 | 49 | 14 |
| | organic | | | 0.142 | -0.138 | 0.089 | 0.142 | 0.22 | 0.375 | 62 | 30 |

| Species | Variable | Original model coefficient | Original model p | Median | Q 0% | Q 25% | Q 50% | Q 75% | Q 100% | Model Counts | P < 0.05 counts |
|---|---|---|---|---|---|---|---|---|---|---|---|
| | packm | | | -0.202 | -0.528 | -0.283 | -0.202 | -0.064 | 0.052 | 43 | 23 |
| | SixMonthPrecip | | | 0.183 | -0.17 | 0.066 | 0.183 | 0.268 | 0.583 | 17 | 11 |
| | Slope | | | -0.119 | -0.359 | -0.149 | -0.119 | -0.09 | 0.257 | 42 | 25 |
| | SoilpH | | | 0.023 | -0.143 | -0.044 | 0.023 | 0.097 | 0.188 | 18 | 3 |
| | Ydays | -0.001 | 0.247 | -0.004 | -0.009 | -0.005 | -0.004 | -0.001 | 0.003 | 182 | 109 |
| | Year | 0.028 | 0.222 | 0.044 | -0.094 | 0.027 | 0.044 | 0.06 | 0.179 | 152 | 47 |
| *Artemisia tridentata* | AnnMinTemp | 0.035 | 0.080 | 0.099 | -0.098 | 0.033 | 0.099 | 0.137 | 0.25 | 42 | 24 |
| | Aspect | | | 0.095 | -0.157 | 0.014 | 0.095 | 0.135 | 0.188 | 36 | 22 |
| | AWC | -0.089 | <0.001 | -0.117 | -0.246 | -0.146 | -0.117 | -0.08 | 0.123 | 34 | 24 |
| | clay | 0.058 | 0.010 | 0.104 | -0.102 | 0.056 | 0.104 | 0.138 | 0.37 | 62 | 36 |
| | DeltFallMax | 0.018 | 0.379 | -0.092 | -0.263 | -0.145 | -0.092 | -0.004 | 0.122 | 37 | 19 |
| | DeltFallMin | | | 0.017 | -0.145 | -0.078 | 0.017 | 0.088 | 0.249 | 20 | 8 |
| | DeltPrecip | -0.089 | 0.001 | -0.099 | -0.258 | -0.132 | -0.099 | -0.071 | 0.105 | 41 | 25 |
| | DeltSpringMax | -0.011 | 0.624 | 0.055 | -0.143 | -0.027 | 0.055 | 0.109 | 0.199 | 37 | 13 |
| | heatload | | | -0.114 | -0.248 | -0.168 | -0.114 | -0.062 | 0.239 | 41 | 20 |

| Species | Variable | Original model coefficient | Original model p | Median | Q 0% | Q 25% | Q 50% | Q 75% | Q 100% | Model Counts | P < 0.05 counts |
|---|---|---|---|---|---|---|---|---|---|---|---|
| | organic | | | -0.097 | -0.51 | -0.187 | -0.097 | 0.001 | 0.496 | 31 | 17 |
| | packm | | | -0.04 | -0.25 | -0.088 | -0.04 | 0.077 | 0.68 | 19 | 5 |
| | SixMonthPrecip | 0.043 | 0.039 | 0.12 | -0.088 | 0.075 | 0.12 | 0.151 | 0.262 | 99 | 64 |
| | Slope | | | -0.095 | -0.437 | -0.159 | -0.095 | -0.017 | 0.17 | 33 | 16 |
| | SoilpH | | | -0.001 | -0.231 | -0.135 | -0.001 | 0.073 | 0.117 | 20 | 11 |
| | Ydays | | | 0 | -0.009 | -0.003 | 0 | 0.002 | 0.009 | 37 | 12 |
| | Year | -0.041 | <0.001 | -0.044 | -0.126 | -0.055 | -0.044 | -0.031 | -0.002 | 95 | 72 |
| *Atriplex canescens* | AnnMinTemp | | | -0.112 | -0.375 | -0.152 | -0.112 | -0.088 | 0.183 | 46 | 18 |
| | Aspect | | | -0.132 | -0.444 | -0.177 | -0.132 | -0.011 | 0.238 | 28 | 16 |
| | AWC | | | -0.099 | -0.282 | -0.157 | -0.099 | 0.133 | 0.238 | 26 | 13 |
| | clay | | | -0.104 | -0.301 | -0.16 | -0.104 | 0.002 | 0.286 | 35 | 18 |
| | DeltFallMax | | | -0.059 | -0.247 | -0.161 | -0.059 | 0.018 | 0.363 | 37 | 12 |
| | DeltFallMin | | | -0.168 | -0.507 | -0.21 | -0.168 | -0.077 | 0.135 | 39 | 22 |
| | DeltPrecip | | | -0.049 | -0.255 | -0.147 | -0.049 | 0.164 | 0.342 | 30 | 19 |
| | DeltSpringMax | | | -0.103 | -0.224 | -0.161 | -0.103 | -0.059 | 0.391 | 28 | 11 |

| Species | Variable | Original model coefficient | Original model p | Median | Q 0% | Q 25% | Q 50% | Q 75% | Q 100% | Model Counts | P < 0.05 counts |
|---|---|---|---|---|---|---|---|---|---|---|---|
| | heatload | -0.077 | 0.039 | -0.116 | -0.389 | -0.166 | -0.116 | -0.056 | 0.133 | 76 | 28 |
| | organic | 0.024 | 0.513 | -0.012 | -0.292 | -0.072 | -0.012 | 0.052 | 0.239 | 64 | 11 |
| | packm | | | 0.093 | -0.424 | -0.171 | 0.093 | 0.247 | 0.379 | 25 | 15 |
| | SixMonthPrecip | | | -0.172 | -0.387 | -0.222 | -0.172 | -0.123 | -0.016 | 69 | 42 |
| | Slope | | | 0.052 | -0.367 | 0.021 | 0.052 | 0.117 | 0.663 | 28 | 3 |
| | SoilpH | | | -0.149 | -0.408 | -0.206 | -0.149 | 0.011 | 0.146 | 31 | 15 |
| | Ydays | | | -0.001 | -0.012 | -0.004 | -0.001 | 0 | 0.004 | 23 | 11 |
| | Year | | | -0.023 | -0.144 | -0.054 | -0.023 | 0.02 | 0.066 | 32 | 11 |
| *Atriplex confertifolia* | AnnMinTemp | | | 0.159 | -0.109 | 0.068 | 0.159 | 0.219 | 0.389 | 42 | 15 |
| | Aspect | -0.059 | 0.178 | -0.162 | -0.291 | -0.214 | -0.162 | -0.11 | 0.075 | 76 | 51 |
| | AWC | | | 0.06 | -0.565 | -0.04 | 0.06 | 0.137 | 0.361 | 94 | 33 |
| | clay | | | 0.071 | -0.11 | 0.024 | 0.071 | 0.178 | 0.369 | 45 | 12 |
| | DeltFallMax | | | -0.158 | -0.377 | -0.211 | -0.158 | -0.042 | 0.104 | 45 | 25 |
| | DeltFallMin | | | 0.131 | -0.206 | 0.061 | 0.131 | 0.198 | 0.392 | 127 | 56 |

| Species | Variable | Original model coefficient | Original model p | Median | Q 0% | Q 25% | Q 50% | Q 75% | Q 100% | Model Counts | P < 0.05 counts |
|---|---|---|---|---|---|---|---|---|---|---|---|
| | DeltPrecip | | | -0.139 | -0.363 | -0.194 | -0.139 | -0.05 | 0.179 | 50 | 21 |
| | DeltSpringMax | | | -0.02 | -0.486 | -0.126 | -0.02 | 0.068 | 0.252 | 62 | 18 |
| | heatload | | | 0.067 | -0.337 | -0.018 | 0.067 | 0.137 | 0.526 | 40 | 9 |
| | organic | | | 0.016 | -0.609 | -0.098 | 0.016 | 0.067 | 0.195 | 56 | 13 |
| | packm | | | -0.162 | -0.59 | -0.318 | -0.162 | -0.102 | 0.121 | 71 | 38 |
| | SixMonthPrecip | 0.097 | 0.050 | 0.086 | -0.303 | 0.017 | 0.086 | 0.175 | 0.512 | 276 | 81 |
| | Slope | | | 0.069 | -0.171 | 0.015 | 0.069 | 0.145 | 0.785 | 36 | 12 |
| | SoilpH | -0.129 | 0.008 | -0.166 | -0.455 | -0.212 | -0.166 | -0.095 | 0.185 | 135 | 79 |
| | Ydays | -0.001 | 0.130 | -0.004 | -0.014 | -0.006 | -0.004 | -0.001 | 0.003 | 143 | 52 |
| | Year | | | -0.013 | -0.126 | -0.039 | -0.013 | 0.007 | 0.18 | 66 | 17 |
| *Atriplex gardneri* | AnnMinTemp | 0.346 | <0.001 | 0.289 | -0.039 | 0.25 | 0.289 | 0.339 | 0.645 | 488 | 454 |
| | Aspect | | | 0.106 | -0.04 | 0.068 | 0.106 | 0.143 | 0.283 | 9 | 4 |
| | AWC | | | -0.002 | -0.201 | -0.064 | -0.002 | 0.095 | 0.167 | 11 | 3 |
| | clay | | | 0.066 | -0.106 | 0.022 | 0.066 | 0.1 | 0.283 | 70 | 18 |
| | DeltFallMax | | | -0.14 | -0.291 | -0.186 | -0.14 | -0.084 | 0.043 | 84 | 48 |

| Species | Variable | Original model coefficient | Original model p | Median | Q 0% | Q 25% | Q 50% | Q 75% | Q 100% | Model Counts | P < 0.05 counts |
|---|---|---|---|---|---|---|---|---|---|---|---|
| | DeltFallMin | | | -0.143 | -0.319 | -0.198 | -0.143 | -0.094 | 0.028 | 115 | 68 |
| | DeltPrecip | | | 0.041 | -0.045 | 0 | 0.041 | 0.075 | 0.1 | 4 | 1 |
| | DeltSpringMax | | | -0.048 | -0.292 | -0.105 | -0.048 | 0.034 | 0.156 | 54 | 14 |
| | heatload | | | -0.037 | -0.262 | -0.059 | -0.037 | 0.025 | 0.213 | 73 | 5 |
| | organic | | | 0.13 | -0.139 | 0.071 | 0.13 | 0.171 | 0.294 | 181 | 97 |
| | packm | | | 0.101 | -0.165 | 0.032 | 0.101 | 0.176 | 0.37 | 57 | 23 |
| | SixMonthPrecip | | | 0.162 | 0.011 | 0.102 | 0.162 | 0.231 | 0.556 | 90 | 64 |
| | Slope | | | 0.05 | -0.226 | -0.022 | 0.05 | 0.109 | 0.304 | 95 | 21 |
| | SoilpH | | | -0.146 | -0.369 | -0.219 | -0.146 | -0.076 | 0.265 | 46 | 29 |
| | Ydays | | | 0.001 | -0.009 | 0 | 0.001 | 0.004 | 0.011 | 77 | 22 |
| | Year | | | -0.02 | -0.081 | -0.033 | -0.02 | -0.002 | 0.114 | 110 | 29 |
| *Atriplex polycarpa* | AnnMinTemp | | | 0.065 | -0.052 | 0.043 | 0.065 | 0.091 | 0.131 | 50 | 25 |
| | Aspect | | | -0.075 | -0.164 | -0.097 | -0.075 | -0.043 | 0.068 | 30 | 18 |
| | AWC | | | 0.09 | -0.092 | 0.018 | 0.09 | 0.15 | 0.392 | 58 | 21 |
| | clay | | | 0.044 | -0.109 | -0.026 | 0.044 | 0.055 | 0.123 | 15 | 3 |

| Species | Variable | Original model coefficient | Original model p | Median | Q 0% | Q 25% | Q 50% | Q 75% | Q 100% | Model Counts | P < 0.05 counts |
|---|---|---|---|---|---|---|---|---|---|---|---|
| | DeltFallMax | | | 0.054 | -0.133 | 0.037 | 0.054 | 0.074 | 0.162 | 62 | 28 |
| | DeltFallMin | -0.041 | 0.103 | -0.053 | -0.198 | -0.081 | -0.053 | -0.028 | 0.064 | 165 | 74 |
| | DeltPrecip | -0.056 | 0.030 | -0.08 | -0.291 | -0.122 | -0.08 | -0.053 | 0.079 | 399 | 274 |
| | DeltSpringMax | -0.042 | 0.100 | -0.066 | -0.196 | -0.103 | -0.066 | -0.041 | 0.11 | 156 | 82 |
| | heatload | | | 0 | -0.109 | -0.03 | 0 | 0.035 | 0.119 | 21 | 5 |
| | organic | | | 0.019 | -0.168 | -0.009 | 0.019 | 0.037 | 0.094 | 38 | 8 |
| | packm | | | -0.076 | -0.38 | -0.103 | -0.076 | -0.053 | -0.009 | 46 | 34 |
| | SixMonthPrecip | -0.009 | 0.737 | 0.013 | -0.215 | -0.029 | 0.013 | 0.053 | 0.219 | 137 | 34 |
| | Slope | | | -0.028 | -0.285 | -0.083 | -0.028 | 0 | 0.093 | 98 | 30 |
| | SoilpH | | | 0.07 | -0.008 | 0.015 | 0.07 | 0.095 | 0.225 | 19 | 8 |
| | Ydays | | | 0.001 | -0.002 | 0.001 | 0.001 | 0.001 | 0.004 | 42 | 26 |
| | Year | | | 0.01 | -0.021 | 0.001 | 0.01 | 0.025 | 0.088 | 56 | 13 |
| *Atriplex torreyi* | AnnMinTemp | | | -0.097 | -0.271 | -0.145 | -0.097 | 0.029 | 0.249 | 63 | 34 |
| | Aspect | | | 0.015 | -0.194 | -0.035 | 0.015 | 0.054 | 0.293 | 33 | 7 |
| | AWC | | | 0.052 | -0.188 | -0.023 | 0.052 | 0.101 | 0.192 | 74 | 8 |

| Species | Variable | Original model coefficient | Original model p | Median | Q 0% | Q 25% | Q 50% | Q 75% | Q 100% | Model Counts | P < 0.05 counts |
|---|---|---|---|---|---|---|---|---|---|---|---|
| | clay | 0.032 | 0.546 | 0.039 | -0.326 | -0.004 | 0.039 | 0.103 | 0.339 | 262 | 59 |
| | DeltFallMax | | | -0.132 | -0.3 | -0.162 | -0.132 | -0.101 | 0.044 | 25 | 19 |
| | DeltFallMin | | | -0.049 | -0.26 | -0.107 | -0.049 | -0.021 | 0.073 | 21 | 1 |
| | DeltPrecip | | | 0.067 | -0.196 | -0.038 | 0.067 | 0.147 | 0.254 | 52 | 31 |
| | DeltSpringMax | | | -0.094 | -0.144 | -0.119 | -0.094 | -0.07 | -0.045 | 3 | 0 |
| | heatload | | | 0.046 | -0.229 | 0.016 | 0.046 | 0.061 | 0.092 | 14 | 2 |
| | organic | | | 0.13 | -0.183 | 0.071 | 0.13 | 0.204 | 0.42 | 39 | 27 |
| | packm | -0.184 | <0.001 | -0.173 | -0.362 | -0.199 | -0.173 | -0.14 | 0.067 | 395 | 358 |
| | SixMonthPrecip | | | -0.08 | -0.246 | -0.114 | -0.08 | -0.05 | 0.038 | 69 | 25 |
| | Slope | | | 0.013 | -0.231 | -0.013 | 0.013 | 0.065 | 0.294 | 11 | 4 |
| | SoilpH | | | 0.089 | -0.114 | 0.013 | 0.089 | 0.132 | 0.303 | 27 | 10 |
| | Ydays | | | 0.002 | 0.001 | 0.002 | 0.002 | 0.003 | 0.004 | 8 | 0 |
| | Year | | | 0.027 | -0.04 | 0.014 | 0.027 | 0.034 | 0.083 | 57 | 24 |
| *Balsamorhiza sagittata* | AnnMinTemp | | | 0.108 | -0.099 | 0.058 | 0.108 | 0.145 | 0.3 | 87 | 39 |

| Species | Variable | Original model coefficient | Original model p | Median | Q 0% | Q 25% | Q 50% | Q 75% | Q 100% | Model Counts | P < 0.05 counts |
|---|---|---|---|---|---|---|---|---|---|---|---|
| | Aspect | | | 0.084 | -0.16 | 0.034 | 0.084 | 0.125 | 0.199 | 27 | 15 |
| | AWC | | | -0.048 | -0.26 | -0.086 | -0.048 | 0.008 | 0.184 | 58 | 24 |
| | clay | | | 0.039 | -0.183 | -0.052 | 0.039 | 0.108 | 0.321 | 76 | 33 |
| | DeltFallMax | | | 0.031 | -0.121 | -0.061 | 0.031 | 0.085 | 0.214 | 17 | 7 |
| | DeltFallMin | | | -0.084 | -0.28 | -0.139 | -0.084 | -0.046 | 0.103 | 22 | 7 |
| | DeltPrecip | 0.051 | 0.095 | 0.083 | -0.068 | 0.049 | 0.083 | 0.118 | 0.324 | 119 | 50 |
| | DeltSpringMax | | | 0.073 | -0.242 | 0.044 | 0.073 | 0.091 | 0.153 | 52 | 25 |
| | heatload | | | 0.012 | -0.294 | -0.045 | 0.012 | 0.042 | 0.112 | 24 | 5 |
| | organic | | | -0.065 | -0.432 | -0.125 | -0.065 | -0.026 | 0.066 | 35 | 12 |
| | packm | | | 0.07 | -0.255 | -0.035 | 0.07 | 0.101 | 0.15 | 31 | 16 |
| | SixMonthPrecip | | | 0.054 | -0.104 | 0.034 | 0.054 | 0.091 | 0.318 | 18 | 8 |
| | Slope | | | 0.092 | -0.255 | 0.058 | 0.092 | 0.143 | 0.42 | 74 | 39 |
| | SoilpH | | | -0.051 | -0.167 | -0.086 | -0.051 | -0.023 | 0.053 | 37 | 14 |
| | Ydays | | | -0.004 | -0.009 | -0.005 | -0.004 | -0.003 | 0.005 | 41 | 33 |
| | Year | | | -0.021 | -0.059 | -0.033 | -0.021 | -0.001 | 0.044 | 47 | 19 |

| Species | Variable | Original model coefficient | Original model p | Median | Q 0% | Q 25% | Q 50% | Q 75% | Q 100% | Model Counts | P < 0.05 counts |
|---|---|---|---|---|---|---|---|---|---|---|---|
| *Bothriochloa barbinodis* | AnnMinTemp | | | 0.111 | 0.028 | 0.088 | 0.111 | 0.136 | 0.245 | 86 | 65 |
| | Aspect | | | 0.03 | -0.145 | 0.003 | 0.03 | 0.058 | 0.123 | 111 | 13 |
| | AWC | | | 0.065 | -0.107 | 0.048 | 0.065 | 0.082 | 0.18 | 64 | 23 |
| | clay | | | 0.018 | -0.146 | -0.021 | 0.018 | 0.059 | 0.176 | 75 | 18 |
| | DeltFallMax | | | 0.003 | -0.169 | -0.038 | 0.003 | 0.039 | 0.118 | 14 | 2 |
| | DeltFallMin | | | 0.082 | 0.023 | 0.059 | 0.082 | 0.097 | 0.166 | 63 | 30 |
| | DeltPrecip | | | 0.079 | -0.119 | -0.031 | 0.079 | 0.095 | 0.182 | 6 | 3 |
| | DeltSpringMax | | | -0.08 | -0.164 | -0.095 | -0.08 | -0.055 | 0.088 | 29 | 13 |
| | heatload | | | -0.07 | -0.236 | -0.101 | -0.07 | -0.049 | 0.068 | 105 | 46 |
| | organic | | | 0.089 | -0.026 | 0.069 | 0.089 | 0.12 | 0.222 | 68 | 39 |
| | packm | | | -0.083 | -0.139 | -0.098 | -0.083 | -0.056 | 0.112 | 30 | 16 |
| | SixMonthPrecip | | | 0.108 | 0.04 | 0.088 | 0.108 | 0.13 | 0.173 | 20 | 13 |
| | Slope | | | 0.028 | -0.06 | -0.015 | 0.028 | 0.069 | 0.123 | 19 | 3 |
| | SoilpH | | | -0.057 | -0.195 | -0.076 | -0.057 | -0.034 | 0.078 | 44 | 8 |

| Species | Variable | Original model coefficient | Original model p | Median | Q 0% | Q 25% | Q 50% | Q 75% | Q 100% | Model Counts | P < 0.05 counts |
|---|---|---|---|---|---|---|---|---|---|---|---|
| | Ydays | | | 0.004 | -0.005 | 0.003 | 0.004 | 0.005 | 0.01 | 68 | 54 |
| | Year | | | -0.023 | -0.038 | -0.03 | -0.023 | -0.007 | 0.021 | 10 | 5 |
| *Bouteloua barbata* | AnnMinTemp | | | -0.063 | -0.229 | -0.099 | -0.063 | -0.028 | 0.176 | 17 | 7 |
| | Aspect | | | -0.12 | -0.25 | -0.138 | -0.12 | -0.092 | 0.031 | 65 | 47 |
| | AWC | | | 0.113 | -0.19 | 0.079 | 0.113 | 0.146 | 0.246 | 128 | 82 |
| | clay | 0.121 | 0.001 | 0.058 | -0.363 | 0.006 | 0.058 | 0.178 | 0.576 | 374 | 127 |
| | DeltFallMax | | | 0.039 | -0.163 | -0.021 | 0.039 | 0.111 | 0.418 | 62 | 18 |
| | DeltFallMin | | | -0.096 | -0.432 | -0.142 | -0.096 | -0.05 | 0.113 | 45 | 20 |
| | DeltPrecip | | | 0.091 | -0.221 | 0.043 | 0.091 | 0.127 | 0.664 | 64 | 21 |
| | DeltSpringMax | | | 0.113 | -0.433 | 0.051 | 0.113 | 0.166 | 0.348 | 193 | 84 |
| | heatload | 0.165 | <0.001 | 0.14 | -0.04 | 0.104 | 0.14 | 0.181 | 0.349 | 276 | 226 |
| | organic | | | 0.057 | -0.177 | -0.006 | 0.057 | 0.147 | 0.463 | 16 | 3 |
| | packm | | | 0.103 | -0.304 | 0.081 | 0.103 | 0.137 | 0.291 | 70 | 48 |
| | SixMonthPrecip | | | 0.037 | -0.14 | -0.01 | 0.037 | 0.098 | 0.282 | 79 | 16 |
| | Slope | | | -0.041 | -0.118 | -0.091 | -0.041 | -0.019 | -0.019 | 5 | 2 |

| Species | Variable | Original model coefficient | Original model p | Median | Q 0% | Q 25% | Q 50% | Q 75% | Q 100% | Model Counts | P < 0.05 counts |
|---|---|---|---|---|---|---|---|---|---|---|---|
| | SoilpH | | | 0.114 | -0.039 | 0.066 | 0.114 | 0.159 | 0.603 | 65 | 28 |
| | Ydays | | | -0.006 | -0.013 | -0.007 | -0.006 | -0.004 | 0 | 22 | 14 |
| | Year | | | 0.003 | -0.07 | -0.045 | 0.003 | 0.025 | 0.071 | 8 | 1 |
| *Bouteloua curtipendula* | AnnMinTemp | | | -0.102 | -0.314 | -0.167 | -0.102 | -0.053 | 0.227 | 22 | 12 |
| | Aspect | | | -0.055 | -0.342 | -0.127 | -0.055 | -0.026 | 0.359 | 25 | 9 |
| | AWC | | | 0.103 | -0.097 | 0.016 | 0.103 | 0.177 | 0.381 | 60 | 20 |
| | clay | | | -0.136 | -0.566 | -0.23 | -0.136 | -0.065 | 0.277 | 44 | 25 |
| | DeltFallMax | 0.03 | 0.683 | 0.014 | -0.777 | -0.082 | 0.014 | 0.137 | 0.62 | 97 | 21 |
| | DeltFallMin | | | 0.03 | -0.171 | -0.032 | 0.03 | 0.093 | 0.327 | 31 | 3 |
| | DeltPrecip | -0.133 | 0.074 | -0.079 | -0.68 | -0.203 | -0.079 | 0.02 | 0.371 | 204 | 59 |
| | DeltSpringMax | | | -0.068 | -0.464 | -0.192 | -0.068 | 0.199 | 0.736 | 22 | 12 |
| | heatload | | | 0.175 | -0.12 | 0.074 | 0.175 | 0.238 | 0.513 | 31 | 18 |
| | organic | | | -0.02 | -0.655 | -0.299 | -0.02 | 0.02 | 0.287 | 17 | 4 |
| | packm | -0.098 | 0.024 | -0.077 | -0.305 | -0.138 | -0.077 | 0.005 | 0.093 | 31 | 9 |

| Species | Variable | Original model coefficient | Original model p | Median | Q 0% | Q 25% | Q 50% | Q 75% | Q 100% | Model Counts | P < 0.05 counts |
|---|---|---|---|---|---|---|---|---|---|---|---|
| | SixMonthPrecip | | | 0.144 | -0.188 | 0.083 | 0.144 | 0.208 | 0.402 | 60 | 31 |
| | Slope | | | -0.059 | -0.295 | -0.101 | -0.059 | 0.017 | 0.224 | 14 | 2 |
| | SoilpH | | | 0.104 | -0.352 | -0.06 | 0.104 | 0.186 | 0.382 | 18 | 8 |
| | Ydays | | | -0.002 | -0.016 | -0.005 | -0.002 | 0.002 | 0.007 | 23 | 8 |
| | Year | 0.147 | <0.001 | 0.162 | -0.038 | 0.114 | 0.162 | 0.203 | 0.319 | 487 | 472 |
| *Bouteloua gracilis* | AnnMinTemp | | | 0.115 | -0.05 | 0.078 | 0.115 | 0.141 | 0.316 | 74 | 45 |
| | Aspect | | | 0.111 | -0.08 | 0.081 | 0.111 | 0.136 | 0.199 | 23 | 14 |
| | AWC | | | -0.062 | -0.227 | -0.105 | -0.062 | -0.032 | 0.155 | 46 | 18 |
| | clay | | | -0.054 | -0.317 | -0.138 | -0.054 | 0.017 | 0.155 | 34 | 10 |
| | DeltFallMax | | | -0.069 | -0.268 | -0.11 | -0.069 | 0.038 | 0.187 | 25 | 11 |
| | DeltFallMin | | | 0.056 | -0.212 | -0.05 | 0.056 | 0.1 | 0.361 | 29 | 9 |
| | DeltPrecip | | | 0.127 | -0.128 | 0.081 | 0.127 | 0.176 | 0.381 | 56 | 39 |
| | DeltSpringMax | 0.003 | 0.928 | 0.093 | -0.099 | 0.037 | 0.093 | 0.141 | 0.339 | 31 | 15 |
| | heatload | | | 0.089 | -0.172 | 0.031 | 0.089 | 0.162 | 0.295 | 31 | 17 |
| | organic | | | -0.032 | -0.97 | -0.272 | -0.032 | 0.029 | 0.546 | 41 | 13 |

| Species | Variable | Original model coefficient | Original model p | Median | Q 0% | Q 25% | Q 50% | Q 75% | Q 100% | Model Counts | P < 0.05 counts |
|---|---|---|---|---|---|---|---|---|---|---|---|
| | packm | | | -0.063 | -0.362 | -0.096 | -0.063 | -0.022 | 0.215 | 75 | 24 |
| | SixMonthPrecip | | | 0.114 | -0.117 | 0.067 | 0.114 | 0.179 | 0.397 | 34 | 19 |
| | Slope | | | 0.05 | -0.225 | 0.003 | 0.05 | 0.11 | 0.682 | 31 | 7 |
| | SoilpH | | | 0.016 | -0.221 | -0.071 | 0.016 | 0.067 | 0.253 | 21 | 6 |
| | Ydays | 0.002 | 0.020 | 0.004 | -0.005 | 0.003 | 0.004 | 0.006 | 0.017 | 78 | 48 |
| | Year | 0.01 | 0.248 | 0.031 | -0.038 | 0.022 | 0.031 | 0.046 | 0.083 | 63 | 33 |
| *Bouteloua hirsuta* | AnnMinTemp | | | 0.073 | -0.282 | -0.081 | 0.073 | 0.156 | 0.22 | 17 | 10 |
| | Aspect | | | 0.125 | -0.119 | -0.033 | 0.125 | 0.155 | 0.249 | 31 | 25 |
| | AWC | | | 0.083 | -0.151 | 0.007 | 0.083 | 0.143 | 0.243 | 127 | 63 |
| | clay | | | 0.069 | -0.281 | 0.031 | 0.069 | 0.113 | 0.159 | 82 | 29 |
| | DeltFallMax | | | -0.054 | -0.613 | -0.098 | -0.054 | 0.008 | 0.229 | 74 | 19 |
| | DeltFallMin | | | 0.099 | -0.087 | 0.031 | 0.099 | 0.177 | 0.325 | 25 | 14 |
| | DeltPrecip | | | 0.028 | -0.183 | -0.03 | 0.028 | 0.071 | 0.374 | 49 | 13 |
| | DeltSpringMax | | | -0.084 | -0.208 | -0.101 | -0.084 | -0.058 | 0.098 | 28 | 15 |
| | heatload | | | 0.054 | -0.093 | -0.009 | 0.054 | 0.161 | 0.267 | 8 | 5 |

| Species | Variable | Original model coefficient | Original model p | Median | Q 0% | Q 25% | Q 50% | Q 75% | Q 100% | Model Counts | P < 0.05 counts |
|---|---|---|---|---|---|---|---|---|---|---|---|
| | organic | | | 0.112 | -0.193 | 0.065 | 0.112 | 0.137 | 0.208 | 70 | 47 |
| | packm | | | -0.111 | -0.599 | -0.142 | -0.111 | -0.087 | 0.032 | 210 | 159 |
| | SixMonthPrecip | | | 0.076 | -0.042 | 0.066 | 0.076 | 0.132 | 0.495 | 9 | 3 |
| | Slope | | | 0.065 | -0.152 | -0.09 | 0.065 | 0.085 | 0.24 | 11 | 6 |
| | SoilpH | | | -0.048 | -0.265 | -0.098 | -0.048 | 0.022 | 0.181 | 106 | 33 |
| | Ydays | | | -0.007 | -0.012 | -0.01 | -0.007 | -0.004 | -0.003 | 12 | 5 |
| | Year | -0.008 | 0.511 | -0.005 | -0.054 | -0.018 | -0.005 | 0.005 | 0.04 | 158 | 32 |
| *Camissonia brevipes* | AnnMinTemp | | | 0.197 | -0.352 | 0.145 | 0.197 | 0.281 | 0.864 | 157 | 105 |
| | Aspect | | | -0.16 | -0.509 | -0.229 | -0.16 | -0.107 | 0.088 | 196 | 100 |
| | AWC | | | 0.151 | 0.036 | 0.076 | 0.151 | 0.241 | 0.461 | 7 | 3 |
| | clay | | | 0.038 | -0.211 | 0.018 | 0.038 | 0.056 | 0.142 | 23 | 2 |
| | DeltFallMax | | | 0.196 | -0.115 | 0.138 | 0.196 | 0.257 | 0.645 | 196 | 131 |
| | DeltFallMin | | | -0.085 | -0.328 | -0.137 | -0.085 | -0.008 | 0.399 | 162 | 53 |
| | DeltPrecip | | | 0.001 | -0.319 | -0.041 | 0.001 | 0.02 | 0.093 | 17 | 4 |

| Species | Variable | Original model coefficient | Original model p | Median | Q 0% | Q 25% | Q 50% | Q 75% | Q 100% | Model Counts | P < 0.05 counts |
|---|---|---|---|---|---|---|---|---|---|---|---|
| | DeltSpringMax | | | 0.116 | -0.04 | 0.03 | 0.116 | 0.178 | 0.322 | 17 | 7 |
| | heatload | | | 0.095 | -0.064 | -0.024 | 0.095 | 0.153 | 0.452 | 10 | 1 |
| | organic | | | 0.011 | -0.198 | -0.045 | 0.011 | 0.046 | 0.536 | 37 | 6 |
| | packm | | | -0.157 | -0.508 | -0.319 | -0.157 | -0.077 | 0.13 | 28 | 21 |
| | SixMonthPrecip | | | 0.122 | -0.16 | 0.069 | 0.122 | 0.175 | 0.277 | 35 | 18 |
| | Slope | | | 0.148 | -0.192 | 0.087 | 0.148 | 0.187 | 0.395 | 29 | 16 |
| | SoilpH | | | -0.137 | -0.679 | -0.252 | -0.137 | -0.081 | 0.274 | 172 | 53 |
| | Ydays | | | 0.001 | -0.007 | 0 | 0.001 | 0.002 | 0.011 | 34 | 8 |
| | Year | -0.011 | 0.712 | 0.069 | -0.068 | 0.022 | 0.069 | 0.125 | 0.167 | 11 | 6 |
| *Cercocarpus ledifolius* | AnnMinTemp | 0.004 | 0.860 | 0.026 | -0.156 | -0.011 | 0.026 | 0.062 | 0.251 | 193 | 67 |
| | Aspect | -0.025 | 0.272 | -0.068 | -0.15 | -0.097 | -0.068 | -0.044 | 0.046 | 68 | 46 |
| | AWC | | | 0.006 | -0.133 | -0.041 | 0.006 | 0.055 | 0.141 | 47 | 20 |
| | clay | 0.042 | 0.097 | 0.05 | -0.199 | 0.022 | 0.05 | 0.07 | 0.148 | 79 | 37 |
| | DeltFallMax | | | -0.039 | -0.254 | -0.066 | -0.039 | 0 | 0.19 | 36 | 13 |

| Species | Variable | Original model coefficient | Original model p | Median | Q 0% | Q 25% | Q 50% | Q 75% | Q 100% | Model Counts | P < 0.05 counts |
|---|---|---|---|---|---|---|---|---|---|---|---|
| | DeltFallMin | 0.042 | 0.088 | 0.053 | -0.195 | 0.027 | 0.053 | 0.09 | 0.199 | 198 | 96 |
| | DeltPrecip | | | 0.05 | -0.108 | 0.022 | 0.05 | 0.106 | 0.229 | 36 | 14 |
| | DeltSpringMax | | | -0.03 | -0.269 | -0.054 | -0.03 | 0.002 | 0.227 | 56 | 17 |
| | heatload | | | -0.054 | -0.148 | -0.086 | -0.054 | -0.008 | 0.092 | 49 | 12 |
| | organic | 0.011 | 0.664 | 0.043 | -0.259 | -0.043 | 0.043 | 0.072 | 0.233 | 72 | 46 |
| | packm | | | -0.02 | -0.16 | -0.104 | -0.02 | 0.016 | 0.058 | 21 | 11 |
| | SixMonthPrecip | | | -0.002 | -0.253 | -0.079 | -0.002 | 0.027 | 0.139 | 31 | 9 |
| | Slope | | | -0.039 | -0.088 | -0.065 | -0.039 | -0.029 | 0.104 | 30 | 20 |
| | SoilpH | -0.04 | 0.105 | -0.067 | -0.218 | -0.092 | -0.067 | -0.035 | 0.054 | 97 | 52 |
| | Ydays | | | 0.001 | -0.002 | -0.001 | 0.001 | 0.002 | 0.005 | 29 | 8 |
| | Year | 0.002 | 0.819 | 0.015 | -0.061 | 0.001 | 0.015 | 0.026 | 0.089 | 91 | 40 |
| *Cercocarpus montanus* | AnnMinTemp | | | 0.104 | -0.418 | 0.045 | 0.104 | 0.15 | 0.21 | 43 | 26 |
| | Aspect | | | -0.031 | -0.078 | -0.049 | -0.031 | -0.009 | 0.042 | 21 | 3 |
| | AWC | | | -0.055 | -0.184 | -0.078 | -0.055 | -0.036 | 0.047 | 130 | 49 |

| Species | Variable | Original model coefficient | Original model p | Median | Q 0% | Q 25% | Q 50% | Q 75% | Q 100% | Model Counts | P < 0.05 counts |
|---|---|---|---|---|---|---|---|---|---|---|---|
| | clay | | | -0.061 | -0.145 | -0.083 | -0.061 | -0.045 | 0 | 16 | 7 |
| | DeltFallMax | | | 0.001 | -0.117 | -0.028 | 0.001 | 0.037 | 0.123 | 31 | 5 |
| | DeltFallMin | | | -0.037 | -0.129 | -0.06 | -0.037 | -0.008 | 0.024 | 10 | 3 |
| | DeltPrecip | 0.066 | 0.105 | 0.067 | -0.195 | 0.038 | 0.067 | 0.09 | 0.458 | 272 | 109 |
| | DeltSpringMax | | | 0.007 | -0.095 | -0.02 | 0.007 | 0.024 | 0.053 | 7 | 1 |
| | heatload | | | -0.001 | -0.254 | -0.086 | -0.001 | 0.02 | 0.441 | 25 | 9 |
| | organic | | | -0.008 | -0.256 | -0.074 | -0.008 | 0.024 | 0.076 | 22 | 5 |
| | packm | | | 0.058 | -0.094 | 0.038 | 0.058 | 0.078 | 0.145 | 60 | 39 |
| | SixMonthPrecip | | | 0.085 | -0.053 | 0.036 | 0.085 | 0.105 | 0.211 | 24 | 16 |
| | Slope | | | -0.038 | -0.273 | -0.085 | -0.038 | -0.014 | 0.143 | 70 | 27 |
| | SoilpH | | | -0.053 | -0.186 | -0.081 | -0.053 | -0.011 | 0.078 | 68 | 30 |
| | Ydays | | | 0.003 | -0.017 | 0.001 | 0.003 | 0.003 | 0.005 | 60 | 33 |
| | Year | | | 0.014 | -0.01 | 0.001 | 0.014 | 0.025 | 0.033 | 4 | 1 |
| *Chaenactis douglasii* | AnnMinTemp | | | 0.078 | -0.007 | 0.062 | 0.078 | 0.108 | 0.14 | 42 | 26 |

| Species | Variable | Original model coefficient | Original model p | Median | Q 0% | Q 25% | Q 50% | Q 75% | Q 100% | Model Counts | P < 0.05 counts |
|---|---|---|---|---|---|---|---|---|---|---|---|
| | Aspect | | | -0.011 | -0.165 | -0.08 | -0.011 | 0.073 | 0.167 | 24 | 12 |
| | AWC | | | 0.099 | -0.031 | 0.068 | 0.099 | 0.121 | 0.168 | 39 | 27 |
| | clay | | | 0.066 | -0.08 | 0.05 | 0.066 | 0.085 | 0.154 | 42 | 18 |
| | DeltFallMax | | | 0.08 | -0.002 | 0.059 | 0.08 | 0.11 | 0.164 | 102 | 65 |
| | DeltFallMin | | | -0.018 | -0.142 | -0.074 | -0.018 | 0.033 | 0.162 | 25 | 12 |
| | DeltPrecip | | | -0.068 | -0.212 | -0.093 | -0.068 | -0.056 | 0.095 | 56 | 30 |
| | DeltSpringMax | | | 0.081 | -0.037 | 0.064 | 0.081 | 0.1 | 0.195 | 50 | 29 |
| | heatload | | | 0.015 | -0.084 | -0.003 | 0.015 | 0.066 | 0.098 | 13 | 5 |
| | organic | | | -0.013 | -0.413 | -0.135 | -0.013 | 0.033 | 0.339 | 24 | 3 |
| | packm | | | -0.056 | -0.255 | -0.071 | -0.056 | -0.025 | 0.094 | 40 | 19 |
| | SixMonthPrecip | | | 0.014 | -0.137 | -0.037 | 0.014 | 0.074 | 0.204 | 30 | 11 |
| | Slope | | | -0.005 | -0.1 | -0.042 | -0.005 | 0.039 | 0.233 | 11 | 4 |
| | SoilpH | | | -0.032 | -0.15 | -0.061 | -0.032 | 0.013 | 0.174 | 59 | 17 |
| | Ydays | | | 0.001 | -0.01 | -0.005 | 0.001 | 0.003 | 0.006 | 47 | 33 |
| | Year | | | -0.01 | -0.066 | -0.02 | -0.01 | 0 | 0.024 | 29 | 7 |

| Species | Variable | Original model coefficient | Original model p | Median | Q 0% | Q 25% | Q 50% | Q 75% | Q 100% | Model Counts | P < 0.05 counts |
|---|---|---|---|---|---|---|---|---|---|---|---|
| *Chaenactis fremontii* | AnnMinTemp | | | 0.147 | -0.227 | 0.055 | 0.147 | 0.206 | 0.312 | 32 | 24 |
| | Aspect | | | 0.017 | -0.104 | -0.041 | 0.017 | 0.044 | 0.079 | 16 | 5 |
| | AWC | | | 0.07 | -0.112 | 0.028 | 0.07 | 0.109 | 0.246 | 49 | 23 |
| | clay | | | -0.097 | -0.326 | -0.14 | -0.097 | -0.077 | 0.076 | 71 | 63 |
| | DeltFallMax | | | -0.068 | -0.146 | -0.086 | -0.068 | -0.052 | -0.016 | 22 | 14 |
| | DeltFallMin | | | 0.051 | -0.081 | 0.026 | 0.051 | 0.081 | 0.226 | 94 | 40 |
| | DeltPrecip | | | -0.039 | -0.168 | -0.073 | -0.039 | 0.005 | 0.125 | 45 | 9 |
| | DeltSpringMax | | | 0.051 | -0.118 | -0.007 | 0.051 | 0.085 | 0.173 | 22 | 8 |
| | heatload | | | 0.104 | -0.36 | 0.04 | 0.104 | 0.193 | 0.668 | 58 | 22 |
| | organic | | | 0.049 | -0.277 | 0.004 | 0.049 | 0.088 | 0.5 | 68 | 20 |
| | packm | | | 0.042 | -0.09 | 0.013 | 0.042 | 0.069 | 0.155 | 44 | 9 |
| | SixMonthPrecip | 0.064 | 0.055 | 0.062 | -0.17 | 0.036 | 0.062 | 0.095 | 0.322 | 257 | 82 |
| | Slope | -0.111 | 0.001 | -0.106 | -0.559 | -0.151 | -0.106 | -0.048 | 0.15 | 153 | 73 |
| | SoilpH | -0.106 | 0.003 | -0.074 | -0.446 | -0.108 | -0.074 | -0.033 | 0.53 | 131 | 56 |

| Species | Variable | Original model coefficient | Original model p | Median | Q 0% | Q 25% | Q 50% | Q 75% | Q 100% | Model Counts | P < 0.05 counts |
|---|---|---|---|---|---|---|---|---|---|---|---|
| | Ydays | | | -0.004 | -0.01 | -0.008 | -0.004 | -0.001 | 0.002 | 31 | 16 |
| | Year | | | 0.033 | -0.056 | 0.019 | 0.033 | 0.042 | 0.07 | 32 | 15 |
| *Chaenactis stevioides* | AnnMinTemp | | | 0.049 | -0.323 | -0.023 | 0.049 | 0.062 | 0.097 | 31 | 17 |
| | Aspect | | | 0.037 | -0.066 | 0.022 | 0.037 | 0.071 | 0.112 | 25 | 15 |
| | AWC | | | -0.012 | -0.136 | -0.02 | -0.012 | 0.01 | 0.054 | 11 | 2 |
| | clay | | | 0.057 | 0.008 | 0.039 | 0.057 | 0.08 | 0.247 | 50 | 19 |
| | DeltFallMax | | | -0.065 | -0.123 | -0.076 | -0.065 | -0.05 | -0.033 | 26 | 18 |
| | DeltFallMin | | | 0.027 | -0.061 | -0.008 | 0.027 | 0.047 | 0.177 | 59 | 19 |
| | DeltPrecip | | | -0.009 | -0.184 | -0.045 | -0.009 | 0.02 | 0.103 | 46 | 6 |
| | DeltSpringMax | | | -0.047 | -0.143 | -0.066 | -0.047 | -0.03 | 0.079 | 46 | 16 |
| | heatload | | | -0.041 | -0.072 | -0.05 | -0.041 | -0.028 | 0.011 | 14 | 3 |
| | organic | | | 0.016 | 0.016 | 0.016 | 0.016 | 0.016 | 0.016 | 1 | 0 |
| | packm | | | 0.041 | -0.061 | 0.021 | 0.041 | 0.059 | 0.277 | 146 | 45 |
| | SixMonthPrecip | | | 0.052 | 0.011 | 0.029 | 0.052 | 0.09 | 0.173 | 36 | 15 |

| Species | Variable | Original model coefficient | Original model p | Median | Q 0% | Q 25% | Q 50% | Q 75% | Q 100% | Model Counts | P < 0.05 counts |
|---|---|---|---|---|---|---|---|---|---|---|---|
| | Slope | | | -0.036 | -0.164 | -0.051 | -0.036 | -0.019 | 0.082 | 39 | 9 |
| | SoilpH | 0.065 | 0.002 | 0.061 | -0.081 | 0.046 | 0.061 | 0.073 | 0.146 | 319 | 190 |
| | Ydays | | | 0 | -0.008 | -0.001 | 0 | 0.001 | 0.002 | 8 | 2 |
| | Year | | | -0.001 | -0.04 | -0.018 | -0.001 | 0.016 | 0.044 | 12 | 6 |
| *Chrysothamnus viscidiflorus* | AnnMinTemp | | | 0.053 | -0.134 | 0.015 | 0.053 | 0.086 | 0.2 | 79 | 25 |
| | Aspect | | | -0.11 | -0.214 | -0.139 | -0.11 | -0.083 | 0.08 | 38 | 25 |
| | AWC | | | 0.106 | -0.196 | 0.067 | 0.106 | 0.157 | 0.304 | 51 | 28 |
| | clay | 0.075 | 0.013 | 0.108 | -0.266 | 0.076 | 0.108 | 0.148 | 0.385 | 111 | 69 |
| | DeltFallMax | | | 0.077 | -0.201 | 0.029 | 0.077 | 0.111 | 0.188 | 37 | 19 |
| | DeltFallMin | | | 0.029 | -0.136 | -0.025 | 0.029 | 0.107 | 0.246 | 30 | 11 |
| | DeltPrecip | 0.055 | 0.072 | 0.085 | -0.251 | 0.042 | 0.085 | 0.121 | 0.194 | 116 | 75 |
| | DeltSpringMax | | | 0.117 | -0.044 | 0.079 | 0.117 | 0.147 | 0.247 | 80 | 52 |
| | heatload | | | 0.109 | -0.058 | 0.05 | 0.109 | 0.159 | 0.276 | 54 | 37 |
| | organic | | | 0.074 | -0.069 | 0.012 | 0.074 | 0.162 | 0.857 | 43 | 8 |

| Species | Variable | Original model coefficient | Original model p | Median | Q 0% | Q 25% | Q 50% | Q 75% | Q 100% | Model Counts | P < 0.05 counts |
|---|---|---|---|---|---|---|---|---|---|---|---|
| | packm | | | -0.077 | -0.366 | -0.097 | -0.077 | -0.036 | 0.117 | 43 | 21 |
| | SixMonthPrecip | | | -0.054 | -0.263 | -0.082 | -0.054 | -0.003 | 0.088 | 37 | 13 |
| | Slope | | | -0.02 | -0.327 | -0.142 | -0.02 | 0.043 | 0.13 | 16 | 8 |
| | SoilpH | | | -0.087 | -0.243 | -0.168 | -0.087 | -0.047 | 0.136 | 31 | 22 |
| | Ydays | | | 0.003 | -0.004 | 0.001 | 0.003 | 0.004 | 0.009 | 100 | 45 |
| | Year | | | -0.017 | -0.062 | -0.027 | -0.017 | -0.01 | 0.04 | 74 | 31 |
| *Cleome lutea* | AnnMinTemp | | | -0.007 | -0.084 | -0.034 | -0.007 | 0.03 | 0.087 | 33 | 10 |
| | Aspect | 0.046 | 0.004 | 0.06 | 0.011 | 0.045 | 0.06 | 0.073 | 0.202 | 183 | 139 |
| | AWC | | | 0.029 | -0.072 | 0.016 | 0.029 | 0.052 | 0.09 | 40 | 15 |
| | clay | | | -0.052 | -0.116 | -0.065 | -0.052 | -0.005 | 0.08 | 63 | 29 |
| | DeltFallMax | | | -0.045 | -0.115 | -0.064 | -0.045 | -0.018 | 0.047 | 28 | 14 |
| | DeltFallMin | | | 0.018 | -0.105 | -0.018 | 0.018 | 0.047 | 0.145 | 82 | 26 |
| | DeltPrecip | | | 0.039 | -0.092 | 0.019 | 0.039 | 0.058 | 0.157 | 71 | 32 |
| | DeltSpringMax | | | -0.02 | -0.132 | -0.051 | -0.02 | 0.006 | 0.093 | 27 | 10 |
| | heatload | | | 0.035 | -0.25 | 0.014 | 0.035 | 0.054 | 0.141 | 81 | 36 |

| Species | Variable | Original model coefficient | Original model p | Median | Q 0% | Q 25% | Q 50% | Q 75% | Q 100% | Model Counts | P < 0.05 counts |
|---|---|---|---|---|---|---|---|---|---|---|---|
| | organic | | | 0.041 | -0.065 | 0.016 | 0.041 | 0.06 | 0.168 | 108 | 36 |
| | packm | | | 0.06 | -0.067 | 0.021 | 0.06 | 0.094 | 0.236 | 117 | 54 |
| | SixMonthPrecip | | | 0.036 | -0.113 | 0.002 | 0.036 | 0.05 | 0.131 | 81 | 30 |
| | Slope | | | 0.057 | -0.065 | 0.023 | 0.057 | 0.085 | 0.269 | 98 | 52 |
| | SoilpH | -0.041 | 0.034 | -0.051 | -0.14 | -0.072 | -0.051 | -0.026 | 0.297 | 164 | 79 |
| | Ydays | | | -0.002 | -0.003 | -0.002 | -0.002 | -0.001 | 0 | 69 | 45 |
| | Year | | | -0.012 | -0.033 | -0.015 | -0.012 | -0.007 | 0 | 24 | 15 |
| *Cleome serrulata* | AnnMinTemp | | | 0.06 | -0.051 | 0.042 | 0.06 | 0.074 | 0.202 | 173 | 126 |
| | Aspect | 0.033 | 0.066 | 0.04 | -0.027 | 0.021 | 0.04 | 0.048 | 0.074 | 36 | 18 |
| | AWC | | | -0.02 | -0.074 | -0.03 | -0.02 | -0.005 | 0.085 | 27 | 5 |
| | clay | | | -0.038 | -0.07 | -0.047 | -0.038 | -0.022 | 0.007 | 20 | 7 |
| | DeltFallMax | | | 0.03 | -0.073 | 0.015 | 0.03 | 0.045 | 0.074 | 31 | 14 |
| | DeltFallMin | | | -0.023 | -0.131 | -0.057 | -0.023 | 0.012 | 0.094 | 114 | 46 |
| | DeltPrecip | | | -0.044 | -0.187 | -0.065 | -0.044 | -0.027 | 0.047 | 24 | 10 |
| | DeltSpringMax | | | -0.029 | -0.136 | -0.059 | -0.029 | -0.001 | 0.111 | 72 | 20 |

| Species | Variable | Original model coefficient | Original model p | Median | Q 0% | Q 25% | Q 50% | Q 75% | Q 100% | Model Counts | P < 0.05 counts |
|---|---|---|---|---|---|---|---|---|---|---|---|
| | heatload | | | -0.028 | -0.098 | -0.049 | -0.028 | -0.001 | 0.054 | 56 | 21 |
| | organic | | | 0.015 | -0.239 | -0.011 | 0.015 | 0.076 | 0.249 | 59 | 14 |
| | packm | | | 0.045 | -0.038 | 0.025 | 0.045 | 0.059 | 0.106 | 46 | 23 |
| | SixMonthPrecip | | | 0.044 | -0.019 | 0.034 | 0.044 | 0.065 | 0.144 | 84 | 36 |
| | Slope | | | 0.023 | -0.049 | 0.016 | 0.023 | 0.04 | 0.068 | 18 | 6 |
| | SoilpH | -0.014 | 0.404 | 0.016 | -0.098 | -0.04 | 0.016 | 0.078 | 0.179 | 61 | 31 |
| | Ydays | | | -0.001 | -0.005 | -0.003 | -0.001 | -0.001 | 0.001 | 44 | 16 |
| | Year | | | 0.004 | -0.043 | -0.001 | 0.004 | 0.011 | 0.039 | 160 | 39 |
| *Crepis acuminata* | AnnMinTemp | | | 0.113 | -0.028 | 0.088 | 0.113 | 0.146 | 0.197 | 111 | 93 |
| | Aspect | | | -0.092 | -0.2 | -0.125 | -0.092 | -0.058 | 0.043 | 23 | 14 |
| | AWC | | | -0.041 | -0.21 | -0.097 | -0.041 | 0.001 | 0.122 | 49 | 15 |
| | clay | | | 0.069 | -0.138 | 0.011 | 0.069 | 0.121 | 0.267 | 75 | 33 |
| | DeltFallMax | | | -0.1 | -0.298 | -0.13 | -0.1 | -0.051 | 0.016 | 68 | 34 |
| | DeltFallMin | | | -0.068 | -0.167 | -0.108 | -0.068 | 0.047 | 0.156 | 18 | 10 |
| | DeltPrecip | | | 0.073 | -0.172 | 0.013 | 0.073 | 0.132 | 0.362 | 31 | 16 |

| Species | Variable | Original model coefficient | Original model p | Median | Q 0% | Q 25% | Q 50% | Q 75% | Q 100% | Model Counts | P < 0.05 counts |
|---|---|---|---|---|---|---|---|---|---|---|---|
| | DeltSpringMax | | | 0.014 | -0.084 | -0.026 | 0.014 | 0.089 | 0.16 | 15 | 6 |
| | heatload | | | -0.06 | -0.298 | -0.107 | -0.06 | 0.033 | 0.123 | 20 | 8 |
| | organic | | | -0.115 | -0.217 | -0.143 | -0.115 | -0.086 | 0.072 | 99 | 75 |
| | packm | | | -0.068 | -0.361 | -0.129 | -0.068 | -0.018 | 0.064 | 30 | 5 |
| | SixMonthPrecip | | | -0.066 | -0.17 | -0.1 | -0.066 | -0.003 | 0.182 | 26 | 14 |
| | Slope | | | 0.07 | -0.143 | 0.009 | 0.07 | 0.108 | 0.171 | 32 | 13 |
| | SoilpH | | | -0.041 | -0.144 | -0.092 | -0.041 | 0.023 | 0.049 | 13 | 3 |
| | Ydays | 0 | 0.899 | -0.002 | -0.014 | -0.004 | -0.002 | 0 | 0.004 | 173 | 46 |
| | Year | | | -0.023 | -0.044 | -0.029 | -0.023 | -0.012 | 0.017 | 26 | 6 |
| *Elymus elymoides* | AnnMinTemp | 0.07 | 0.004 | 0.135 | -0.083 | 0.082 | 0.135 | 0.192 | 0.449 | 80 | 48 |
| | Aspect | | | -0.101 | -0.238 | -0.132 | -0.101 | 0.058 | 0.159 | 21 | 13 |
| | AWC | | | 0.055 | -0.25 | -0.105 | 0.055 | 0.116 | 0.361 | 25 | 15 |
| | clay | | | 0.082 | -0.029 | 0.046 | 0.082 | 0.126 | 0.224 | 38 | 12 |
| | DeltFallMax | | | 0.011 | -0.423 | -0.118 | 0.011 | 0.058 | 0.231 | 34 | 14 |
| | DeltFallMin | | | 0.061 | -0.249 | -0.004 | 0.061 | 0.099 | 0.222 | 18 | 7 |

| Species | Variable | Original model coefficient | Original model p | Median | Q 0% | Q 25% | Q 50% | Q 75% | Q 100% | Model Counts | P < 0.05 counts |
|---|---|---|---|---|---|---|---|---|---|---|---|
| | DeltPrecip | | | -0.068 | -0.203 | -0.115 | -0.068 | -0.053 | 0.226 | 23 | 10 |
| | DeltSpringMax | | | -0.077 | -0.214 | -0.128 | -0.077 | 0.017 | 0.124 | 20 | 7 |
| | heatload | | | -0.059 | -0.179 | -0.115 | -0.059 | -0.019 | 0.082 | 23 | 6 |
| | organic | -0.003 | 0.905 | 0.023 | -0.402 | -0.025 | 0.023 | 0.087 | 0.843 | 119 | 19 |
| | packm | | | 0.057 | -0.194 | 0 | 0.057 | 0.077 | 0.172 | 17 | 7 |
| | SixMonthPrecip | 0.034 | 0.158 | 0.139 | -0.436 | 0.055 | 0.139 | 0.271 | 0.737 | 217 | 88 |
| | Slope | -0.002 | 0.935 | 0.117 | -0.151 | 0.067 | 0.117 | 0.194 | 0.35 | 87 | 41 |
| | SoilpH | -0.141 | <0.001 | -0.166 | -0.439 | -0.204 | -0.166 | -0.121 | 0.095 | 309 | 226 |
| | Ydays | 0.004 | <0.001 | 0.006 | -0.003 | 0.004 | 0.006 | 0.007 | 0.014 | 65 | 44 |
| | Year | | | 0.024 | -0.077 | -0.033 | 0.024 | 0.046 | 0.065 | 18 | 12 |
| *Encelia farinosa* | AnnMinTemp | | | -0.006 | -0.123 | -0.046 | -0.006 | 0.024 | 0.151 | 44 | 10 |
| | Aspect | | | -0.057 | -0.174 | -0.091 | -0.057 | 0.002 | 0.16 | 31 | 21 |
| | AWC | | | 0.063 | -0.101 | 0.028 | 0.063 | 0.08 | 0.133 | 38 | 17 |
| | clay | | | -0.046 | -0.155 | -0.106 | -0.046 | -0.007 | 0.144 | 18 | 2 |
| | DeltFallMax | | | -0.069 | -0.172 | -0.087 | -0.069 | -0.034 | 0.067 | 51 | 22 |

| Species | Variable | Original model coefficient | Original model p | Median | Q 0% | Q 25% | Q 50% | Q 75% | Q 100% | Model Counts | P < 0.05 counts |
|---|---|---|---|---|---|---|---|---|---|---|---|
| | DeltFallMin | | | -0.079 | -0.156 | -0.115 | -0.079 | -0.066 | 0.038 | 18 | 7 |
| | DeltPrecip | | | 0.003 | -0.173 | -0.026 | 0.003 | 0.065 | 0.092 | 12 | 6 |
| | DeltSpringMax | | | -0.064 | -0.172 | -0.09 | -0.064 | -0.035 | 0.031 | 89 | 33 |
| | heatload | | | -0.049 | -0.098 | -0.057 | -0.049 | -0.008 | 0.145 | 14 | 8 |
| | organic | -0.045 | 0.319 | 0.024 | -0.202 | -0.028 | 0.024 | 0.064 | 0.526 | 111 | 15 |
| | packm | | | -0.043 | -0.145 | -0.056 | -0.043 | -0.021 | 0.039 | 39 | 10 |
| | SixMonthPrecip | | | -0.084 | -0.16 | -0.094 | -0.084 | -0.065 | 0.035 | 26 | 18 |
| | Slope | | | 0.04 | -0.057 | 0.001 | 0.04 | 0.063 | 0.1 | 36 | 13 |
| | SoilpH | -0.081 | 0.080 | -0.064 | -0.333 | -0.106 | -0.064 | -0.046 | 0.11 | 167 | 51 |
| | Ydays | | | -0.002 | -0.007 | -0.003 | -0.002 | -0.001 | 0.003 | 109 | 20 |
| | Year | | | 0.02 | -0.039 | 0.011 | 0.02 | 0.026 | 0.045 | 34 | 15 |
| *Encelia virginensis* | AnnMinTemp | -0.103 | 0.053 | -0.08 | -0.429 | -0.136 | -0.08 | 0 | 0.315 | 259 | 102 |
| | Aspect | | | 0.058 | -0.128 | 0.012 | 0.058 | 0.088 | 0.237 | 54 | 19 |
| | AWC | | | 0.065 | -0.29 | 0.024 | 0.065 | 0.12 | 0.299 | 30 | 11 |
| | clay | | | -0.028 | -0.382 | -0.072 | -0.028 | 0.017 | 0.61 | 119 | 29 |

| Species | Variable | Original model coefficient | Original model p | Median | Q 0% | Q 25% | Q 50% | Q 75% | Q 100% | Model Counts | P < 0.05 counts |
|---|---|---|---|---|---|---|---|---|---|---|---|
| | DeltFallMax | | | -0.02 | -0.194 | -0.043 | -0.02 | 0.017 | 0.203 | 60 | 15 |
| | DeltFallMin | | | -0.059 | -0.294 | -0.114 | -0.059 | -0.031 | 0.126 | 58 | 19 |
| | DeltPrecip | | | -0.036 | -0.213 | -0.114 | -0.036 | 0.064 | 0.144 | 51 | 29 |
| | DeltSpringMax | | | 0.054 | -0.191 | 0.002 | 0.054 | 0.125 | 0.768 | 42 | 10 |
| | heatload | | | -0.059 | -0.452 | -0.095 | -0.059 | -0.019 | 0.374 | 76 | 37 |
| | organic | | | 0.03 | -0.449 | -0.061 | 0.03 | 0.111 | 0.28 | 10 | 4 |
| | packm | | | 0.02 | -0.211 | -0.008 | 0.02 | 0.034 | 0.177 | 85 | 14 |
| | SixMonthPrecip | | | 0.015 | -0.134 | -0.055 | 0.015 | 0.044 | 0.105 | 9 | 1 |
| | Slope | | | 0.062 | -0.029 | 0.015 | 0.062 | 0.095 | 0.258 | 30 | 11 |
| | SoilpH | | | -0.069 | -0.241 | -0.12 | -0.069 | -0.04 | 0.166 | 34 | 18 |
| | Ydays | 0 | 0.864 | 0.002 | -0.009 | 0 | 0.002 | 0.004 | 0.016 | 241 | 75 |
| | Year | | | 0.026 | -0.04 | 0.015 | 0.026 | 0.05 | 0.212 | 88 | 47 |
| *Ericameria nauseosa* | AnnMinTemp | | | -0.12 | -0.311 | -0.202 | -0.12 | -0.027 | 0.22 | 42 | 25 |
| | Aspect | | | 0.079 | -0.115 | -0.009 | 0.079 | 0.107 | 0.478 | 11 | 5 |

| Species | Variable | Original model coefficient | Original model p | Median | Q 0% | Q 25% | Q 50% | Q 75% | Q 100% | Model Counts | P < 0.05 counts |
|---|---|---|---|---|---|---|---|---|---|---|---|
| | AWC | | | -0.05 | -1.023 | -0.15 | -0.05 | 0.087 | 0.195 | 20 | 7 |
| | clay | | | 0.097 | -0.093 | 0.052 | 0.097 | 0.167 | 0.629 | 51 | 18 |
| | DeltFallMax | 0.059 | 0.180 | 0.108 | -0.168 | 0.056 | 0.108 | 0.175 | 0.658 | 151 | 60 |
| | DeltFallMin | | | -0.013 | -0.378 | -0.139 | -0.013 | 0.09 | 0.157 | 30 | 8 |
| | DeltPrecip | | | -0.119 | -0.584 | -0.216 | -0.119 | 0.055 | 0.125 | 20 | 9 |
| | DeltSpringMax | | | -0.078 | -0.359 | -0.177 | -0.078 | 0.013 | 0.446 | 33 | 13 |
| | heatload | | | -0.124 | -0.65 | -0.211 | -0.124 | -0.045 | 0.243 | 70 | 38 |
| | organic | 0.09 | 0.023 | 0.13 | -1.111 | 0.068 | 0.13 | 0.378 | 2.137 | 250 | 103 |
| | packm | | | 0.098 | -0.326 | 0.029 | 0.098 | 0.168 | 0.984 | 85 | 29 |
| | SixMonthPrecip | | | 0.072 | -0.181 | -0.014 | 0.072 | 0.244 | 0.542 | 25 | 5 |
| | Slope | | | 0.114 | -0.284 | 0.057 | 0.114 | 0.196 | 0.511 | 101 | 52 |
| | SoilpH | -0.173 | <0.001 | -0.174 | -0.58 | -0.252 | -0.174 | -0.103 | 0.523 | 292 | 176 |
| | Ydays | | | -0.005 | -0.016 | -0.008 | -0.005 | -0.003 | 0.009 | 58 | 28 |
| | Year | -0.06 | <0.001 | -0.071 | -0.257 | -0.096 | -0.071 | -0.053 | 0.006 | 144 | 112 |
| *Erigeron pumilus* | AnnMinTemp | | | -0.197 | -0.732 | -0.27 | -0.197 | -0.115 | 0.83 | 266 | 188 |

| Species | Variable | Original model coefficient | Original model p | Median | Q 0% | Q 25% | Q 50% | Q 75% | Q 100% | Model Counts | P < 0.05 counts |
|---|---|---|---|---|---|---|---|---|---|---|---|
| | Aspect | | | 0.102 | -0.061 | 0.045 | 0.102 | 0.164 | 0.248 | 28 | 12 |
| | AWC | | | -0.045 | -0.19 | -0.163 | -0.045 | 0.14 | 0.294 | 21 | 15 |
| | clay | | | 0.143 | -0.057 | 0.083 | 0.143 | 0.218 | 0.387 | 41 | 18 |
| | DeltFallMax | 0.316 | 0.005 | 0.193 | -1.112 | 0.112 | 0.193 | 0.297 | 0.944 | 208 | 136 |
| | DeltFallMin | | | -0.116 | -0.433 | -0.236 | -0.116 | 0.006 | 0.206 | 15 | 8 |
| | DeltPrecip | | | -0.04 | -0.04 | -0.04 | -0.04 | -0.04 | -0.04 | 1 | 0 |
| | DeltSpringMax | | | 0.184 | -0.405 | 0.093 | 0.184 | 0.28 | 0.626 | 236 | 139 |
| | heatload | | | 0.131 | -0.269 | -0.032 | 0.131 | 0.185 | 0.258 | 21 | 14 |
| | organic | -0.134 | 0.265 | 0.064 | -1.076 | -0.1 | 0.064 | 0.146 | 2.112 | 124 | 51 |
| | packm | | | 0.163 | -0.242 | 0.12 | 0.163 | 0.218 | 0.991 | 170 | 125 |
| | SixMonthPrecip | | | 0.107 | -0.3 | 0.021 | 0.107 | 0.18 | 1.258 | 158 | 61 |
| | Slope | | | -0.118 | -0.182 | -0.156 | -0.118 | -0.096 | 0.009 | 32 | 13 |
| | SoilpH | -0.157 | 0.085 | 0.006 | -0.94 | -0.079 | 0.006 | 0.094 | 1.285 | 122 | 26 |
| | Ydays | | | 0.002 | -0.04 | -0.004 | 0.002 | 0.007 | 0.022 | 223 | 63 |
| | Year | -0.035 | 0.038 | 0.012 | -0.01 | 0.001 | 0.012 | 0.02 | 0.028 | 3 | 0 |

| Species | Variable | Original model coefficient | Original model p | Median | Q 0% | Q 25% | Q 50% | Q 75% | Q 100% | Model Counts | P < 0.05 counts |
|---|---|---|---|---|---|---|---|---|---|---|---|
| *Eriogonum heracleoides* | AnnMinTemp | | | -0.017 | -0.065 | -0.053 | -0.017 | 0.062 | 0.101 | 7 | 4 |
| | Aspect | | | -0.085 | -0.145 | -0.104 | -0.085 | -0.059 | 0.137 | 55 | 41 |
| | AWC | | | -0.089 | -0.211 | -0.127 | -0.089 | -0.05 | 0.095 | 96 | 63 |
| | clay | | | 0.053 | -0.135 | -0.006 | 0.053 | 0.082 | 0.174 | 34 | 17 |
| | DeltFallMax | | | 0.1 | -0.082 | 0.049 | 0.1 | 0.132 | 0.236 | 53 | 30 |
| | DeltFallMin | | | -0.025 | -0.228 | -0.056 | -0.025 | 0.029 | 0.177 | 27 | 9 |
| | DeltPrecip | | | 0.017 | -0.074 | -0.013 | 0.017 | 0.063 | 0.125 | 8 | 1 |
| | DeltSpringMax | 0.073 | 0.016 | 0.086 | -0.109 | 0.055 | 0.086 | 0.115 | 0.334 | 359 | 228 |
| | heatload | | | 0.031 | -0.137 | 0.01 | 0.031 | 0.079 | 0.29 | 44 | 17 |
| | organic | | | 0.066 | -0.135 | 0.033 | 0.066 | 0.159 | 0.661 | 168 | 70 |
| | packm | | | 0.058 | -0.054 | 0.035 | 0.058 | 0.084 | 0.156 | 45 | 25 |
| | SixMonthPrecip | 0.07 | 0.028 | 0.081 | -0.096 | 0.049 | 0.081 | 0.109 | 0.242 | 284 | 154 |
| | Slope | 0.05 | 0.113 | 0.062 | -0.107 | 0.03 | 0.062 | 0.102 | 0.233 | 203 | 90 |
| | SoilpH | | | -0.037 | -0.095 | -0.05 | -0.037 | -0.014 | 0.041 | 36 | 5 |

| Species | Variable | Original model coefficient | Original model p | Median | Q 0% | Q 25% | Q 50% | Q 75% | Q 100% | Model Counts | P < 0.05 counts |
|---|---|---|---|---|---|---|---|---|---|---|---|
| | Ydays | | | 0.001 | -0.005 | -0.001 | 0.001 | 0.002 | 0.008 | 32 | 9 |
| | Year | | | -0.014 | -0.058 | -0.031 | -0.014 | -0.004 | 0.067 | 47 | 14 |
| *Eriogonum umbellatum* | AnnMinTemp | 0.051 | 0.146 | 0.108 | -0.137 | 0.051 | 0.108 | 0.136 | 0.248 | 71 | 37 |
| | Aspect | | | -0.09 | -0.181 | -0.114 | -0.09 | -0.037 | 0.185 | 31 | 18 |
| | AWC | | | 0.082 | -0.17 | 0.019 | 0.082 | 0.149 | 0.26 | 34 | 18 |
| | clay | 0.03 | 0.251 | 0.084 | -0.187 | 0.032 | 0.084 | 0.132 | 0.289 | 101 | 41 |
| | DeltFallMax | | | -0.115 | -0.259 | -0.161 | -0.115 | -0.096 | 0.179 | 29 | 19 |
| | DeltFallMin | | | -0.069 | -0.288 | -0.107 | -0.069 | -0.005 | 0.194 | 16 | 9 |
| | DeltPrecip | | | 0.001 | -0.197 | -0.071 | 0.001 | 0.048 | 0.125 | 39 | 8 |
| | DeltSpringMax | 0.009 | 0.806 | 0.115 | -0.249 | 0.063 | 0.115 | 0.151 | 0.627 | 180 | 88 |
| | heatload | 0.015 | 0.576 | 0.087 | -0.155 | 0.041 | 0.087 | 0.122 | 0.289 | 149 | 61 |
| | organic | | | 0.049 | -0.106 | 0.004 | 0.049 | 0.087 | 0.271 | 16 | 3 |
| | packm | | | 0.109 | -0.17 | 0.068 | 0.109 | 0.133 | 0.217 | 59 | 31 |
| | SixMonthPrecip | | | 0.075 | -0.124 | 0.008 | 0.075 | 0.125 | 0.213 | 19 | 7 |

| Species | Variable | Original model coefficient | Original model p | Median | Q 0% | Q 25% | Q 50% | Q 75% | Q 100% | Model Counts | P < 0.05 counts |
|---|---|---|---|---|---|---|---|---|---|---|---|
| | Slope | | | 0.07 | -0.138 | -0.001 | 0.07 | 0.128 | 0.237 | 39 | 16 |
| | SoilpH | -0.048 | 0.114 | -0.128 | -0.268 | -0.166 | -0.128 | -0.08 | 0.143 | 112 | 65 |
| | Ydays | 0.002 | 0.166 | 0.004 | -0.012 | 0.002 | 0.004 | 0.006 | 0.013 | 206 | 111 |
| | Year | | | -0.006 | -0.048 | -0.023 | -0.006 | 0.003 | 0.042 | 33 | 5 |
| *Eriophyllum lanatum* | AnnMinTemp | | | 0.054 | -0.073 | 0.002 | 0.054 | 0.136 | 0.351 | 16 | 6 |
| | Aspect | | | 0.009 | -0.137 | -0.056 | 0.009 | 0.061 | 0.174 | 12 | 3 |
| | AWC | 0.076 | 0.019 | 0.129 | -0.267 | 0.064 | 0.129 | 0.187 | 0.313 | 268 | 181 |
| | clay | | | 0.071 | -0.246 | 0.026 | 0.071 | 0.119 | 0.261 | 46 | 19 |
| | DeltFallMax | | | 0.101 | -0.047 | 0.07 | 0.101 | 0.123 | 0.318 | 24 | 12 |
| | DeltFallMin | | | 0.045 | -0.186 | 0.001 | 0.045 | 0.097 | 0.221 | 88 | 29 |
| | DeltPrecip | | | 0.104 | -0.038 | 0.064 | 0.104 | 0.149 | 0.379 | 39 | 20 |
| | DeltSpringMax | | | 0.041 | -0.172 | -0.029 | 0.041 | 0.062 | 0.252 | 36 | 12 |
| | heatload | | | -0.093 | -0.219 | -0.145 | -0.093 | -0.061 | 0.171 | 45 | 30 |
| | organic | | | -0.043 | -0.361 | -0.13 | -0.043 | 0.051 | 0.214 | 45 | 20 |

| Species | Variable | Original model coefficient | Original model p | Median | Q 0% | Q 25% | Q 50% | Q 75% | Q 100% | Model Counts | P < 0.05 counts |
|---|---|---|---|---|---|---|---|---|---|---|---|
| | packm | | | -0.071 | -0.183 | -0.109 | -0.071 | -0.036 | 0.156 | 45 | 19 |
| | SixMonthPrecip | | | 0.008 | -0.211 | -0.064 | 0.008 | 0.043 | 0.084 | 9 | 1 |
| | Slope | | | 0.019 | -0.166 | -0.098 | 0.019 | 0.067 | 0.252 | 10 | 5 |
| | SoilpH | 0.085 | 0.087 | 0.073 | -0.363 | 0.007 | 0.073 | 0.124 | 0.397 | 304 | 105 |
| | Ydays | -0.003 | 0.137 | -0.005 | -0.022 | -0.008 | -0.005 | -0.003 | 0.01 | 400 | 206 |
| | Year | | | -0.014 | -0.074 | -0.041 | -0.014 | 0.017 | 0.055 | 18 | 9 |
| *Fallugia paradoxa* | AnnMinTemp | -0.096 | 0.034 | -0.096 | -0.262 | -0.14 | -0.096 | -0.034 | 0.128 | 60 | 33 |
| | Aspect | 0.016 | 0.539 | -0.009 | -0.233 | -0.064 | -0.009 | 0.031 | 0.183 | 95 | 30 |
| | AWC | -0.059 | 0.167 | -0.06 | -0.405 | -0.133 | -0.06 | 0.005 | 0.304 | 231 | 93 |
| | clay | -0.048 | 0.173 | -0.108 | -0.305 | -0.157 | -0.108 | -0.054 | 0.392 | 208 | 133 |
| | DeltFallMax | | | 0.052 | -0.11 | -0.001 | 0.052 | 0.06 | 0.22 | 10 | 4 |
| | DeltFallMin | | | -0.087 | -0.205 | -0.108 | -0.087 | -0.064 | 0.114 | 39 | 24 |
| | DeltPrecip | | | -0.043 | -0.148 | -0.091 | -0.043 | 0.005 | 0.167 | 28 | 10 |
| | DeltSpringMax | | | -0.108 | -0.361 | -0.142 | -0.108 | -0.049 | 0.278 | 131 | 82 |
| | heatload | | | -0.025 | -0.432 | -0.136 | -0.025 | 0.036 | 0.263 | 24 | 9 |

| Species | Variable | Original model coefficient | Original model p | Median | Q 0% | Q 25% | Q 50% | Q 75% | Q 100% | Model Counts | P < 0.05 counts |
|---|---|---|---|---|---|---|---|---|---|---|---|
| | organic | -0.059 | 0.052 | -0.036 | -0.511 | -0.148 | -0.036 | 0.004 | 0.607 | 98 | 35 |
| | packm | | | -0.088 | -0.339 | -0.135 | -0.088 | -0.027 | 0.473 | 177 | 109 |
| | SixMonthPrecip | | | -0.057 | -0.199 | -0.118 | -0.057 | 0.009 | 0.259 | 19 | 11 |
| | Slope | | | -0.029 | -0.074 | -0.044 | -0.029 | 0.051 | 0.178 | 7 | 1 |
| | SoilpH | -0.038 | 0.221 | -0.001 | -0.395 | -0.05 | -0.001 | 0.051 | 0.485 | 199 | 48 |
| | Ydays | -0.001 | 0.099 | -0.002 | -0.007 | -0.003 | -0.002 | -0.001 | 0.004 | 277 | 136 |
| | Year | | | -0.034 | -0.055 | -0.045 | -0.034 | -0.025 | -0.017 | 3 | 1 |
| *Festuca idahoensis* | AnnMinTemp | -0.014 | 0.708 | 0.069 | -0.423 | 0.016 | 0.069 | 0.106 | 0.189 | 166 | 75 |
| | Aspect | 0.019 | 0.513 | -0.051 | -0.083 | -0.067 | -0.051 | -0.035 | -0.019 | 2 | 1 |
| | AWC | 0.09 | 0.026 | 0.1 | -0.006 | 0.069 | 0.1 | 0.131 | 0.207 | 96 | 69 |
| | clay | 0.05 | 0.119 | 0.067 | -0.083 | 0.041 | 0.067 | 0.092 | 0.241 | 212 | 119 |
| | DeltFallMax | | | -0.068 | -0.176 | -0.088 | -0.068 | -0.049 | 0.11 | 120 | 74 |
| | DeltFallMin | | | 0.073 | -0.012 | 0.056 | 0.073 | 0.105 | 0.15 | 30 | 22 |
| | DeltPrecip | | | -0.009 | -0.055 | -0.04 | -0.009 | 0.049 | 0.092 | 26 | 6 |
| | DeltSpringMax | | | -0.065 | -0.208 | -0.091 | -0.065 | -0.032 | 0.04 | 58 | 23 |

| Species | Variable | Original model coefficient | Original model p | Median | Q 0% | Q 25% | Q 50% | Q 75% | Q 100% | Model Counts | P < 0.05 counts |
|---|---|---|---|---|---|---|---|---|---|---|---|
| | heatload | | | 0.041 | -0.011 | 0.024 | 0.041 | 0.068 | 0.145 | 25 | 7 |
| | organic | -0.116 | 0.010 | -0.008 | -0.134 | -0.059 | -0.008 | 0.025 | 0.683 | 37 | 5 |
| | packm | | | 0.004 | -0.114 | -0.077 | 0.004 | 0.055 | 0.166 | 24 | 14 |
| | SixMonthPrecip | | | 0.073 | -0.155 | 0.034 | 0.073 | 0.091 | 0.27 | 141 | 76 |
| | Slope | | | 0.004 | -0.111 | -0.026 | 0.004 | 0.037 | 0.076 | 11 | 4 |
| | SoilpH | -0.103 | 0.009 | -0.081 | -0.212 | -0.103 | -0.081 | -0.059 | 0.084 | 46 | 26 |
| | Ydays | 0 | 0.872 | 0.005 | -0.004 | 0.002 | 0.005 | 0.007 | 0.012 | 50 | 28 |
| | Year | | | -0.015 | -0.047 | -0.018 | -0.015 | -0.012 | -0.003 | 13 | 4 |
| *Grayia spinosa* | AnnMinTemp | | | 0.059 | 0.023 | 0.036 | 0.059 | 0.087 | 0.293 | 8 | 5 |
| | Aspect | | | 0.041 | -0.117 | 0.004 | 0.041 | 0.063 | 0.105 | 39 | 15 |
| | AWC | | | -0.059 | -0.172 | -0.099 | -0.059 | -0.001 | 0.213 | 31 | 14 |
| | clay | | | 0.027 | -0.098 | -0.032 | 0.027 | 0.051 | 0.104 | 32 | 10 |
| | DeltFallMax | -0.043 | 0.050 | -0.063 | -0.178 | -0.097 | -0.063 | -0.034 | 0.041 | 158 | 83 |
| | DeltFallMin | | | 0.057 | -0.082 | 0.017 | 0.057 | 0.075 | 0.132 | 19 | 7 |
| | DeltPrecip | | | -0.051 | -0.121 | -0.08 | -0.051 | -0.024 | 0.145 | 62 | 24 |

| Species | Variable | Original model coefficient | Original model p | Median | Q 0% | Q 25% | Q 50% | Q 75% | Q 100% | Model Counts | P < 0.05 counts |
|---|---|---|---|---|---|---|---|---|---|---|---|
| | DeltSpringMax | | | -0.034 | -0.117 | -0.074 | -0.034 | -0.022 | 0.064 | 29 | 13 |
| | heatload | | | -0.063 | -0.224 | -0.086 | -0.063 | -0.043 | 0.065 | 119 | 62 |
| | organic | | | 0.046 | -0.131 | 0.021 | 0.046 | 0.069 | 0.279 | 56 | 16 |
| | packm | | | -0.018 | -0.43 | -0.075 | -0.018 | 0.002 | 0.225 | 74 | 20 |
| | SixMonthPrecip | 0.037 | 0.075 | 0.07 | -0.01 | 0.053 | 0.07 | 0.084 | 0.299 | 145 | 92 |
| | Slope | | | 0.025 | -0.122 | -0.01 | 0.025 | 0.057 | 0.258 | 46 | 12 |
| | SoilpH | | | 0.066 | -0.057 | 0.035 | 0.066 | 0.088 | 0.13 | 34 | 20 |
| | Ydays | | | 0.002 | -0.003 | 0.001 | 0.002 | 0.003 | 0.008 | 27 | 10 |
| | Year | | | 0.004 | -0.048 | -0.012 | 0.004 | 0.019 | 0.042 | 37 | 10 |
| *Helianthus annuus* | AnnMinTemp | | | -0.014 | -0.092 | -0.049 | -0.014 | 0.01 | 0.087 | 33 | 13 |
| | Aspect | | | 0.033 | -0.021 | 0.023 | 0.033 | 0.055 | 0.093 | 40 | 16 |
| | AWC | | | 0.017 | -0.063 | -0.023 | 0.017 | 0.019 | 0.02 | 3 | 1 |
| | clay | | | 0.042 | -0.025 | 0.014 | 0.042 | 0.056 | 0.152 | 102 | 39 |
| | DeltFallMax | | | 0.036 | -0.049 | -0.008 | 0.036 | 0.051 | 0.07 | 13 | 5 |
| | DeltFallMin | | | 0.029 | -0.034 | 0.017 | 0.029 | 0.044 | 0.082 | 23 | 7 |

| Species | Variable | Original model coefficient | Original model p | Median | Q 0% | Q 25% | Q 50% | Q 75% | Q 100% | Model Counts | P < 0.05 counts |
|---|---|---|---|---|---|---|---|---|---|---|---|
| | DeltPrecip | | | 0.051 | -0.042 | 0.032 | 0.051 | 0.062 | 0.082 | 9 | 6 |
| | DeltSpringMax | | | 0.026 | -0.031 | 0.013 | 0.026 | 0.038 | 0.077 | 34 | 8 |
| | heatload | -0.068 | <0.001 | -0.068 | -0.147 | -0.08 | -0.068 | -0.046 | 0.123 | 132 | 95 |
| | organic | 0.031 | 0.078 | 0.039 | -0.337 | 0.017 | 0.039 | 0.107 | 0.326 | 181 | 83 |
| | packm | | | 0.009 | -0.091 | -0.014 | 0.009 | 0.026 | 0.102 | 46 | 10 |
| | SixMonthPrecip | 0.05 | 0.005 | 0.043 | -0.062 | 0.027 | 0.043 | 0.065 | 0.188 | 228 | 112 |
| | Slope | | | 0.033 | -0.062 | -0.002 | 0.033 | 0.058 | 0.135 | 69 | 28 |
| | SoilpH | -0.037 | 0.047 | -0.05 | -0.161 | -0.066 | -0.05 | -0.031 | 0.407 | 306 | 156 |
| | Ydays | | | -0.001 | -0.002 | -0.001 | -0.001 | 0 | 0.004 | 24 | 3 |
| | Year | | | -0.01 | -0.015 | -0.011 | -0.01 | -0.006 | -0.001 | 7 | 0 |
| *Heliomeris multiflora* | AnnMinTemp | | | 0.096 | -0.212 | 0.028 | 0.096 | 0.162 | 0.644 | 169 | 60 |
| | Aspect | | | 0.17 | -0.194 | 0.069 | 0.17 | 0.266 | 0.454 | 64 | 44 |
| | AWC | | | -0.115 | -0.364 | -0.164 | -0.115 | -0.043 | 0.359 | 95 | 51 |
| | clay | | | -0.124 | -0.387 | -0.196 | -0.124 | -0.058 | 0.265 | 128 | 72 |

| Species | Variable | Original model coefficient | Original model p | Median | Q 0% | Q 25% | Q 50% | Q 75% | Q 100% | Model Counts | P < 0.05 counts |
|---|---|---|---|---|---|---|---|---|---|---|---|
| | DeltFallMax | | | 0.008 | -0.232 | -0.037 | 0.008 | 0.033 | 0.154 | 33 | 8 |
| | DeltFallMin | | | -0.041 | -0.188 | -0.078 | -0.041 | 0.008 | 0.234 | 29 | 4 |
| | DeltPrecip | | | -0.023 | -0.3 | -0.076 | -0.023 | 0.063 | 0.211 | 79 | 28 |
| | DeltSpringMax | | | -0.077 | -0.209 | -0.115 | -0.077 | -0.025 | 0.062 | 20 | 6 |
| | heatload | | | -0.056 | -0.146 | -0.107 | -0.056 | -0.002 | 0.04 | 7 | 1 |
| | organic | | | -0.005 | -0.854 | -0.142 | -0.005 | 0.077 | 0.412 | 78 | 17 |
| | packm | | | -0.041 | -0.208 | -0.095 | -0.041 | 0.01 | 0.382 | 42 | 16 |
| | SixMonthPrecip | 0.058 | 0.347 | -0.042 | -1.317 | -0.215 | -0.042 | 0.086 | 0.496 | 161 | 53 |
| | Slope | | | 0.119 | -0.129 | 0.086 | 0.119 | 0.131 | 0.239 | 15 | 13 |
| | SoilpH | | | -0.022 | -0.373 | -0.08 | -0.022 | 0.054 | 0.267 | 38 | 15 |
| | Ydays | -0.005 | <0.001 | -0.003 | -0.013 | -0.004 | -0.003 | -0.002 | 0.015 | 262 | 163 |
| | Year | | | -0.018 | -0.077 | -0.026 | -0.018 | -0.007 | 0.055 | 24 | 5 |
| *Hesperostipa comata* | AnnMinTemp | 0.04 | 0.126 | 0.111 | -0.094 | 0.055 | 0.111 | 0.148 | 0.205 | 54 | 33 |
| | Aspect | | | 0.082 | -0.176 | -0.093 | 0.082 | 0.117 | 0.161 | 21 | 14 |

| Species | Variable | Original model coefficient | Original model p | Median | Q 0% | Q 25% | Q 50% | Q 75% | Q 100% | Model Counts | P < 0.05 counts |
|---|---|---|---|---|---|---|---|---|---|---|---|
| | AWC | | | 0.109 | -0.151 | 0.051 | 0.109 | 0.143 | 0.224 | 41 | 22 |
| | clay | | | 0.109 | -0.124 | 0.001 | 0.109 | 0.135 | 0.266 | 24 | 15 |
| | DeltFallMax | | | -0.122 | -0.23 | -0.145 | -0.122 | -0.068 | 0.166 | 30 | 21 |
| | DeltFallMin | | | -0.098 | -0.213 | -0.132 | -0.098 | -0.073 | 0.154 | 32 | 17 |
| | DeltPrecip | 0.081 | 0.004 | 0.116 | -0.054 | 0.078 | 0.116 | 0.147 | 0.266 | 71 | 43 |
| | DeltSpringMax | | | 0.08 | -0.132 | 0.054 | 0.08 | 0.119 | 0.263 | 51 | 18 |
| | heatload | 0.032 | 0.128 | 0.094 | -0.202 | 0.054 | 0.094 | 0.16 | 0.337 | 101 | 45 |
| | organic | | | 0.065 | -0.277 | 0.007 | 0.065 | 0.134 | 0.273 | 41 | 17 |
| | packm | | | -0.036 | -0.211 | -0.086 | -0.036 | 0.054 | 0.207 | 35 | 9 |
| | SixMonthPrecip | | | 0.098 | -0.121 | 0.053 | 0.098 | 0.136 | 0.224 | 44 | 20 |
| | Slope | | | 0.061 | -0.414 | 0.003 | 0.061 | 0.122 | 0.235 | 30 | 12 |
| | SoilpH | | | -0.061 | -0.222 | -0.108 | -0.061 | -0.025 | 0.252 | 35 | 17 |
| | Ydays | | | 0.002 | -0.012 | -0.007 | 0.002 | 0.007 | 0.014 | 25 | 17 |
| | Year | 0.024 | 0.008 | 0.015 | -0.065 | -0.019 | 0.015 | 0.029 | 0.047 | 50 | 14 |
| *Heterotheca villosa* | AnnMinTemp | | | -0.042 | -0.206 | -0.11 | -0.042 | 0.026 | 0.193 | 45 | 15 |

| Species | Variable | Original model coefficient | Original model p | Median | Q 0% | Q 25% | Q 50% | Q 75% | Q 100% | Model Counts | P < 0.05 counts |
|---|---|---|---|---|---|---|---|---|---|---|---|
| | Aspect | | | 0.01 | -0.275 | -0.059 | 0.01 | 0.075 | 0.1 | 20 | 11 |
| | AWC | | | 0.063 | -0.561 | 0.022 | 0.063 | 0.11 | 0.276 | 37 | 17 |
| | clay | | | -0.007 | -0.109 | -0.043 | -0.007 | 0.089 | 0.388 | 26 | 7 |
| | DeltFallMax | -0.065 | 0.097 | -0.094 | -0.198 | -0.13 | -0.094 | -0.063 | 0.015 | 64 | 28 |
| | DeltFallMin | | | -0.005 | -0.187 | -0.065 | -0.005 | 0.064 | 0.139 | 14 | 2 |
| | DeltPrecip | | | -0.087 | -0.236 | -0.142 | -0.087 | -0.037 | 0.125 | 39 | 15 |
| | DeltSpringMax | -0.056 | 0.134 | -0.091 | -0.385 | -0.152 | -0.091 | -0.042 | 0.224 | 79 | 33 |
| | heatload | | | -0.153 | -0.471 | -0.207 | -0.153 | -0.076 | 0.221 | 34 | 19 |
| | organic | | | 0.047 | -1.158 | 0.014 | 0.047 | 0.093 | 1.396 | 77 | 25 |
| | packm | | | -0.028 | -0.155 | -0.066 | -0.028 | 0.053 | 0.265 | 30 | 9 |
| | SixMonthPrecip | 0.149 | <0.001 | 0.148 | -0.109 | 0.086 | 0.148 | 0.218 | 0.507 | 287 | 186 |
| | Slope | | | -0.204 | -0.714 | -0.268 | -0.204 | -0.142 | 0.174 | 28 | 24 |
| | SoilpH | | | -0.069 | -0.186 | -0.107 | -0.069 | -0.015 | 0.275 | 45 | 17 |
| | Ydays | 0.001 | 0.365 | 0.003 | -0.005 | 0.002 | 0.003 | 0.005 | 0.017 | 61 | 29 |
| | Year | | | -0.028 | -0.066 | -0.046 | -0.028 | -0.007 | 0.08 | 60 | 20 |

| Species | Variable | Original model coefficient | Original model p | Median | Q 0% | Q 25% | Q 50% | Q 75% | Q 100% | Model Counts | P < 0.05 counts |
|---|---|---|---|---|---|---|---|---|---|---|---|
| *Koeleria macrantha* | AnnMinTemp | | | 0.068 | -0.183 | 0.013 | 0.068 | 0.112 | 0.287 | 26 | 11 |
| | Aspect | | | -0.041 | -0.116 | -0.064 | -0.041 | 0.06 | 0.184 | 14 | 5 |
| | AWC | | | -0.108 | -0.259 | -0.199 | -0.108 | -0.047 | 0.03 | 10 | 5 |
| | clay | | | 0.073 | -0.027 | 0.035 | 0.073 | 0.114 | 0.239 | 87 | 25 |
| | DeltFallMax | | | 0.077 | -0.109 | 0.053 | 0.077 | 0.091 | 0.42 | 8 | 3 |
| | DeltFallMin | | | 0.09 | -0.128 | 0.059 | 0.09 | 0.121 | 0.237 | 31 | 9 |
| | DeltPrecip | -0.156 | 0.004 | -0.139 | -0.453 | -0.17 | -0.139 | -0.109 | 0.183 | 428 | 286 |
| | DeltSpringMax | 0.02 | 0.717 | -0.029 | -0.428 | -0.076 | -0.029 | 0.016 | 0.227 | 194 | 32 |
| | heatload | | | 0.09 | -0.161 | 0.026 | 0.09 | 0.124 | 0.181 | 18 | 7 |
| | organic | | | 0.006 | -0.678 | -0.052 | 0.006 | 0.089 | 0.533 | 57 | 11 |
| | packm | | | -0.075 | -0.75 | -0.234 | -0.075 | -0.028 | 0.115 | 67 | 30 |
| | SixMonthPrecip | | | 0.01 | -0.163 | -0.023 | 0.01 | 0.127 | 0.343 | 39 | 4 |
| | Slope | | | 0.066 | -0.55 | 0.04 | 0.066 | 0.088 | 0.217 | 23 | 11 |
| | SoilpH | | | -0.06 | -0.3 | -0.1 | -0.06 | -0.028 | 0.151 | 16 | 5 |
| | Ydays | | | -0.001 | -0.007 | -0.002 | -0.001 | 0 | 0.005 | 29 | 7 |

| Species | Variable | Original model coefficient | Original model p | Median | Q 0% | Q 25% | Q 50% | Q 75% | Q 100% | Model Counts | P < 0.05 counts |
|---|---|---|---|---|---|---|---|---|---|---|---|
| | Year | | | 0.012 | -0.095 | -0.002 | 0.012 | 0.034 | 0.092 | 63 | 15 |
| *Krascheninnikovia lanata* | AnnMinTemp | 0.103 | 0.005 | 0.133 | -0.301 | 0.081 | 0.133 | 0.179 | 0.369 | 315 | 194 |
| | Aspect | | | 0.07 | -0.168 | -0.013 | 0.07 | 0.114 | 0.229 | 45 | 19 |
| | AWC | | | -0.056 | -0.192 | -0.089 | -0.056 | -0.004 | 0.168 | 67 | 16 |
| | clay | | | 0.049 | -0.142 | 0.002 | 0.049 | 0.091 | 0.252 | 47 | 13 |
| | DeltFallMax | | | -0.011 | -0.137 | -0.061 | -0.011 | 0.062 | 0.159 | 12 | 5 |
| | DeltFallMin | | | 0.091 | -0.098 | 0.042 | 0.091 | 0.132 | 0.214 | 60 | 28 |
| | DeltPrecip | | | -0.108 | -0.271 | -0.13 | -0.108 | -0.035 | 0.171 | 28 | 21 |
| | DeltSpringMax | | | -0.094 | -0.161 | -0.118 | -0.094 | -0.02 | 0.234 | 12 | 7 |
| | heatload | | | -0.037 | -0.306 | -0.123 | -0.037 | 0.044 | 0.319 | 47 | 21 |
| | organic | | | 0.089 | -0.118 | 0.029 | 0.089 | 0.145 | 0.262 | 30 | 14 |
| | packm | 0.007 | 0.833 | -0.049 | -0.374 | -0.115 | -0.049 | -0.004 | 0.189 | 117 | 36 |
| | SixMonthPrecip | | | 0.108 | -0.079 | 0.055 | 0.108 | 0.159 | 0.309 | 75 | 41 |
| | Slope | 0.058 | 0.028 | 0.109 | -0.149 | 0.063 | 0.109 | 0.148 | 0.233 | 132 | 81 |

| Species | Variable | Original model coefficient | Original model p | Median | Q 0% | Q 25% | Q 50% | Q 75% | Q 100% | Model Counts | P < 0.05 counts |
|---|---|---|---|---|---|---|---|---|---|---|---|
| | SoilpH | -0.085 | 0.002 | -0.106 | -0.338 | -0.15 | -0.106 | -0.059 | 0.117 | 116 | 69 |
| | Ydays | 0 | 0.519 | -0.001 | -0.004 | -0.002 | -0.001 | 0 | 0.004 | 219 | 69 |
| | Year | | | 0.019 | -0.055 | 0.014 | 0.019 | 0.036 | 0.058 | 33 | 15 |
| *Larrea tridentata* | AnnMinTemp | -0.091 | 0.142 | -0.138 | -0.605 | -0.184 | -0.138 | -0.089 | 0.093 | 134 | 69 |
| | Aspect | | | -0.074 | -0.176 | -0.097 | -0.074 | 0.104 | 0.376 | 27 | 15 |
| | AWC | | | -0.15 | -0.457 | -0.339 | -0.15 | -0.054 | 0.246 | 40 | 15 |
| | clay | | | 0.065 | -0.363 | 0.001 | 0.065 | 0.131 | 0.628 | 44 | 14 |
| | DeltFallMax | | | 0.082 | -0.25 | -0.062 | 0.082 | 0.228 | 0.412 | 32 | 10 |
| | DeltFallMin | | | 0.055 | -0.153 | -0.074 | 0.055 | 0.094 | 0.145 | 21 | 9 |
| | DeltPrecip | | | 0.056 | -0.5 | 0.001 | 0.056 | 0.096 | 0.272 | 66 | 23 |
| | DeltSpringMax | | | 0.105 | -0.438 | -0.095 | 0.105 | 0.298 | 1.218 | 44 | 15 |
| | heatload | | | -0.018 | -0.321 | -0.159 | -0.018 | 0.056 | 0.284 | 28 | 9 |
| | organic | | | 0.081 | -1.008 | 0.017 | 0.081 | 0.155 | 0.323 | 61 | 20 |
| | packm | 0.084 | 0.190 | 0.098 | -1.278 | 0.062 | 0.098 | 0.171 | 1.279 | 145 | 61 |
| | SixMonthPrecip | 0.059 | 0.336 | 0.114 | -0.25 | 0.089 | 0.114 | 0.151 | 0.415 | 105 | 63 |

| Species | Variable | Original model coefficient | Original model p | Median | Q 0% | Q 25% | Q 50% | Q 75% | Q 100% | Model Counts | P < 0.05 counts |
|---|---|---|---|---|---|---|---|---|---|---|---|
| | Slope | | | -0.049 | -0.325 | -0.146 | -0.049 | 0.061 | 0.401 | 26 | 7 |
| | SoilpH | | | 0.137 | -0.253 | -0.061 | 0.137 | 0.546 | 1.028 | 42 | 26 |
| | Ydays | | | 0.001 | -0.005 | 0 | 0.001 | 0.003 | 0.007 | 36 | 12 |
| | Year | | | 0.045 | -0.036 | 0.016 | 0.045 | 0.063 | 0.447 | 48 | 26 |
| *Leymus cinereus* | AnnMinTemp | 0.03 | 0.047 | 0.07 | -0.056 | 0.051 | 0.07 | 0.09 | 0.176 | 85 | 50 |
| | Aspect | | | -0.005 | -0.154 | -0.044 | -0.005 | 0.058 | 0.091 | 22 | 12 |
| | AWC | | | -0.059 | -0.125 | -0.069 | -0.059 | -0.023 | 0.131 | 41 | 28 |
| | clay | | | 0.048 | -0.09 | 0.025 | 0.048 | 0.065 | 0.091 | 27 | 11 |
| | DeltFallMax | | | 0.015 | -0.11 | -0.03 | 0.015 | 0.045 | 0.11 | 28 | 9 |
| | DeltFallMin | | | 0.03 | -0.127 | 0.004 | 0.03 | 0.056 | 0.097 | 35 | 14 |
| | DeltPrecip | | | 0.076 | -0.083 | -0.019 | 0.076 | 0.127 | 0.186 | 49 | 28 |
| | DeltSpringMax | -0.024 | 0.112 | -0.062 | -0.25 | -0.086 | -0.062 | -0.043 | 0.037 | 86 | 42 |
| | heatload | | | -0.065 | -0.287 | -0.097 | -0.065 | -0.034 | 0.087 | 35 | 19 |
| | organic | | | -0.036 | -0.296 | -0.074 | -0.036 | -0.009 | 0.096 | 35 | 5 |
| | packm | | | -0.047 | -0.135 | -0.06 | -0.047 | -0.018 | 0.059 | 34 | 17 |

| Species | Variable | Original model coefficient | Original model p | Median | Q 0% | Q 25% | Q 50% | Q 75% | Q 100% | Model Counts | P < 0.05 counts |
|---|---|---|---|---|---|---|---|---|---|---|---|
| | SixMonthPrecip | | | -0.051 | -0.119 | -0.084 | -0.051 | -0.018 | 0.065 | 34 | 14 |
| | Slope | | | 0.006 | -0.134 | -0.063 | 0.006 | 0.068 | 0.104 | 28 | 15 |
| | SoilpH | -0.023 | 0.096 | -0.042 | -0.175 | -0.059 | -0.042 | -0.009 | 0.124 | 35 | 15 |
| | Ydays | | | 0.002 | -0.002 | 0 | 0.002 | 0.003 | 0.005 | 22 | 13 |
| | Year | -0.005 | 0.166 | 0 | -0.051 | -0.013 | 0 | 0.007 | 0.028 | 35 | 14 |
| *Linum lewisii* | AnnMinTemp | 0.112 | 0.049 | 0.114 | -0.22 | 0.079 | 0.114 | 0.145 | 0.336 | 305 | 163 |
| | Aspect | | | 0.036 | -0.207 | -0.022 | 0.036 | 0.087 | 0.177 | 56 | 24 |
| | AWC | -0.127 | 0.029 | -0.152 | -0.386 | -0.178 | -0.152 | -0.117 | 0.187 | 265 | 188 |
| | clay | | | 0.031 | -0.411 | -0.04 | 0.031 | 0.09 | 0.157 | 41 | 7 |
| | DeltFallMax | | | -0.084 | -0.157 | -0.135 | -0.084 | -0.078 | 0.034 | 5 | 1 |
| | DeltFallMin | | | -0.077 | -0.384 | -0.132 | -0.077 | -0.044 | 0.168 | 37 | 11 |
| | DeltPrecip | | | 0.142 | -0.014 | 0.081 | 0.142 | 0.215 | 0.349 | 8 | 5 |
| | DeltSpringMax | | | 0.066 | -0.277 | 0.021 | 0.066 | 0.111 | 0.214 | 68 | 19 |
| | heatload | 0.02 | 0.709 | 0.044 | -0.137 | 0.02 | 0.044 | 0.217 | 1.374 | 195 | 57 |
| | organic | | | 0.004 | -0.408 | -0.048 | 0.004 | 0.021 | 0.19 | 14 | 5 |

| Species | Variable | Original model coefficient | Original model p | Median | Q 0% | Q 25% | Q 50% | Q 75% | Q 100% | Model Counts | P < 0.05 counts |
|---|---|---|---|---|---|---|---|---|---|---|---|
| | packm | 0.026 | 0.654 | -0.041 | -0.639 | -0.098 | -0.041 | -0.005 | 0.114 | 135 | 21 |
| | SixMonthPrecip | | | -0.061 | -0.226 | -0.086 | -0.061 | 0.006 | 0.188 | 24 | 9 |
| | Slope | | | 0.059 | -0.128 | -0.116 | 0.059 | 0.346 | 1.118 | 6 | 3 |
| | SoilpH | | | -0.102 | -0.283 | -0.185 | -0.102 | -0.035 | 0.118 | 46 | 23 |
| | Ydays | | | 0 | -0.008 | -0.002 | 0 | 0.003 | 0.02 | 50 | 16 |
| | Year | | | -0.032 | -0.091 | -0.051 | -0.032 | -0.019 | 0.045 | 39 | 15 |
| *Lomatium triternatum* | AnnMinTemp | | | -0.086 | -0.176 | -0.117 | -0.086 | -0.015 | 0.083 | 19 | 8 |
| | Aspect | | | 0.105 | -0.045 | 0.063 | 0.105 | 0.143 | 0.328 | 72 | 45 |
| | AWC | | | -0.064 | -0.205 | -0.104 | -0.064 | -0.02 | 0.059 | 24 | 9 |
| | clay | | | 0.068 | -0.118 | 0.019 | 0.068 | 0.123 | 0.567 | 48 | 21 |
| | DeltFallMax | 0.116 | 0.010 | 0.096 | -0.262 | 0.05 | 0.096 | 0.134 | 0.3 | 259 | 133 |
| | DeltFallMin | 0.072 | 0.065 | 0.111 | -0.226 | 0.068 | 0.111 | 0.182 | 0.636 | 198 | 117 |
| | DeltPrecip | | | 0.032 | -0.231 | -0.021 | 0.032 | 0.089 | 0.215 | 65 | 13 |
| | DeltSpringMax | | | 0.027 | -0.129 | -0.004 | 0.027 | 0.104 | 0.217 | 50 | 14 |

| Species | Variable | Original model coefficient | Original model p | Median | Q 0% | Q 25% | Q 50% | Q 75% | Q 100% | Model Counts | P < 0.05 counts |
|---|---|---|---|---|---|---|---|---|---|---|---|
| | heatload | 0 | 0.999 | 0.101 | -0.317 | 0.03 | 0.101 | 0.153 | 0.315 | 150 | 63 |
| | organic | | | 0.056 | -0.05 | 0.03 | 0.056 | 0.083 | 0.259 | 65 | 13 |
| | packm | | | -0.063 | -0.201 | -0.098 | -0.063 | -0.035 | 0.113 | 19 | 7 |
| | SixMonthPrecip | | | -0.102 | -0.256 | -0.128 | -0.102 | -0.081 | 0.249 | 65 | 43 |
| | Slope | -0.129 | 0.109 | -0.128 | -0.561 | -0.184 | -0.128 | -0.058 | 0.482 | 292 | 152 |
| | SoilpH | | | -0.045 | -0.172 | -0.091 | -0.045 | -0.006 | 0.166 | 42 | 11 |
| | Ydays | | | 0.004 | -0.008 | 0.002 | 0.004 | 0.006 | 0.015 | 200 | 93 |
| | Year | | | -0.033 | -0.182 | -0.067 | -0.033 | -0.014 | 0.104 | 79 | 34 |
| *Machaeranthera canescens* | AnnMinTemp | | | -0.018 | -0.278 | -0.088 | -0.018 | 0.099 | 0.381 | 24 | 7 |
| | Aspect | | | -0.131 | -0.345 | -0.176 | -0.131 | -0.096 | 0.025 | 79 | 48 |
| | AWC | | | 0.097 | -0.144 | -0.018 | 0.097 | 0.14 | 0.293 | 25 | 12 |
| | clay | | | 0.053 | -0.134 | 0.009 | 0.053 | 0.124 | 0.274 | 35 | 12 |
| | DeltFallMax | -0.052 | 0.228 | -0.129 | -0.296 | -0.193 | -0.129 | -0.079 | 0.094 | 68 | 34 |
| | DeltFallMin | | | 0.092 | -0.095 | 0.009 | 0.092 | 0.131 | 0.286 | 58 | 17 |

| Species | Variable | Original model coefficient | Original model p | Median | Q 0% | Q 25% | Q 50% | Q 75% | Q 100% | Model Counts | P < 0.05 counts |
|---|---|---|---|---|---|---|---|---|---|---|---|
| | DeltPrecip | | | 0.105 | -0.185 | -0.026 | 0.105 | 0.146 | 0.264 | 11 | 5 |
| | DeltSpringMax | | | -0.164 | -0.31 | -0.202 | -0.164 | -0.119 | 0.164 | 55 | 38 |
| | heatload | -0.101 | 0.084 | -0.124 | -0.343 | -0.18 | -0.124 | -0.061 | 0.145 | 108 | 40 |
| | organic | | | 0.207 | -0.531 | 0.04 | 0.207 | 0.568 | 1.336 | 88 | 40 |
| | packm | | | -0.058 | -0.384 | -0.122 | -0.058 | -0.006 | 0.468 | 52 | 17 |
| | SixMonthPrecip | 0.11 | 0.023 | 0.129 | -0.184 | 0.084 | 0.129 | 0.185 | 0.333 | 141 | 72 |
| | Slope | 0.004 | 0.936 | 0.142 | -0.082 | 0.07 | 0.142 | 0.211 | 0.423 | 163 | 78 |
| | SoilpH | | | -0.084 | -0.236 | -0.134 | -0.084 | 0.054 | 0.205 | 23 | 10 |
| | Ydays | -0.002 | 0.198 | -0.004 | -0.014 | -0.005 | -0.004 | -0.002 | 0.006 | 113 | 49 |
| | Year | | | -0.018 | -0.103 | -0.029 | -0.018 | 0.004 | 0.056 | 41 | 12 |
| *Machaeranthera pinnatifida* | AnnMinTemp | | | -0.092 | -0.475 | -0.141 | -0.092 | -0.007 | 0.562 | 72 | 34 |
| | Aspect | | | -0.004 | -0.139 | -0.042 | -0.004 | 0.034 | 0.199 | 124 | 23 |
| | AWC | | | 0.104 | -0.014 | 0.017 | 0.104 | 0.111 | 0.165 | 5 | 2 |
| | clay | | | 0.072 | -0.12 | 0.015 | 0.072 | 0.11 | 0.245 | 14 | 7 |

| Species | Variable | Original model coefficient | Original model p | Median | Q 0% | Q 25% | Q 50% | Q 75% | Q 100% | Model Counts | P < 0.05 counts |
|---|---|---|---|---|---|---|---|---|---|---|---|
| | DeltFallMax | | | -0.051 | -0.329 | -0.089 | -0.051 | 0.038 | 0.05 | 7 | 1 |
| | DeltFallMin | | | 0.031 | -0.336 | -0.192 | 0.031 | 0.125 | 0.216 | 19 | 13 |
| | DeltPrecip | | | -0.006 | -0.486 | -0.075 | -0.006 | 0.041 | 0.16 | 46 | 5 |
| | DeltSpringMax | | | -0.082 | -0.445 | -0.136 | -0.082 | -0.008 | 0.27 | 125 | 61 |
| | heatload | -0.201 | <0.001 | -0.16 | -0.424 | -0.208 | -0.16 | -0.115 | 0.098 | 387 | 317 |
| | organic | | | -0.075 | -0.232 | -0.11 | -0.075 | -0.04 | 0.045 | 43 | 19 |
| | packm | | | -0.028 | -0.228 | -0.078 | -0.028 | 0.003 | 0.142 | 100 | 20 |
| | SixMonthPrecip | | | 0.091 | -0.156 | -0.041 | 0.091 | 0.212 | 0.333 | 29 | 14 |
| | Slope | | | 0.046 | -0.419 | -0.029 | 0.046 | 0.115 | 0.319 | 151 | 68 |
| | SoilpH | | | -0.001 | -0.077 | -0.062 | -0.001 | 0.031 | 0.098 | 8 | 3 |
| | Ydays | -0.001 | 0.144 | -0.001 | -0.005 | -0.003 | -0.001 | 0 | 0.006 | 128 | 60 |
| | Year | | | -0.044 | -0.136 | -0.059 | -0.044 | -0.031 | 0.068 | 213 | 161 |
| *Machaeranthera tanacetifolia* | AnnMinTemp | | | -0.091 | -0.196 | -0.123 | -0.091 | -0.049 | 0.129 | 38 | 26 |
| | Aspect | | | 0.105 | 0 | 0.075 | 0.105 | 0.14 | 0.211 | 121 | 98 |

| Species | Variable | Original model coefficient | Original model p | Median | Q 0% | Q 25% | Q 50% | Q 75% | Q 100% | Model Counts | P < 0.05 counts |
|---|---|---|---|---|---|---|---|---|---|---|---|
| | AWC | | | -0.064 | -0.152 | -0.103 | -0.064 | -0.007 | 0.123 | 16 | 12 |
| | clay | | | -0.07 | -0.197 | -0.094 | -0.07 | -0.057 | -0.009 | 37 | 18 |
| | DeltFallMax | | | 0.068 | -0.095 | 0.044 | 0.068 | 0.095 | 0.217 | 26 | 16 |
| | DeltFallMin | | | -0.021 | -0.089 | -0.048 | -0.021 | 0.044 | 0.119 | 12 | 2 |
| | DeltPrecip | | | -0.033 | -0.107 | -0.054 | -0.033 | -0.015 | 0.029 | 7 | 1 |
| | DeltSpringMax | | | -0.052 | -0.218 | -0.086 | -0.052 | -0.019 | 0.11 | 81 | 17 |
| | heatload | -0.029 | 0.396 | -0.04 | -0.158 | -0.078 | -0.04 | 0.009 | 0.211 | 90 | 20 |
| | organic | | | 0.013 | -0.129 | -0.007 | 0.013 | 0.052 | 0.145 | 41 | 7 |
| | packm | 0.067 | 0.057 | 0.09 | -0.057 | 0.061 | 0.09 | 0.122 | 0.301 | 197 | 113 |
| | SixMonthPrecip | | | 0.091 | -0.044 | 0.052 | 0.091 | 0.124 | 0.239 | 75 | 44 |
| | Slope | | | 0.064 | -0.155 | -0.036 | 0.064 | 0.091 | 0.318 | 48 | 25 |
| | SoilpH | | | 0.116 | -0.081 | 0.083 | 0.116 | 0.147 | 0.229 | 110 | 81 |
| | Ydays | | | -0.002 | -0.005 | -0.002 | -0.002 | -0.001 | 0.002 | 107 | 42 |
| | Year | | | 0.036 | 0.014 | 0.019 | 0.036 | 0.042 | 0.054 | 8 | 3 |

| Species | Variable | Original model coefficient | Original model p | Median | Q 0% | Q 25% | Q 50% | Q 75% | Q 100% | Model Counts | P < 0.05 counts |
|---|---|---|---|---|---|---|---|---|---|---|---|
| *Malacothrix glabrata* | AnnMinTemp | | | -0.076 | -0.262 | -0.101 | -0.076 | -0.032 | 0.206 | 108 | 42 |
| | Aspect | | | -0.046 | -0.176 | -0.087 | -0.046 | -0.014 | 0.058 | 30 | 7 |
| | AWC | | | 0.091 | -0.044 | 0.058 | 0.091 | 0.121 | 0.423 | 140 | 73 |
| | clay | -0.091 | 0.008 | -0.108 | -0.292 | -0.131 | -0.108 | -0.082 | 0.024 | 209 | 162 |
| | DeltFallMax | | | 0.057 | -0.415 | 0.013 | 0.057 | 0.093 | 0.129 | 38 | 14 |
| | DeltFallMin | | | 0.052 | -0.042 | 0.029 | 0.052 | 0.112 | 0.188 | 25 | 7 |
| | DeltPrecip | | | -0.066 | -0.113 | -0.08 | -0.066 | -0.039 | 0.055 | 124 | 62 |
| | DeltSpringMax | | | 0.038 | -0.107 | 0.004 | 0.038 | 0.051 | 0.107 | 27 | 4 |
| | heatload | | | -0.057 | -0.3 | -0.088 | -0.057 | -0.028 | 0.096 | 87 | 21 |
| | organic | | | 0.009 | -0.104 | -0.018 | 0.009 | 0.02 | 0.113 | 14 | 2 |
| | packm | | | -0.022 | -0.151 | -0.075 | -0.022 | -0.005 | 0.036 | 14 | 1 |
| | SixMonthPrecip | | | 0.053 | -0.071 | -0.018 | 0.053 | 0.071 | 0.116 | 17 | 5 |
| | Slope | | | 0.025 | -0.134 | -0.062 | 0.025 | 0.061 | 0.07 | 8 | 3 |
| | SoilpH | | | -0.057 | -0.19 | -0.089 | -0.057 | -0.034 | 0.033 | 42 | 13 |

| Species | Variable | Original model coefficient | Original model p | Median | Q 0% | Q 25% | Q 50% | Q 75% | Q 100% | Model Counts | P < 0.05 counts |
|---|---|---|---|---|---|---|---|---|---|---|---|
| | Ydays | | | 0.001 | -0.006 | 0 | 0.001 | 0.003 | 0.006 | 42 | 12 |
| | Year | | | -0.002 | -0.042 | -0.011 | -0.002 | -0.001 | 0.018 | 7 | 2 |
| *Mentzelia albicaulis* | AnnMinTemp | 0.011 | 0.800 | -0.022 | -0.142 | -0.053 | -0.022 | 0.013 | 0.227 | 165 | 56 |
| | Aspect | | | -0.03 | -0.082 | -0.053 | -0.03 | -0.005 | 0.058 | 13 | 5 |
| | AWC | -0.001 | 0.974 | 0.051 | -0.125 | 0.014 | 0.051 | 0.097 | 0.287 | 429 | 152 |
| | clay | 0.031 | 0.378 | 0.047 | -0.334 | 0.014 | 0.047 | 0.073 | 0.296 | 255 | 104 |
| | DeltFallMax | | | -0.056 | -0.133 | -0.085 | -0.056 | -0.03 | 0.146 | 31 | 13 |
| | DeltFallMin | -0.036 | 0.248 | -0.066 | -0.221 | -0.099 | -0.066 | -0.045 | 0.046 | 160 | 90 |
| | DeltPrecip | | | -0.049 | -0.145 | -0.07 | -0.049 | -0.036 | -0.003 | 24 | 14 |
| | DeltSpringMax | | | -0.016 | -0.079 | -0.04 | -0.016 | 0.007 | 0.161 | 26 | 10 |
| | heatload | -0.06 | 0.093 | -0.068 | -0.243 | -0.104 | -0.068 | -0.033 | 0.129 | 213 | 91 |
| | organic | 0.057 | 0.133 | 0.049 | -0.273 | 0.01 | 0.049 | 0.086 | 0.242 | 268 | 80 |
| | packm | | | -0.051 | -0.268 | -0.162 | -0.051 | 0.011 | 0.097 | 22 | 11 |
| | SixMonthPrecip | | | 0.041 | -0.129 | 0.004 | 0.041 | 0.063 | 0.175 | 51 | 12 |
| | Slope | | | 0.028 | -0.186 | 0.008 | 0.028 | 0.052 | 0.108 | 21 | 10 |

| Species | Variable | Original model coefficient | Original model p | Median | Q 0% | Q 25% | Q 50% | Q 75% | Q 100% | Model Counts | P < 0.05 counts |
|---|---|---|---|---|---|---|---|---|---|---|---|
| | SoilpH | | | -0.069 | -0.214 | -0.111 | -0.069 | -0.037 | 0.084 | 11 | 5 |
| | Ydays | | | 0.003 | -0.003 | 0.001 | 0.003 | 0.004 | 0.011 | 114 | 58 |
| | Year | | | 0.001 | -0.058 | -0.025 | 0.001 | 0.004 | 0.016 | 11 | 4 |
| *Mentzelia laevicaulis* | AnnMinTemp | 0.016 | 0.246 | 0.046 | -0.02 | 0.025 | 0.046 | 0.057 | 0.088 | 84 | 58 |
| | Aspect | | | 0.046 | -0.019 | 0.038 | 0.046 | 0.054 | 0.088 | 149 | 135 |
| | AWC | 0.004 | 0.806 | 0.011 | -0.062 | -0.012 | 0.011 | 0.036 | 0.055 | 30 | 11 |
| | clay | -0.031 | 0.158 | 0.003 | -0.08 | -0.019 | 0.003 | 0.025 | 0.063 | 24 | 7 |
| | DeltFallMax | | | 0.043 | -0.053 | 0.029 | 0.043 | 0.054 | 0.11 | 182 | 136 |
| | DeltFallMin | -0.026 | 0.047 | -0.017 | -0.07 | -0.029 | -0.017 | -0.004 | 0.022 | 82 | 21 |
| | DeltPrecip | | | -0.026 | -0.061 | -0.044 | -0.026 | -0.014 | 0.064 | 10 | 2 |
| | DeltSpringMax | | | -0.012 | -0.046 | -0.031 | -0.012 | 0.018 | 0.043 | 16 | 4 |
| | heatload | -0.023 | 0.099 | -0.027 | -0.052 | -0.037 | -0.027 | -0.016 | 0.04 | 44 | 22 |
| | organic | 0.047 | 0.010 | 0.03 | -0.022 | 0.022 | 0.03 | 0.037 | 0.084 | 105 | 73 |
| | packm | | | -0.019 | -0.077 | -0.048 | -0.019 | -0.002 | 0.07 | 12 | 7 |

| Species | Variable | Original model coefficient | Original model p | Median | Q 0% | Q 25% | Q 50% | Q 75% | Q 100% | Model Counts | P < 0.05 counts |
|---|---|---|---|---|---|---|---|---|---|---|---|
| | SixMonthPrecip | | | 0.015 | -0.03 | 0.005 | 0.015 | 0.028 | 0.081 | 34 | 11 |
| | Slope | | | 0.012 | -0.022 | -0.001 | 0.012 | 0.016 | 0.028 | 9 | 0 |
| | SoilpH | | | -0.033 | -0.101 | -0.044 | -0.033 | -0.022 | 0.071 | 114 | 71 |
| | Ydays | | | 0 | -0.002 | -0.001 | 0 | 0.002 | 0.002 | 8 | 4 |
| | Year | | | -0.012 | -0.028 | -0.013 | -0.012 | -0.008 | 0.013 | 98 | 69 |
| *Mimulus guttatus* | AnnMinTemp | | | 0.207 | -0.155 | 0.158 | 0.207 | 0.262 | 0.492 | 128 | 84 |
| | Aspect | | | 0.156 | -0.192 | 0.067 | 0.156 | 0.227 | 0.372 | 113 | 62 |
| | AWC | | | -0.115 | -0.194 | -0.158 | -0.115 | 0.1 | 0.28 | 9 | 2 |
| | clay | | | -0.061 | -0.224 | -0.164 | -0.061 | 0.116 | 0.268 | 7 | 3 |
| | DeltFallMax | | | -0.191 | -0.443 | -0.269 | -0.191 | -0.109 | -0.01 | 33 | 19 |
| | DeltFallMin | | | 0.028 | -0.422 | -0.04 | 0.028 | 0.126 | 0.321 | 111 | 20 |
| | DeltPrecip | | | -0.108 | -0.656 | -0.146 | -0.108 | -0.036 | 0.177 | 58 | 8 |
| | DeltSpringMax | | | -0.076 | -0.684 | -0.133 | -0.076 | -0.012 | 0.241 | 62 | 10 |
| | heatload | | | -0.171 | -0.486 | -0.242 | -0.171 | -0.124 | -0.051 | 39 | 23 |
| | organic | | | -0.076 | -0.149 | -0.097 | -0.076 | -0.063 | 0.692 | 9 | 1 |

| Species | Variable | Original model coefficient | Original model p | Median | Q 0% | Q 25% | Q 50% | Q 75% | Q 100% | Model Counts | P < 0.05 counts |
|---|---|---|---|---|---|---|---|---|---|---|---|
| | packm | | | 0.048 | -0.509 | -0.034 | 0.048 | 0.121 | 0.317 | 15 | 5 |
| | SixMonthPrecip | | | 0.215 | -0.036 | 0.076 | 0.215 | 0.299 | 0.731 | 17 | 8 |
| | Slope | | | -0.129 | -0.705 | -0.264 | -0.129 | -0.041 | 0.185 | 138 | 44 |
| | SoilpH | | | -0.154 | -0.324 | -0.194 | -0.154 | -0.035 | 0.091 | 12 | 2 |
| | Ydays | | | -0.003 | -0.01 | -0.005 | -0.003 | 0 | 0.007 | 94 | 16 |
| | Year | | | -0.025 | -0.061 | -0.036 | -0.025 | -0.022 | -0.017 | 4 | 2 |
| *Muhlenbergia porteri* | AnnMinTemp | -0.053 | 0.145 | -0.03 | -0.095 | -0.061 | -0.03 | 0.018 | 0.075 | 11 | 2 |
| | Aspect | | | 0.023 | -0.122 | -0.018 | 0.023 | 0.056 | 0.09 | 20 | 4 |
| | AWC | -0.037 | 0.350 | -0.088 | -0.322 | -0.136 | -0.088 | -0.035 | 0.142 | 98 | 39 |
| | clay | 0.068 | 0.249 | -0.061 | -0.308 | -0.118 | -0.061 | -0.005 | 0.175 | 126 | 26 |
| | DeltFallMax | | | -0.022 | -0.253 | -0.058 | -0.022 | 0.043 | 0.099 | 11 | 3 |
| | DeltFallMin | 0.111 | 0.004 | 0.077 | -0.234 | 0.033 | 0.077 | 0.118 | 0.244 | 136 | 62 |
| | DeltPrecip | | | -0.011 | -0.511 | -0.099 | -0.011 | 0.038 | 0.087 | 25 | 5 |
| | DeltSpringMax | | | -0.016 | -0.508 | -0.07 | -0.016 | 0.025 | 0.212 | 83 | 15 |

| Species | Variable | Original model coefficient | Original model p | Median | Q 0% | Q 25% | Q 50% | Q 75% | Q 100% | Model Counts | P < 0.05 counts |
|---|---|---|---|---|---|---|---|---|---|---|---|
| | heatload | -0.074 | 0.069 | -0.073 | -0.382 | -0.112 | -0.073 | -0.042 | 0.183 | 112 | 49 |
| | organic | -0.16 | 0.003 | -0.135 | -0.345 | -0.17 | -0.135 | -0.076 | 0.111 | 98 | 61 |
| | packm | | | -0.068 | -0.228 | -0.144 | -0.068 | 0.075 | 0.164 | 44 | 33 |
| | SixMonthPrecip | | | 0.115 | -0.077 | 0.067 | 0.115 | 0.168 | 0.216 | 18 | 8 |
| | Slope | | | -0.13 | -0.294 | -0.168 | -0.13 | -0.038 | 0.114 | 35 | 22 |
| | SoilpH | | | 0.109 | -0.202 | 0.041 | 0.109 | 0.16 | 0.411 | 153 | 68 |
| | Ydays | | | 0.002 | -0.009 | 0 | 0.002 | 0.003 | 0.016 | 40 | 5 |
| | Year | | | 0 | -0.082 | -0.029 | 0 | 0.023 | 0.082 | 42 | 10 |
| *Nicotiana attenuata* | AnnMinTemp | -0.025 | 0.014 | -0.019 | -0.037 | -0.023 | -0.019 | -0.012 | 0.034 | 9 | 2 |
| | Aspect | | | 0.035 | -0.049 | 0.022 | 0.035 | 0.046 | 0.08 | 114 | 68 |
| | AWC | 0.018 | 0.263 | -0.033 | -0.174 | -0.048 | -0.033 | -0.015 | 0.065 | 38 | 20 |
| | clay | -0.01 | 0.480 | -0.018 | -0.07 | -0.032 | -0.018 | -0.009 | 0.054 | 32 | 10 |
| | DeltFallMax | | | 0.036 | -0.041 | 0.025 | 0.036 | 0.047 | 0.091 | 171 | 121 |
| | DeltFallMin | 0.04 | <0.001 | 0.031 | -0.068 | 0.019 | 0.031 | 0.04 | 0.068 | 139 | 90 |
| | DeltPrecip | | | -0.012 | -0.036 | -0.019 | -0.012 | -0.007 | 0.038 | 6 | 2 |

| Species | Variable | Original model coefficient | Original model p | Median | Q 0% | Q 25% | Q 50% | Q 75% | Q 100% | Model Counts | P < 0.05 counts |
|---|---|---|---|---|---|---|---|---|---|---|---|
| | DeltSpringMax | | | -0.036 | -0.138 | -0.052 | -0.036 | -0.022 | 0.046 | 136 | 76 |
| | heatload | 0.023 | 0.046 | 0.031 | 0 | 0.022 | 0.031 | 0.039 | 0.067 | 33 | 20 |
| | organic | -0.056 | <0.001 | -0.038 | -0.161 | -0.042 | -0.038 | -0.024 | 0.106 | 41 | 25 |
| | packm | | | 0.028 | -0.039 | 0.021 | 0.028 | 0.035 | 0.041 | 38 | 23 |
| | SixMonthPrecip | | | 0.019 | -0.073 | -0.006 | 0.019 | 0.044 | 0.095 | 140 | 70 |
| | Slope | | | -0.019 | -0.071 | -0.042 | -0.019 | 0.015 | 0.024 | 20 | 10 |
| | SoilpH | | | -0.018 | -0.038 | -0.023 | -0.018 | -0.006 | 0.011 | 15 | 4 |
| | Ydays | | | 0.001 | -0.004 | 0 | 0.001 | 0.001 | 0.003 | 23 | 9 |
| | Year | | | 0.007 | -0.019 | 0.001 | 0.007 | 0.011 | 0.024 | 142 | 64 |
| *Pectis papposa* | AnnMinTemp | | | -0.05 | -0.162 | -0.077 | -0.05 | -0.006 | 0.116 | 71 | 27 |
| | Aspect | | | 0.044 | -0.027 | 0.02 | 0.044 | 0.051 | 0.069 | 8 | 1 |
| | AWC | | | -0.016 | -0.152 | -0.035 | -0.016 | 0.001 | 0.073 | 9 | 2 |
| | clay | | | -0.033 | -0.105 | -0.047 | -0.033 | 0.007 | 0.067 | 20 | 5 |
| | DeltFallMax | | | 0.034 | -0.049 | 0.007 | 0.034 | 0.074 | 0.168 | 24 | 9 |
| | DeltFallMin | | | -0.012 | -0.088 | -0.036 | -0.012 | 0.009 | 0.084 | 48 | 5 |

| Species | Variable | Original model coefficient | Original model p | Median | Q 0% | Q 25% | Q 50% | Q 75% | Q 100% | Model Counts | P < 0.05 counts |
|---|---|---|---|---|---|---|---|---|---|---|---|
| | DeltPrecip | | | -0.048 | -0.191 | -0.068 | -0.048 | -0.022 | 0.08 | 153 | 57 |
| | DeltSpringMax | | | 0.016 | -0.115 | -0.01 | 0.016 | 0.049 | 0.07 | 7 | 2 |
| | heatload | | | 0.028 | -0.148 | 0.006 | 0.028 | 0.057 | 0.151 | 22 | 7 |
| | organic | | | -0.023 | -0.229 | -0.066 | -0.023 | 0.029 | 0.506 | 61 | 13 |
| | packm | 0.049 | 0.050 | 0.059 | -0.058 | 0.028 | 0.059 | 0.097 | 0.151 | 234 | 125 |
| | SixMonthPrecip | | | 0.053 | -0.125 | 0.016 | 0.053 | 0.084 | 0.153 | 100 | 55 |
| | Slope | | | 0.066 | -0.235 | -0.005 | 0.066 | 0.11 | 0.218 | 14 | 5 |
| | SoilpH | 0.027 | 0.287 | 0.045 | -0.088 | 0.016 | 0.045 | 0.1 | 0.564 | 160 | 47 |
| | Ydays | | | -0.001 | -0.007 | -0.004 | -0.001 | 0.002 | 0.005 | 31 | 13 |
| | Year | 0.07 | <0.001 | 0.065 | -0.017 | 0.053 | 0.065 | 0.079 | 0.197 | 490 | 449 |
| *Penstemon deustus* | AnnMinTemp | | | 0.077 | -0.061 | 0.049 | 0.077 | 0.105 | 0.178 | 86 | 49 |
| | Aspect | | | -0.095 | -0.224 | -0.108 | -0.095 | -0.066 | 0.073 | 10 | 5 |
| | AWC | 0.046 | 0.321 | 0.02 | -0.277 | -0.019 | 0.02 | 0.057 | 0.19 | 175 | 44 |
| | clay | 0.04 | 0.410 | 0.083 | -0.137 | 0.052 | 0.083 | 0.116 | 0.422 | 368 | 194 |
| | DeltFallMax | | | -0.046 | -0.11 | -0.065 | -0.046 | -0.029 | 0.007 | 17 | 7 |

| Species | Variable | Original model coefficient | Original model p | Median | Q 0% | Q 25% | Q 50% | Q 75% | Q 100% | Model Counts | P < 0.05 counts |
|---|---|---|---|---|---|---|---|---|---|---|---|
| | DeltFallMin | | | 0.068 | -0.086 | 0.025 | 0.068 | 0.077 | 0.263 | 13 | 4 |
| | DeltPrecip | | | -0.022 | -0.194 | -0.112 | -0.022 | 0.021 | 0.091 | 13 | 6 |
| | DeltSpringMax | | | 0.062 | -0.099 | 0.026 | 0.062 | 0.091 | 0.201 | 19 | 10 |
| | heatload | | | -0.021 | -0.258 | -0.066 | -0.021 | 0.023 | 0.125 | 99 | 28 |
| | organic | | | -0.031 | -0.227 | -0.061 | -0.031 | 0.025 | 0.23 | 49 | 13 |
| | packm | | | -0.094 | -0.232 | -0.127 | -0.094 | -0.072 | 0.021 | 32 | 25 |
| | SixMonthPrecip | | | -0.029 | -0.149 | -0.052 | -0.029 | 0.008 | 0.158 | 56 | 10 |
| | Slope | | | 0.006 | -0.218 | -0.023 | 0.006 | 0.037 | 0.113 | 67 | 14 |
| | SoilpH | 0.078 | 0.055 | 0.046 | -0.222 | 0.011 | 0.046 | 0.079 | 0.283 | 316 | 75 |
| | Ydays | | | 0.001 | -0.004 | -0.002 | 0.001 | 0.002 | 0.006 | 57 | 20 |
| | Year | | | -0.051 | -0.078 | -0.063 | -0.051 | -0.04 | -0.004 | 6 | 4 |
| *Plantago ovata* | AnnMinTemp | -0.088 | <0.001 | -0.087 | -0.19 | -0.112 | -0.087 | -0.055 | 0.117 | 302 | 216 |
| | Aspect | | | 0.049 | -0.057 | 0.038 | 0.049 | 0.085 | 0.134 | 39 | 24 |
| | AWC | | | -0.042 | -0.102 | -0.071 | -0.042 | 0 | 0.053 | 43 | 20 |
| | clay | -0.028 | 0.261 | -0.046 | -0.171 | -0.071 | -0.046 | -0.016 | 0.06 | 211 | 73 |

| Species | Variable | Original model coefficient | Original model p | Median | Q 0% | Q 25% | Q 50% | Q 75% | Q 100% | Model Counts | P < 0.05 counts |
|---|---|---|---|---|---|---|---|---|---|---|---|
| | DeltFallMax | | | 0.078 | -0.013 | 0.056 | 0.078 | 0.084 | 0.13 | 30 | 23 |
| | DeltFallMin | | | 0.063 | -0.026 | 0.037 | 0.063 | 0.091 | 0.152 | 55 | 32 |
| | DeltPrecip | | | -0.055 | -0.149 | -0.09 | -0.055 | -0.028 | 0.062 | 32 | 18 |
| | DeltSpringMax | | | 0.062 | -0.122 | 0.038 | 0.062 | 0.098 | 0.217 | 75 | 49 |
| | heatload | -0.01 | 0.753 | -0.043 | -0.343 | -0.085 | -0.043 | -0.015 | 0.14 | 207 | 92 |
| | organic | | | -0.029 | -0.24 | -0.059 | -0.029 | 0.023 | 0.271 | 99 | 30 |
| | packm | | | 0.076 | -0.097 | 0.035 | 0.076 | 0.116 | 0.225 | 98 | 62 |
| | SixMonthPrecip | | | -0.028 | -0.078 | -0.053 | -0.028 | 0.045 | 0.08 | 17 | 4 |
| | Slope | 0.035 | 0.223 | 0.041 | -0.229 | 0.021 | 0.041 | 0.07 | 0.307 | 176 | 61 |
| | SoilpH | 0.052 | 0.017 | 0.059 | -0.082 | 0.036 | 0.059 | 0.113 | 0.338 | 322 | 153 |
| | Ydays | | | 0.002 | -0.006 | 0 | 0.002 | 0.004 | 0.011 | 60 | 23 |
| | Year | | | -0.007 | -0.056 | -0.027 | -0.007 | 0.049 | 0.067 | 14 | 8 |
| *Plantago patagonica* | AnnMinTemp | 0.117 | <0.001 | 0.096 | -0.157 | 0.055 | 0.096 | 0.139 | 0.273 | 430 | 295 |
| | Aspect | | | 0.024 | -0.106 | -0.01 | 0.024 | 0.051 | 0.213 | 34 | 11 |

| Species | Variable | Original model coefficient | Original model p | Median | Q 0% | Q 25% | Q 50% | Q 75% | Q 100% | Model Counts | P < 0.05 counts |
|---|---|---|---|---|---|---|---|---|---|---|---|
| | AWC | | | 0.05 | -0.098 | 0.009 | 0.05 | 0.067 | 0.126 | 41 | 17 |
| | clay | 0.081 | <0.001 | 0.087 | -0.057 | 0.053 | 0.087 | 0.121 | 0.291 | 235 | 168 |
| | DeltFallMax | | | -0.057 | -0.131 | -0.083 | -0.057 | -0.039 | 0.025 | 37 | 19 |
| | DeltFallMin | | | 0.041 | -0.001 | 0.032 | 0.041 | 0.066 | 0.103 | 13 | 6 |
| | DeltPrecip | -0.001 | 0.962 | 0.043 | -0.176 | 0 | 0.043 | 0.086 | 0.222 | 214 | 90 |
| | DeltSpringMax | | | -0.007 | -0.135 | -0.047 | -0.007 | 0.04 | 0.141 | 67 | 21 |
| | heatload | | | 0.003 | -0.17 | -0.04 | 0.003 | 0.046 | 0.107 | 42 | 13 |
| | organic | 0.012 | 0.522 | 0.039 | -0.111 | -0.006 | 0.039 | 0.087 | 0.243 | 90 | 42 |
| | packm | | | -0.003 | -0.14 | -0.047 | -0.003 | 0.04 | 0.116 | 26 | 8 |
| | SixMonthPrecip | | | 0.067 | -0.226 | 0.021 | 0.067 | 0.103 | 0.228 | 105 | 54 |
| | Slope | | | -0.032 | -0.083 | -0.053 | -0.032 | 0.01 | 0.09 | 8 | 5 |
| | SoilpH | | | -0.01 | -0.12 | -0.061 | -0.01 | 0.036 | 0.195 | 60 | 21 |
| | Ydays | 0 | 0.660 | -0.002 | -0.011 | -0.004 | -0.002 | 0 | 0.007 | 318 | 133 |
| | Year | -0.024 | 0.001 | -0.025 | -0.097 | -0.038 | -0.025 | -0.012 | 0.044 | 246 | 139 |
| *Pleuraphis jamesii* | AnnMinTemp | | | 0.067 | -0.118 | 0.039 | 0.067 | 0.103 | 0.161 | 12 | 6 |

| Species | Variable | Original model coefficient | Original model p | Median | Q 0% | Q 25% | Q 50% | Q 75% | Q 100% | Model Counts | P < 0.05 counts |
|---|---|---|---|---|---|---|---|---|---|---|---|
| | Aspect | | | -0.073 | -0.073 | -0.073 | -0.073 | -0.073 | -0.073 | 1 | 1 |
| | AWC | | | -0.05 | -0.233 | -0.099 | -0.05 | -0.022 | 0.131 | 61 | 26 |
| | clay | | | 0.112 | 0.061 | 0.076 | 0.112 | 0.113 | 0.124 | 5 | 3 |
| | DeltFallMax | | | -0.071 | -0.195 | -0.118 | -0.071 | 0.023 | 0.153 | 8 | 5 |
| | DeltFallMin | 0.14 | 0.001 | 0.12 | -0.253 | 0.079 | 0.12 | 0.173 | 0.328 | 338 | 241 |
| | DeltPrecip | | | -0.019 | -0.206 | -0.07 | -0.019 | 0.031 | 0.237 | 116 | 30 |
| | DeltSpringMax | -0.047 | 0.373 | -0.059 | -0.311 | -0.104 | -0.059 | 0 | 0.273 | 334 | 105 |
| | heatload | | | 0.067 | -0.019 | 0.045 | 0.067 | 0.129 | 0.241 | 6 | 4 |
| | organic | | | 0.094 | -0.133 | 0.067 | 0.094 | 0.131 | 0.301 | 71 | 45 |
| | packm | | | -0.032 | -0.032 | -0.032 | -0.032 | -0.032 | -0.032 | 1 | 0 |
| | SixMonthPrecip | | | 0.043 | -0.146 | 0.006 | 0.043 | 0.116 | 0.265 | 88 | 35 |
| | Slope | | | 0.027 | -0.1 | 0.018 | 0.027 | 0.043 | 0.166 | 17 | 5 |
| | SoilpH | | | 0.074 | -0.171 | 0.03 | 0.074 | 0.113 | 0.218 | 81 | 43 |
| | Ydays | -0.001 | 0.303 | -0.002 | -0.008 | -0.003 | -0.002 | -0.001 | 0.005 | 182 | 75 |
| | Year | 0.014 | 0.466 | 0.033 | -0.098 | 0.011 | 0.033 | 0.053 | 0.157 | 495 | 226 |

| Species | Variable | Original model coefficient | Original model p | Median | Q 0% | Q 25% | Q 50% | Q 75% | Q 100% | Model Counts | P < 0.05 counts |
|---|---|---|---|---|---|---|---|---|---|---|---|
| *Poa secunda* | AnnMinTemp | | | 0.135 | -0.072 | 0.075 | 0.135 | 0.161 | 0.327 | 34 | 23 |
| | Aspect | | | -0.128 | -0.314 | -0.208 | -0.128 | 0.008 | 0.132 | 34 | 21 |
| | AWC | | | -0.104 | -0.272 | -0.124 | -0.104 | -0.064 | 0.027 | 22 | 9 |
| | clay | 0.058 | 0.036 | 0.103 | 0.006 | 0.08 | 0.103 | 0.134 | 0.549 | 100 | 62 |
| | DeltFallMax | | | -0.089 | -0.32 | -0.138 | -0.089 | -0.024 | 0.083 | 40 | 19 |
| | DeltFallMin | | | 0.074 | -0.166 | 0.028 | 0.074 | 0.127 | 0.32 | 47 | 19 |
| | DeltPrecip | | | 0.046 | -0.219 | -0.14 | 0.046 | 0.099 | 0.268 | 27 | 17 |
| | DeltSpringMax | | | -0.054 | -0.217 | -0.102 | -0.054 | 0.034 | 0.127 | 18 | 9 |
| | heatload | | | 0.071 | -0.108 | 0.035 | 0.071 | 0.098 | 0.272 | 34 | 8 |
| | organic | | | 0.066 | -1.613 | -0.043 | 0.066 | 0.354 | 1.272 | 64 | 10 |
| | packm | | | -0.092 | -0.206 | -0.133 | -0.092 | -0.073 | 0.261 | 15 | 10 |
| | SixMonthPrecip | 0.086 | 0.005 | 0.127 | -0.038 | 0.079 | 0.127 | 0.174 | 0.457 | 221 | 118 |
| | Slope | | | 0.058 | -0.09 | -0.034 | 0.058 | 0.093 | 0.269 | 27 | 7 |
| | SoilpH | -0.051 | 0.094 | -0.118 | -0.782 | -0.163 | -0.118 | -0.074 | 0.229 | 98 | 49 |
| | Ydays | | | 0.003 | -0.008 | -0.001 | 0.003 | 0.004 | 0.014 | 30 | 11 |

| Species | Variable | Original model coefficient | Original model p | Median | Q 0% | Q 25% | Q 50% | Q 75% | Q 100% | Model Counts | P < 0.05 counts |
|---|---|---|---|---|---|---|---|---|---|---|---|
| | Year | 0.004 | 0.678 | -0.005 | -0.039 | -0.023 | -0.005 | 0.019 | 0.07 | 36 | 10 |
| *Pseudoroegneria spicata* | AnnMinTemp | | | 0.055 | -0.115 | 0.024 | 0.055 | 0.08 | 0.158 | 60 | 37 |
| | Aspect | | | -0.056 | -0.224 | -0.078 | -0.056 | -0.004 | 0.103 | 42 | 24 |
| | AWC | | | -0.032 | -0.097 | -0.064 | -0.032 | 0.019 | 0.108 | 19 | 10 |
| | clay | | | 0.057 | -0.076 | 0.01 | 0.057 | 0.081 | 0.147 | 41 | 18 |
| | DeltFallMax | | | -0.059 | -0.148 | -0.078 | -0.059 | -0.043 | 0.029 | 29 | 17 |
| | DeltFallMin | | | 0.008 | -0.094 | -0.055 | 0.008 | 0.041 | 0.059 | 17 | 5 |
| | DeltPrecip | 0.016 | 0.446 | -0.025 | -0.136 | -0.057 | -0.025 | 0.023 | 0.124 | 91 | 25 |
| | DeltSpringMax | -0.051 | 0.012 | -0.063 | -0.238 | -0.086 | -0.063 | -0.021 | 0.225 | 97 | 55 |
| | heatload | | | 0.017 | -0.113 | -0.047 | 0.017 | 0.064 | 0.2 | 65 | 27 |
| | organic | -0.049 | 0.004 | -0.044 | -0.461 | -0.131 | -0.044 | -0.023 | 0.448 | 69 | 37 |
| | packm | | | -0.032 | -0.15 | -0.048 | -0.032 | 0.011 | 0.114 | 38 | 10 |
| | SixMonthPrecip | | | 0.057 | -0.038 | 0.041 | 0.057 | 0.073 | 0.138 | 57 | 31 |
| | Slope | | | 0.061 | -0.139 | 0.014 | 0.061 | 0.086 | 0.148 | 47 | 29 |

| Species | Variable | Original model coefficient | Original model p | Median | Q 0% | Q 25% | Q 50% | Q 75% | Q 100% | Model Counts | P < 0.05 counts |
|---|---|---|---|---|---|---|---|---|---|---|---|
| | SoilpH | -0.041 | 0.032 | -0.066 | -0.12 | -0.079 | -0.066 | -0.044 | 0.009 | 80 | 50 |
| | Ydays | | | 0.002 | -0.007 | 0 | 0.002 | 0.003 | 0.006 | 49 | 12 |
| | Year | | | 0.011 | -0.04 | 0 | 0.011 | 0.018 | 0.045 | 58 | 20 |
| *Purshia stansburiana* | AnnMinTemp | | | -0.02 | -0.122 | -0.046 | -0.02 | 0 | 0.078 | 40 | 13 |
| | Aspect | | | 0.061 | -0.094 | 0.047 | 0.061 | 0.081 | 0.143 | 54 | 43 |
| | AWC | 0.061 | 0.004 | 0.063 | -0.033 | 0.043 | 0.063 | 0.082 | 0.241 | 390 | 291 |
| | clay | 0.028 | 0.160 | 0.023 | -0.294 | 0.003 | 0.023 | 0.043 | 0.171 | 405 | 110 |
| | DeltFallMax | | | -0.027 | -0.048 | -0.037 | -0.027 | -0.012 | 0.004 | 3 | 1 |
| | DeltFallMin | | | 0.019 | -0.057 | -0.008 | 0.019 | 0.038 | 0.13 | 50 | 18 |
| | DeltPrecip | | | 0.046 | 0.003 | 0.034 | 0.046 | 0.064 | 0.107 | 37 | 27 |
| | DeltSpringMax | 0.005 | 0.824 | 0.013 | -0.221 | -0.01 | 0.013 | 0.038 | 0.188 | 332 | 80 |
| | heatload | | | 0.045 | 0.012 | 0.027 | 0.045 | 0.055 | 0.078 | 9 | 5 |
| | organic | -0.005 | 0.838 | 0.012 | -0.121 | -0.014 | 0.012 | 0.04 | 0.202 | 435 | 98 |
| | packm | | | 0.041 | -0.182 | 0.002 | 0.041 | 0.071 | 0.179 | 110 | 38 |

| Species | Variable | Original model coefficient | Original model p | Median | Q 0% | Q 25% | Q 50% | Q 75% | Q 100% | Model Counts | P < 0.05 counts |
|---|---|---|---|---|---|---|---|---|---|---|---|
| | SixMonthPrecip | | | 0.011 | -0.056 | -0.007 | 0.011 | 0.029 | 0.125 | 78 | 16 |
| | Slope | | | -0.035 | -0.151 | -0.082 | -0.035 | 0.004 | 0.063 | 62 | 29 |
| | SoilpH | -0.087 | <0.001 | -0.07 | -0.206 | -0.095 | -0.07 | -0.041 | 0.151 | 490 | 332 |
| | Ydays | | | -0.003 | -0.003 | -0.003 | -0.003 | -0.002 | -0.002 | 3 | 3 |
| | Year | | | -0.009 | -0.041 | -0.019 | -0.009 | -0.002 | 0.009 | 38 | 11 |
| *Purshia tridentata* | AnnMinTemp | | | 0.053 | -0.076 | 0.021 | 0.053 | 0.084 | 0.194 | 37 | 12 |
| | Aspect | | | -0.071 | -0.164 | -0.093 | -0.071 | -0.047 | 0.089 | 35 | 19 |
| | AWC | | | 0.055 | -0.129 | 0.022 | 0.055 | 0.068 | 0.122 | 32 | 12 |
| | clay | -0.009 | 0.722 | -0.067 | -0.282 | -0.114 | -0.067 | -0.029 | 0.114 | 91 | 42 |
| | DeltFallMax | | | 0.048 | -0.104 | -0.042 | 0.048 | 0.065 | 0.102 | 19 | 10 |
| | DeltFallMin | | | -0.08 | -0.169 | -0.101 | -0.08 | -0.041 | 0.078 | 48 | 33 |
| | DeltPrecip | | | 0.051 | -0.146 | 0 | 0.051 | 0.073 | 0.121 | 25 | 13 |
| | DeltSpringMax | 0.011 | 0.676 | 0.012 | -0.159 | -0.023 | 0.012 | 0.041 | 0.133 | 77 | 13 |
| | heatload | | | -0.012 | -0.208 | -0.063 | -0.012 | 0.098 | 0.129 | 33 | 18 |
| | organic | | | 0.047 | -0.155 | -0.035 | 0.047 | 0.086 | 0.167 | 38 | 22 |

| Species | Variable | Original model coefficient | Original model p | Median | Q 0% | Q 25% | Q 50% | Q 75% | Q 100% | Model Counts | P < 0.05 counts |
|---|---|---|---|---|---|---|---|---|---|---|---|
| | packm | 0.027 | 0.291 | 0.067 | -0.106 | 0.042 | 0.067 | 0.094 | 0.192 | 97 | 53 |
| | SixMonthPrecip | 0.026 | 0.222 | 0.123 | -0.033 | 0.03 | 0.123 | 0.175 | 0.348 | 102 | 56 |
| | Slope | | | -0.057 | -0.245 | -0.083 | -0.057 | -0.045 | 0.024 | 25 | 12 |
| | SoilpH | 0.06 | 0.006 | 0.081 | -0.037 | 0.06 | 0.081 | 0.097 | 0.171 | 162 | 113 |
| | Ydays | 0.001 | 0.275 | 0.004 | -0.004 | 0.003 | 0.004 | 0.006 | 0.009 | 106 | 60 |
| | Year | | | -0.021 | -0.066 | -0.033 | -0.021 | -0.013 | 0.022 | 59 | 26 |
| *Ratibida columnifera* | AnnMinTemp | | | -0.057 | -0.353 | -0.072 | -0.057 | -0.035 | 0.086 | 194 | 87 |
| | Aspect | -0.07 | 0.012 | -0.085 | -0.172 | -0.101 | -0.085 | -0.072 | -0.03 | 255 | 244 |
| | AWC | | | 0.049 | -0.063 | 0.028 | 0.049 | 0.058 | 0.128 | 45 | 22 |
| | clay | | | 0.004 | -0.075 | -0.043 | 0.004 | 0.127 | 0.19 | 8 | 5 |
| | DeltFallMax | | | 0.049 | -0.128 | 0.027 | 0.049 | 0.07 | 0.256 | 127 | 63 |
| | DeltFallMin | | | -0.037 | -0.176 | -0.057 | -0.037 | -0.017 | 0.067 | 77 | 23 |
| | DeltPrecip | | | -0.038 | -0.076 | -0.057 | -0.038 | 0.016 | 0.07 | 3 | 1 |
| | DeltSpringMax | | | 0.061 | -0.044 | 0.026 | 0.061 | 0.093 | 0.264 | 8 | 4 |

| Species | Variable | Original model coefficient | Original model p | Median | Q 0% | Q 25% | Q 50% | Q 75% | Q 100% | Model Counts | P < 0.05 counts |
|---|---|---|---|---|---|---|---|---|---|---|---|
| | heatload | | | -0.006 | -0.063 | -0.028 | -0.006 | 0.008 | 0.106 | 54 | 4 |
| | organic | | | 0.036 | -0.078 | 0.026 | 0.036 | 0.115 | 0.203 | 11 | 6 |
| | packm | | | 0.043 | 0.001 | 0.028 | 0.043 | 0.063 | 0.156 | 58 | 21 |
| | SixMonthPrecip | | | 0.059 | -0.122 | 0.034 | 0.059 | 0.085 | 0.169 | 180 | 96 |
| | Slope | | | 0.013 | -0.094 | -0.004 | 0.013 | 0.044 | 0.164 | 17 | 6 |
| | SoilpH | | | 0.03 | -0.01 | 0.006 | 0.03 | 0.044 | 0.109 | 10 | 2 |
| | Ydays | | | 0.002 | -0.003 | 0 | 0.002 | 0.003 | 0.005 | 63 | 27 |
| | Year | | | -0.015 | -0.068 | -0.029 | -0.015 | -0.006 | 0.033 | 29 | 13 |
| *Rosa woodsii* | AnnMinTemp | | | -0.075 | -0.392 | -0.106 | -0.075 | -0.041 | 0.104 | 172 | 97 |
| | Aspect | | | 0.074 | -0.033 | 0.056 | 0.074 | 0.097 | 0.158 | 13 | 8 |
| | AWC | | | -0.02 | -0.114 | -0.024 | -0.02 | 0.153 | 0.197 | 9 | 4 |
| | clay | | | -0.014 | -0.133 | -0.068 | -0.014 | 0.059 | 0.137 | 13 | 5 |
| | DeltFallMax | | | -0.103 | -0.181 | -0.126 | -0.103 | -0.058 | 0.01 | 24 | 13 |
| | DeltFallMin | | | -0.013 | -0.202 | -0.049 | -0.013 | -0.003 | 0.132 | 8 | 3 |
| | DeltPrecip | | | -0.07 | -0.266 | -0.11 | -0.07 | 0 | 0.039 | 7 | 2 |

| Species | Variable | Original model coefficient | Original model p | Median | Q 0% | Q 25% | Q 50% | Q 75% | Q 100% | Model Counts | P < 0.05 counts |
|---|---|---|---|---|---|---|---|---|---|---|---|
| | DeltSpringMax | 0.027 | 0.416 | 0.061 | -0.187 | 0.027 | 0.061 | 0.097 | 0.334 | 193 | 77 |
| | heatload | | | 0.016 | -0.156 | -0.037 | 0.016 | 0.085 | 0.173 | 13 | 5 |
| | organic | | | 0.018 | -0.231 | -0.01 | 0.018 | 0.042 | 0.205 | 96 | 21 |
| | packm | | | 0.053 | -0.077 | 0.027 | 0.053 | 0.091 | 0.253 | 124 | 49 |
| | SixMonthPrecip | 0.116 | <0.001 | 0.11 | -0.068 | 0.07 | 0.11 | 0.14 | 0.32 | 291 | 212 |
| | Slope | | | -0.076 | -0.248 | -0.096 | -0.076 | -0.054 | 0.066 | 53 | 33 |
| | SoilpH | 0.063 | 0.022 | 0.082 | -0.146 | 0.047 | 0.082 | 0.119 | 0.229 | 127 | 80 |
| | Ydays | | | -0.001 | -0.007 | -0.002 | -0.001 | 0 | 0.002 | 45 | 8 |
| | Year | -0.021 | 0.059 | -0.027 | -0.082 | -0.036 | -0.027 | -0.019 | 0.023 | 169 | 102 |
| *Salvia columbariae* | AnnMinTemp | | | -0.067 | -0.185 | -0.086 | -0.067 | -0.038 | 0.047 | 100 | 67 |
| | Aspect | | | -0.044 | -0.086 | -0.054 | -0.044 | -0.038 | 0.129 | 26 | 14 |
| | AWC | | | -0.052 | -0.135 | -0.08 | -0.052 | -0.021 | 0.09 | 62 | 29 |
| | clay | | | -0.067 | -0.124 | -0.081 | -0.067 | -0.054 | 0.001 | 128 | 97 |
| | DeltFallMax | | | 0.046 | -0.012 | 0.031 | 0.046 | 0.059 | 0.176 | 53 | 23 |
| | DeltFallMin | | | 0.012 | -0.08 | -0.036 | 0.012 | 0.041 | 0.099 | 42 | 13 |

| Species | Variable | Original model coefficient | Original model p | Median | Q 0% | Q 25% | Q 50% | Q 75% | Q 100% | Model Counts | P < 0.05 counts |
|---|---|---|---|---|---|---|---|---|---|---|---|
| | DeltPrecip | | | 0.049 | -0.051 | 0.031 | 0.049 | 0.071 | 0.104 | 18 | 10 |
| | DeltSpringMax | 0.05 | 0.075 | 0.046 | -0.047 | 0.02 | 0.046 | 0.07 | 0.129 | 60 | 20 |
| | heatload | | | -0.053 | -0.114 | -0.074 | -0.053 | -0.032 | 0.015 | 44 | 23 |
| | organic | | | -0.046 | -0.3 | -0.061 | -0.046 | -0.027 | 0.439 | 181 | 80 |
| | packm | | | -0.022 | -0.074 | -0.048 | -0.022 | 0.008 | 0.039 | 3 | 2 |
| | SixMonthPrecip | -0.047 | 0.110 | -0.054 | -0.117 | -0.074 | -0.054 | -0.036 | 0.032 | 33 | 18 |
| | Slope | | | -0.034 | -0.095 | -0.069 | -0.034 | 0.009 | 0.065 | 34 | 14 |
| | SoilpH | 0.012 | 0.634 | 0.054 | -0.044 | 0.033 | 0.054 | 0.068 | 0.188 | 132 | 62 |
| | Ydays | | | -0.001 | -0.002 | -0.002 | -0.001 | -0.001 | 0 | 11 | 4 |
| | Year | 0.005 | 0.518 | -0.012 | -0.022 | -0.017 | -0.012 | -0.008 | -0.002 | 8 | 3 |
| *Sedum lanceolatum* | AnnMinTemp | | | 0.101 | -0.218 | 0.014 | 0.101 | 0.147 | 0.304 | 64 | 29 |
| | Aspect | | | 0.013 | -0.224 | -0.081 | 0.013 | 0.057 | 0.107 | 16 | 6 |
| | AWC | 0.051 | 0.402 | 0.074 | -0.734 | 0.023 | 0.074 | 0.18 | 0.594 | 366 | 100 |
| | clay | | | 0.043 | -0.456 | -0.021 | 0.043 | 0.101 | 0.284 | 57 | 13 |
| | DeltFallMax | | | 0.112 | -0.009 | 0.075 | 0.112 | 0.154 | 0.299 | 96 | 42 |

| Species | Variable | Original model coefficient | Original model p | Median | Q 0% | Q 25% | Q 50% | Q 75% | Q 100% | Model Counts | P < 0.05 counts |
|---|---|---|---|---|---|---|---|---|---|---|---|
| | DeltFallMin | | | -0.195 | -0.503 | -0.241 | -0.195 | -0.108 | 0.003 | 12 | 9 |
| | DeltPrecip | | | 0.123 | -0.118 | 0.085 | 0.123 | 0.184 | 0.514 | 38 | 19 |
| | DeltSpringMax | | | -0.076 | -0.204 | -0.115 | -0.076 | -0.048 | 0.107 | 10 | 2 |
| | heatload | | | 0 | -1.375 | -0.086 | 0 | 0.085 | 0.33 | 93 | 25 |
| | organic | | | -0.095 | -0.564 | -0.218 | -0.095 | 0.102 | 1.446 | 21 | 11 |
| | packm | | | -0.104 | -0.319 | -0.143 | -0.104 | -0.04 | 0.422 | 54 | 33 |
| | SixMonthPrecip | | | 0.095 | -0.131 | 0.052 | 0.095 | 0.163 | 0.276 | 6 | 4 |
| | Slope | | | -0.223 | -1.124 | -0.355 | -0.223 | -0.156 | 0.086 | 59 | 48 |
| | SoilpH | | | 0.107 | -0.15 | 0.028 | 0.107 | 0.16 | 1.249 | 48 | 22 |
| | Ydays | | | 0.004 | -0.008 | -0.003 | 0.004 | 0.008 | 0.033 | 25 | 11 |
| | Year | | | 0.009 | -0.056 | -0.031 | 0.009 | 0.009 | 0.027 | 5 | 2 |
| *Sphaeralcea ambigua* | AnnMinTemp | | | 0.049 | -0.044 | 0.032 | 0.049 | 0.067 | 0.118 | 135 | 71 |
| | Aspect | | | -0.053 | -0.196 | -0.068 | -0.053 | -0.038 | 0.037 | 67 | 46 |
| | AWC | | | -0.048 | -0.16 | -0.07 | -0.048 | 0.002 | 0.104 | 56 | 34 |

| Species | Variable | Original model coefficient | Original model p | Median | Q 0% | Q 25% | Q 50% | Q 75% | Q 100% | Model Counts | P < 0.05 counts |
|---|---|---|---|---|---|---|---|---|---|---|---|
| | clay | | | -0.069 | -0.203 | -0.086 | -0.069 | -0.048 | 0.027 | 127 | 89 |
| | DeltFallMax | | | -0.033 | -0.131 | -0.053 | -0.033 | -0.011 | 0.097 | 101 | 38 |
| | DeltFallMin | | | -0.045 | -0.117 | -0.071 | -0.045 | -0.012 | 0.121 | 33 | 19 |
| | DeltPrecip | | | -0.051 | -0.105 | -0.076 | -0.051 | -0.031 | 0.025 | 22 | 9 |
| | DeltSpringMax | | | -0.033 | -0.184 | -0.062 | -0.033 | -0.012 | 0.165 | 45 | 15 |
| | heatload | | | 0.008 | -0.087 | -0.035 | 0.008 | 0.061 | 0.168 | 45 | 19 |
| | organic | | | -0.044 | -0.278 | -0.072 | -0.044 | -0.027 | 0.064 | 100 | 49 |
| | packm | | | -0.047 | -0.158 | -0.069 | -0.047 | -0.033 | -0.008 | 26 | 9 |
| | SixMonthPrecip | | | 0.067 | -0.078 | 0.049 | 0.067 | 0.077 | 0.108 | 23 | 18 |
| | Slope | | | 0.04 | -0.088 | 0.012 | 0.04 | 0.067 | 0.138 | 56 | 28 |
| | SoilpH | | | 0.038 | -0.096 | -0.001 | 0.038 | 0.051 | 0.068 | 10 | 4 |
| | Ydays | -0.002 | 0.001 | -0.002 | -0.006 | -0.003 | -0.002 | -0.001 | 0.004 | 228 | 148 |
| | Year | | | 0.004 | -0.034 | -0.012 | 0.004 | 0.015 | 0.035 | 37 | 19 |
| *Sphaeralcea coccinea* | AnnMinTemp | | | -0.133 | -0.463 | -0.199 | -0.133 | -0.06 | 0.109 | 94 | 54 |

| Species | Variable | Original model coefficient | Original model p | Median | Q 0% | Q 25% | Q 50% | Q 75% | Q 100% | Model Counts | P < 0.05 counts |
|---|---|---|---|---|---|---|---|---|---|---|---|
| | Aspect | | | -0.002 | -0.181 | -0.107 | -0.002 | 0.067 | 0.18 | 34 | 18 |
| | AWC | | | 0.049 | -0.098 | 0.012 | 0.049 | 0.143 | 0.315 | 14 | 7 |
| | clay | | | 0.054 | -0.029 | 0.045 | 0.054 | 0.132 | 0.17 | 5 | 1 |
| | DeltFallMax | | | -0.101 | -0.239 | -0.151 | -0.101 | -0.024 | 0.005 | 10 | 5 |
| | DeltFallMin | | | -0.098 | -0.332 | -0.144 | -0.098 | -0.042 | 0.17 | 144 | 66 |
| | DeltPrecip | | | 0.059 | -0.106 | 0.003 | 0.059 | 0.069 | 0.182 | 11 | 6 |
| | DeltSpringMax | -0.131 | 0.010 | -0.146 | -0.408 | -0.193 | -0.146 | -0.063 | 0.117 | 168 | 111 |
| | heatload | | | -0.05 | -0.461 | -0.122 | -0.05 | -0.015 | 0.219 | 107 | 34 |
| | organic | | | -0.042 | -0.465 | -0.136 | -0.042 | 0.037 | 0.129 | 18 | 6 |
| | packm | | | 0.06 | -0.328 | 0.025 | 0.06 | 0.086 | 0.24 | 35 | 13 |
| | SixMonthPrecip | | | -0.031 | -0.238 | -0.085 | -0.031 | 0.021 | 0.165 | 75 | 14 |
| | Slope | | | 0.008 | -0.461 | -0.026 | 0.008 | 0.059 | 1.02 | 64 | 17 |
| | SoilpH | | | -0.049 | -0.305 | -0.081 | -0.049 | -0.006 | 0.112 | 68 | 20 |
| | Ydays | -0.004 | 0.042 | -0.004 | -0.024 | -0.006 | -0.004 | -0.001 | 0.008 | 347 | 130 |
| | Year | | | 0.017 | -0.108 | -0.004 | 0.017 | 0.039 | 0.104 | 106 | 25 |

| Species | Variable | Original model coefficient | Original model p | Median | Q 0% | Q 25% | Q 50% | Q 75% | Q 100% | Model Counts | P < 0.05 counts |
|---|---|---|---|---|---|---|---|---|---|---|---|
| *Sphaeralcea parvifolia* | AnnMinTemp | | | -0.006 | -0.141 | -0.041 | -0.006 | 0.015 | 0.1 | 49 | 11 |
| | Aspect | | | -0.057 | -0.133 | -0.081 | -0.057 | -0.012 | 0.053 | 24 | 10 |
| | AWC | | | -0.045 | -0.152 | -0.083 | -0.045 | 0.037 | 0.096 | 60 | 30 |
| | clay | | | 0.07 | -0.076 | 0.032 | 0.07 | 0.103 | 0.263 | 67 | 33 |
| | DeltFallMax | | | 0.034 | -0.062 | 0.009 | 0.034 | 0.069 | 0.095 | 24 | 9 |
| | DeltFallMin | | | -0.083 | -0.211 | -0.118 | -0.083 | -0.059 | 0.025 | 47 | 32 |
| | DeltPrecip | | | 0.085 | -0.023 | 0.061 | 0.085 | 0.116 | 0.166 | 51 | 36 |
| | DeltSpringMax | | | -0.056 | -0.136 | -0.097 | -0.056 | -0.005 | 0.097 | 25 | 12 |
| | heatload | | | -0.072 | -0.158 | -0.118 | -0.072 | -0.007 | 0.04 | 33 | 14 |
| | organic | -0.027 | 0.293 | -0.089 | -0.3 | -0.138 | -0.089 | -0.041 | 0.088 | 161 | 92 |
| | packm | | | 0.094 | -0.034 | 0.027 | 0.094 | 0.142 | 0.225 | 19 | 10 |
| | SixMonthPrecip | | | -0.03 | -0.149 | -0.06 | -0.03 | 0.001 | 0.219 | 66 | 22 |
| | Slope | | | 0.061 | -0.006 | 0.042 | 0.061 | 0.082 | 0.156 | 42 | 21 |
| | SoilpH | 0.073 | 0.015 | 0.076 | -0.242 | 0.041 | 0.076 | 0.116 | 0.321 | 384 | 193 |

| Species | Variable | Original model coefficient | Original model p | Median | Q 0% | Q 25% | Q 50% | Q 75% | Q 100% | Model Counts | P < 0.05 counts |
|---|---|---|---|---|---|---|---|---|---|---|---|
| | Ydays | -0.001 | 0.583 | -0.002 | -0.008 | -0.003 | -0.002 | -0.001 | 0.005 | 154 | 45 |
| | Year | | | -0.027 | -0.058 | -0.036 | -0.027 | -0.017 | 0.005 | 39 | 23 |
| *Sporobolus airoides* | AnnMinTemp | | | 0.061 | -0.08 | 0.042 | 0.061 | 0.112 | 0.167 | 17 | 5 |
| | Aspect | | | -0.062 | -0.175 | -0.099 | -0.062 | -0.042 | 0.102 | 30 | 14 |
| | AWC | | | -0.096 | -0.175 | -0.138 | -0.096 | -0.066 | 0.074 | 35 | 16 |
| | clay | | | -0.009 | -0.177 | -0.093 | -0.009 | 0.072 | 0.127 | 18 | 8 |
| | DeltFallMax | 0.102 | <0.001 | 0.122 | -0.048 | 0.084 | 0.122 | 0.16 | 0.246 | 103 | 67 |
| | DeltFallMin | | | 0.098 | -0.155 | 0.062 | 0.098 | 0.138 | 0.217 | 30 | 21 |
| | DeltPrecip | | | 0.074 | -0.197 | 0.033 | 0.074 | 0.102 | 0.353 | 71 | 22 |
| | DeltSpringMax | | | 0.103 | -0.045 | 0.076 | 0.103 | 0.156 | 0.296 | 105 | 55 |
| | heatload | 0.069 | 0.009 | 0.086 | -0.206 | 0.058 | 0.086 | 0.126 | 0.287 | 172 | 81 |
| | organic | | | 0.031 | -0.146 | -0.076 | 0.031 | 0.093 | 0.189 | 21 | 5 |
| | packm | 0.038 | 0.150 | 0.067 | -0.088 | 0.044 | 0.067 | 0.112 | 0.3 | 78 | 23 |
| | SixMonthPrecip | | | 0.053 | -0.226 | 0.004 | 0.053 | 0.102 | 0.344 | 25 | 7 |
| | Slope | | | 0.025 | -0.202 | -0.022 | 0.025 | 0.089 | 0.249 | 34 | 7 |

| Species | Variable | Original model coefficient | Original model p | Median | Q 0% | Q 25% | Q 50% | Q 75% | Q 100% | Model Counts | P < 0.05 counts |
|---|---|---|---|---|---|---|---|---|---|---|---|
| | SoilpH | | | -0.059 | -0.318 | -0.112 | -0.059 | -0.013 | 0.129 | 42 | 17 |
| | Ydays | | | 0.003 | -0.002 | 0.002 | 0.003 | 0.004 | 0.005 | 63 | 43 |
| | Year | | | -0.009 | -0.073 | -0.019 | -0.009 | -0.004 | 0.047 | 38 | 11 |
| *Sporobolus contractus* | AnnMinTemp | | | 0.072 | -0.161 | 0.052 | 0.072 | 0.089 | 0.158 | 127 | 85 |
| | Aspect | | | 0.025 | -0.083 | -0.004 | 0.025 | 0.049 | 0.103 | 17 | 5 |
| | AWC | | | 0.063 | -0.02 | 0.04 | 0.063 | 0.083 | 0.166 | 61 | 32 |
| | clay | | | 0.019 | -0.156 | -0.044 | 0.019 | 0.05 | 0.128 | 8 | 2 |
| | DeltFallMax | -0.021 | 0.346 | -0.051 | -0.162 | -0.077 | -0.051 | -0.026 | 0.158 | 150 | 69 |
| | DeltFallMin | | | -0.04 | -0.16 | -0.066 | -0.04 | -0.025 | 0.034 | 15 | 4 |
| | DeltPrecip | | | -0.038 | -0.093 | -0.061 | -0.038 | 0 | 0.063 | 41 | 7 |
| | DeltSpringMax | | | 0.057 | -0.077 | 0.033 | 0.057 | 0.074 | 0.118 | 15 | 8 |
| | heatload | 0.047 | 0.032 | 0.045 | -0.129 | 0.024 | 0.045 | 0.078 | 0.361 | 192 | 79 |
| | organic | -0.079 | 0.001 | -0.092 | -0.39 | -0.107 | -0.092 | -0.074 | 0.093 | 301 | 234 |
| | packm | | | -0.046 | -0.176 | -0.066 | -0.046 | -0.025 | 0.182 | 119 | 46 |

| Species | Variable | Original model coefficient | Original model p | Median | Q 0% | Q 25% | Q 50% | Q 75% | Q 100% | Model Counts | P < 0.05 counts |
|---|---|---|---|---|---|---|---|---|---|---|---|
| | SixMonthPrecip | | | 0.002 | -0.111 | -0.044 | 0.002 | 0.071 | 0.275 | 19 | 10 |
| | Slope | | | 0.021 | -0.004 | 0.011 | 0.021 | 0.066 | 0.115 | 6 | 2 |
| | SoilpH | | | -0.003 | -0.177 | -0.022 | -0.003 | 0.015 | 0.12 | 18 | 6 |
| | Ydays | | | 0.002 | -0.005 | -0.001 | 0.002 | 0.003 | 0.006 | 18 | 12 |
| | Year | | | 0.007 | -0.013 | 0.002 | 0.007 | 0.017 | 0.039 | 10 | 2 |
| *Sporobolus cryptandrus* | AnnMinTemp | | | -0.117 | -0.263 | -0.178 | -0.117 | -0.064 | -0.018 | 32 | 19 |
| | Aspect | | | 0.13 | -0.144 | 0.105 | 0.13 | 0.184 | 0.383 | 28 | 23 |
| | AWC | | | -0.128 | -0.333 | -0.153 | -0.128 | -0.084 | 0.05 | 55 | 34 |
| | clay | | | -0.08 | -0.308 | -0.153 | -0.08 | -0.043 | 0.45 | 61 | 19 |
| | DeltFallMax | 0.118 | 0.003 | 0.175 | -0.022 | 0.127 | 0.175 | 0.226 | 0.416 | 112 | 76 |
| | DeltFallMin | | | -0.11 | -0.253 | -0.138 | -0.11 | -0.068 | 0.057 | 38 | 19 |
| | DeltPrecip | | | 0.118 | -0.109 | 0.022 | 0.118 | 0.196 | 0.283 | 31 | 16 |
| | DeltSpringMax | | | 0.054 | -0.292 | -0.083 | 0.054 | 0.129 | 0.441 | 24 | 9 |
| | heatload | | | 0.083 | -0.168 | 0.011 | 0.083 | 0.18 | 0.421 | 29 | 5 |

| Species | Variable | Original model coefficient | Original model p | Median | Q 0% | Q 25% | Q 50% | Q 75% | Q 100% | Model Counts | P < 0.05 counts |
|---|---|---|---|---|---|---|---|---|---|---|---|
| | organic | | | -0.226 | -0.773 | -0.38 | -0.226 | -0.059 | 0.107 | 105 | 50 |
| | packm | | | -0.082 | -0.237 | -0.131 | -0.082 | -0.018 | 0.238 | 23 | 5 |
| | SixMonthPrecip | | | -0.067 | -0.146 | -0.105 | -0.067 | -0.03 | 0.114 | 19 | 6 |
| | Slope | | | 0.034 | -0.091 | -0.002 | 0.034 | 0.118 | 0.209 | 14 | 3 |
| | SoilpH | | | 0.003 | -0.538 | -0.097 | 0.003 | 0.102 | 0.438 | 45 | 16 |
| | Ydays | | | -0.003 | -0.009 | -0.005 | -0.003 | -0.002 | -0.001 | 38 | 26 |
| | Year | -0.022 | 0.084 | -0.066 | -0.211 | -0.106 | -0.066 | -0.035 | 0.024 | 132 | 91 |
| *Sporobolus flexuosus* | AnnMinTemp | | | 0.049 | -0.294 | -0.018 | 0.049 | 0.072 | 0.226 | 86 | 38 |
| | Aspect | | | 0.065 | -0.109 | 0.001 | 0.065 | 0.134 | 0.233 | 87 | 53 |
| | AWC | | | -0.005 | -0.149 | -0.034 | -0.005 | 0.026 | 0.037 | 11 | 3 |
| | clay | | | -0.044 | -0.583 | -0.077 | -0.044 | -0.015 | 0.035 | 27 | 6 |
| | DeltFallMax | | | 0.064 | -0.197 | 0.002 | 0.064 | 0.112 | 0.258 | 53 | 28 |
| | DeltFallMin | | | -0.013 | -0.26 | -0.058 | -0.013 | 0.006 | 0.047 | 13 | 4 |
| | DeltPrecip | | | 0.044 | -0.163 | 0.025 | 0.044 | 0.054 | 0.138 | 74 | 44 |

| Species | Variable | Original model coefficient | Original model p | Median | Q 0% | Q 25% | Q 50% | Q 75% | Q 100% | Model Counts | P < 0.05 counts |
|---|---|---|---|---|---|---|---|---|---|---|---|
| | DeltSpringMax | | | 0.036 | -0.367 | 0.002 | 0.036 | 0.079 | 0.199 | 64 | 34 |
| | heatload | 0.208 | <0.001 | 0.281 | -0.16 | 0.112 | 0.281 | 0.337 | 0.54 | 333 | 303 |
| | organic | | | -0.098 | -0.509 | -0.166 | -0.098 | -0.052 | 0.02 | 16 | 7 |
| | packm | | | 0.023 | -0.126 | -0.018 | 0.023 | 0.081 | 0.371 | 59 | 21 |
| | SixMonthPrecip | | | -0.071 | -0.345 | -0.153 | -0.071 | 0.022 | 0.113 | 67 | 34 |
| | Slope | | | -0.118 | -0.678 | -0.207 | -0.118 | -0.068 | -0.003 | 65 | 46 |
| | SoilpH | | | -0.008 | -0.074 | -0.012 | -0.008 | 0.007 | 0.027 | 9 | 0 |
| | Ydays | | | 0.001 | -0.006 | -0.001 | 0.001 | 0.002 | 0.003 | 14 | 5 |
| | Year | | | -0.059 | -0.117 | -0.092 | -0.059 | -0.037 | 0.011 | 25 | 21 |
| *Stanleya pinnata* | AnnMinTemp | | | 0.126 | -0.428 | -0.002 | 0.126 | 0.172 | 0.393 | 24 | 8 |
| | Aspect | | | -0.043 | -0.287 | -0.091 | -0.043 | 0.02 | 0.261 | 59 | 13 |
| | AWC | | | -0.066 | -0.275 | -0.128 | -0.066 | -0.005 | 0.53 | 172 | 56 |
| | clay | | | -0.06 | -0.485 | -0.165 | -0.06 | 0.012 | 0.361 | 155 | 45 |
| | DeltFallMax | | | 0.123 | -0.021 | 0.033 | 0.123 | 0.159 | 0.297 | 8 | 3 |
| | DeltFallMin | | | -0.004 | -0.405 | -0.062 | -0.004 | 0.146 | 0.464 | 44 | 15 |

| Species | Variable | Original model coefficient | Original model p | Median | Q 0% | Q 25% | Q 50% | Q 75% | Q 100% | Model Counts | P < 0.05 counts |
|---|---|---|---|---|---|---|---|---|---|---|---|
| | DeltPrecip | | | 0.276 | -0.102 | 0.146 | 0.276 | 0.339 | 1.141 | 125 | 89 |
| | DeltSpringMax | | | 0.022 | -0.22 | -0.064 | 0.022 | 0.056 | 0.106 | 17 | 4 |
| | heatload | 0.296 | 0.002 | 0.18 | -0.219 | 0.09 | 0.18 | 0.25 | 0.46 | 40 | 22 |
| | organic | | | -0.231 | -0.859 | -0.325 | -0.231 | -0.087 | 0.228 | 245 | 139 |
| | packm | | | -0.003 | -0.258 | -0.024 | -0.003 | 0.049 | 0.315 | 12 | 1 |
| | SixMonthPrecip | | | -0.01 | -0.14 | -0.061 | -0.01 | 0.014 | 0.099 | 6 | 1 |
| | Slope | | | -0.187 | -0.571 | -0.332 | -0.187 | -0.081 | 0.303 | 93 | 56 |
| | SoilpH | | | -0.023 | -0.319 | -0.079 | -0.023 | 0.098 | 0.376 | 27 | 6 |
| | Ydays | | | 0.005 | -0.01 | 0.003 | 0.005 | 0.007 | 0.014 | 147 | 37 |
| | Year | | | -0.01 | -0.05 | -0.021 | -0.01 | -0.001 | 0.19 | 68 | 10 |

**Table S7.** Results for the phylogenetic generalized least squares model testing the effect of climate, environment, and space on seed mass variation. Data is aggregated at the species level using the 10 collection dataset (N = 119 species aggregated across 5100 populations). Seed mass coefficient of variation is calculated as $\text{CVunbiased}=(1+14\times N)\times\text{CV}$. Covariates were selected using random forest and are centered and scaled.

| *Covariate* | *Estimate* | *SE* | *t* | *p* | *lower 95% CI* | *upper 95% CI* |
|---|---|---|---|---|---|---|
| Maximum latitude | 0.091 | 0.037 | | <0.001 | 0.018 | 0.164 |
| Range of collecting days | 0.055 | 0.041 | | 0.180 | -0.025 | 0.134 |
| Range of annual minimum temperature | 0.068 | 0.041 | | 0.101 | -0.013 | 0.148 |

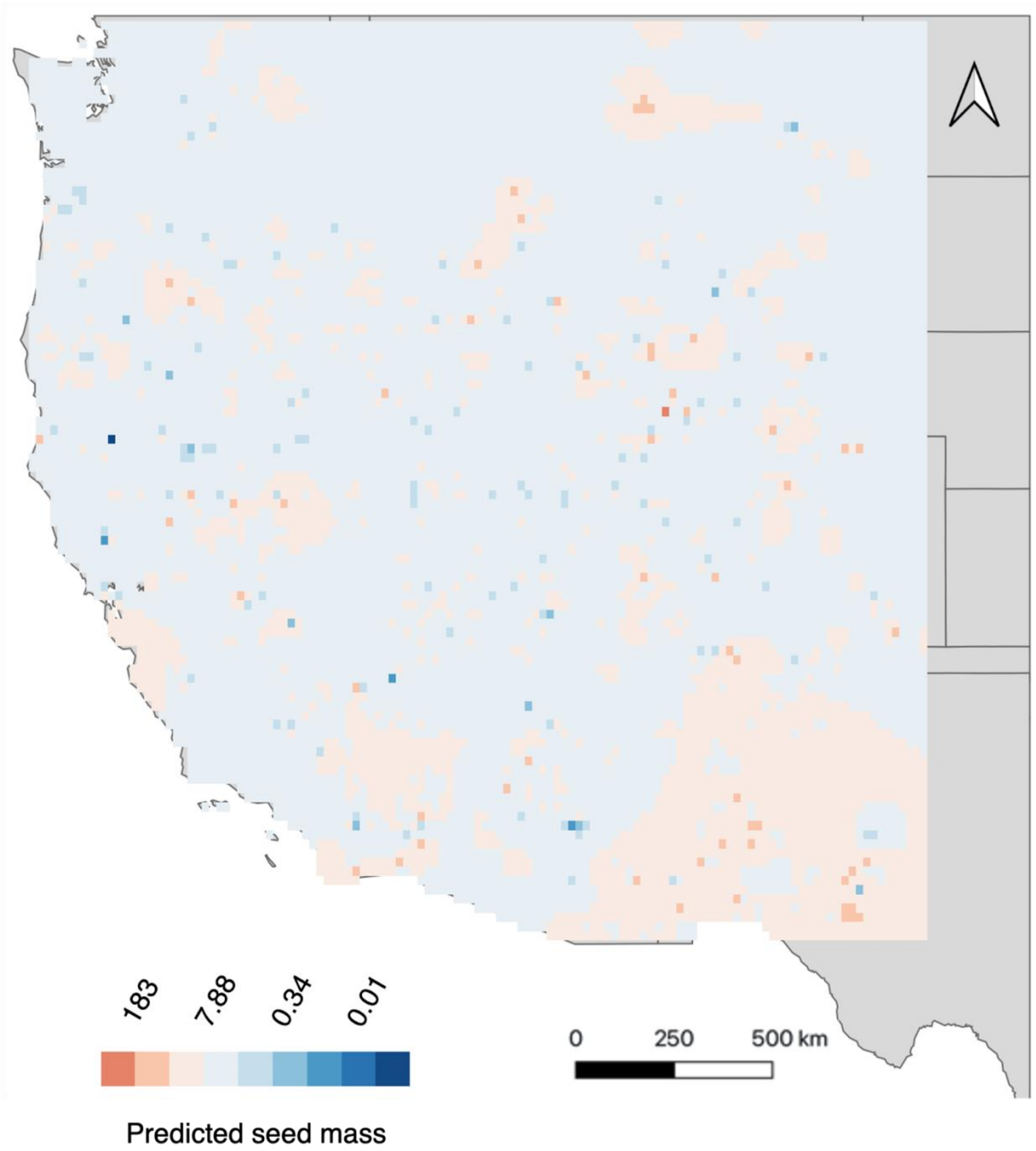

**Figure S1.** Results of inverse distance weighted interpolation predicting seed mass across the western United States. Seed mass was log transformed from 13,004 seed lots from Seeds of Success.

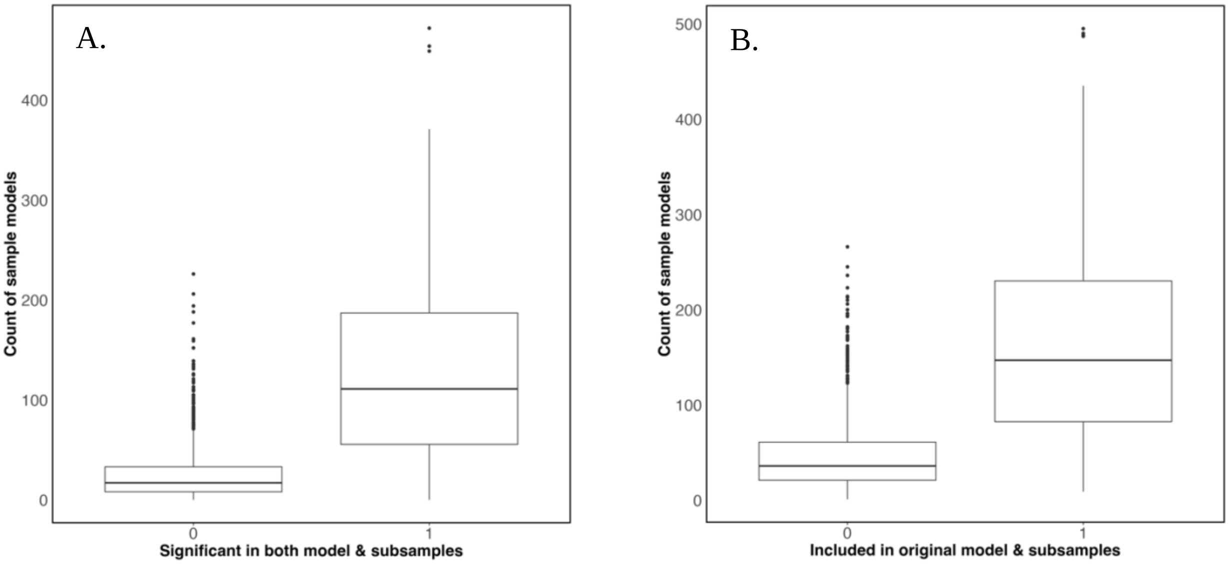

**Figure S2.** Boxplots representing summary of subsampling. We randomly sampled 20 populations 500 times for each species that had been collected from at least 25 populations. Following the random forest pipeline to determine covariates we then ran spatial error GLMs. Boxplots represent how many times the original model coefficients were included in the subsampled model (A) and the number of times the coefficient was statistically significant ($p < 0.05$) in both the original and subsampled models, 1 indicates a match and 0 indicates a non-match between subsamples and original model.
